# FHIR as a Unifying Format for Genomic Research Data Tracking, Aggregation, and Integration

**DOI:** 10.64898/2025.12.22.695544

**Authors:** Nasim Sanati, Brian Walsh, Parker Gray, Lauren Hagen, Robert Carroll, Allison Heath, Boris Aguilar, Amanda Charbonneau, Kyle Ellrott

## Abstract

The increasing complexity of genomic research demands standardized data sharing and integration. The Fast Healthcare Interoperability Resources (FHIR) specification has become a well-established standard for exchanging data among health data systems. While designed primarily for clinical and patient data in health care environments, it also has applicability to represent genomic research data and offers a path for aggregating and integrating extremely rich datasets that have traditionally remained siloed and disparate. To study this potential, we developed FHIR Aggregator, an integration of seven major biomedical repositories, including the Genomic Data Commons, GTEx, HTAN, and DepMap, that covered 142334 patients, 819251 specimens, 1096491 observations and 711166 documents. We explore the various ways the FHIR standard can be applied to structure genomic research data and enable new possibilities. We demonstrate how FHIR can be used, where it succeeds or falls short, and which concepts must be extended to better support large-scale clinical and genomics research projects.

## Background

The Fast Healthcare Interoperability Resources (FHIR) standard has become a foundational model for structuring and exchanging clinical and patient data across electronic medical record (EMR) systems, with broad international adoption in over 52 countries[1]. Its modular and extensible design has enabled broad interoperability across healthcare institutions, supporting cohort analysis, clinical decision-making, and patient-centric data exchange. In the United States, the federal mandate to support EHR interoperability through the design and implementation of the US Core Data for Interoperability (USCDI) has demonstrated the value of the base FHIR model and the benefit of establishing common semantics[2]. Beyond data exchange, the inherently graph-structured nature of FHIR (**Figure 1**), in which resources such as patients, conditions, and observations reference one another, provides a powerful foundation for computational analysis. Recent work has shown that this structure can be leveraged for descriptive analytics by traversing these relationships to support patient-centered and cohort-based queries, positioning FHIR as a viable platform for both interoperability and analytical insight. Many groups have begun to utilize FHIR for integrative clinical analysis [3–6], utilizing its data model to merge datasets between multiple hospitals. While FHIR has found utilization for supporting aggregation studies in the field of Health Informatics, there is still the question of if it can provide similar benefits for genomic research[7].

**Figure 1.**
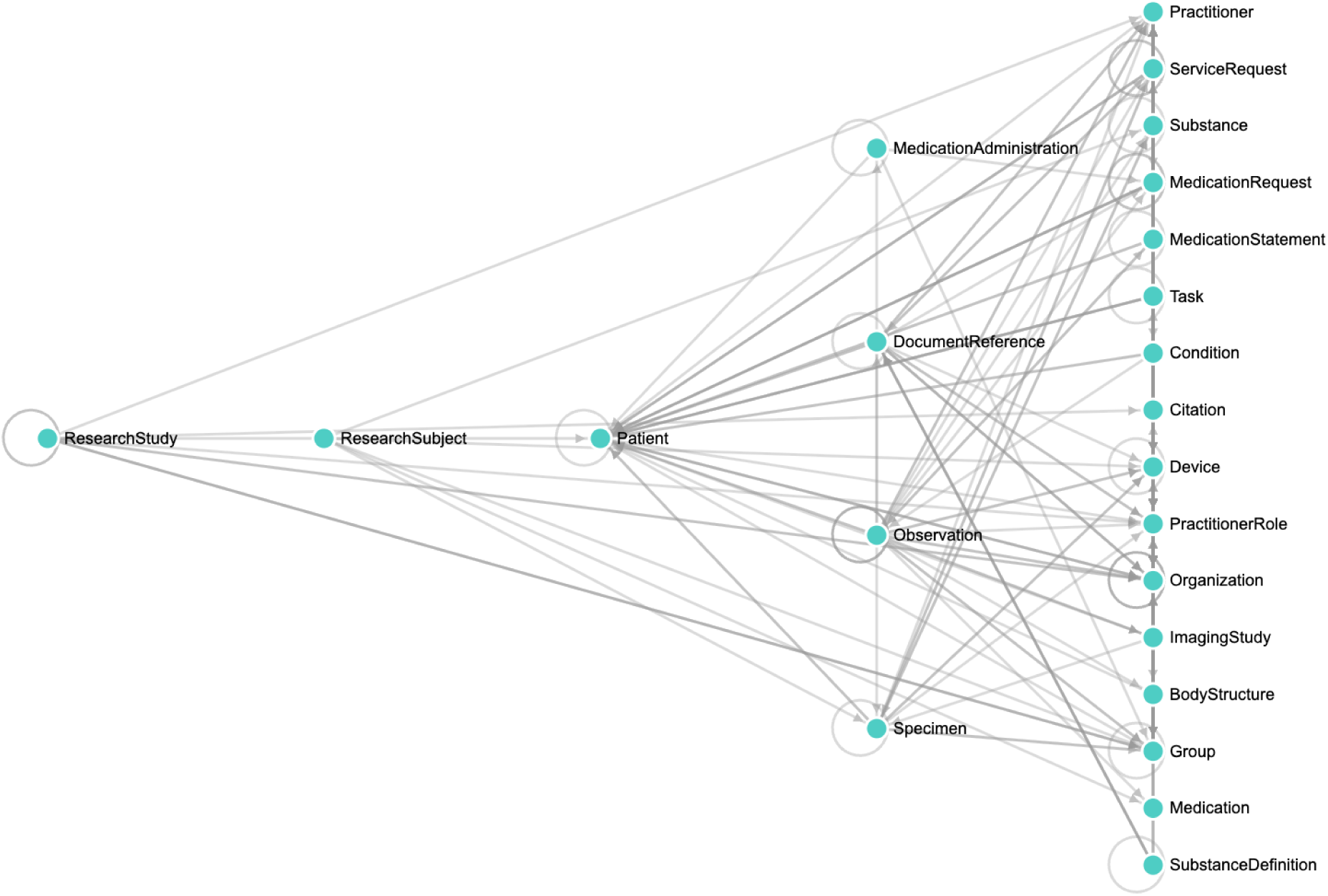
Schema structure implemented in the FHIR Aggregator framework. This diagram reflects the actual FHIR resource relationships used across the integrated dataset. Key entities such as Patient, Specimen, Observation, and DocumentReference are connected through schema-defined references, illustrating the graph-oriented structure of FHIR and its alignment with study-level organization.

Despite their importance for precision medicine, therapeutic decision-making, and biomedical discovery, genomic datasets consented for research are seldom represented using the structured, interoperable formats widely adopted in healthcare systems. This gap significantly constrains their clinical and analytical utility. For example, the NCI’s Genomic Data Commons[8] is structured as a graph-based model with field properties defined by the Cancer Data Standards Registry and Repository (caDSR) metadata system[9], whereas datasets from GTEx[10] and the 1000 Genomes Project[11], are provided primarily as tab-separated value (TSV) files with custom field names. Integrating the meta-data of patients and samples across these disparate cohorts requires labor and specialized knowledge to build mappings of terminology and find a unified data model. More recently, the eMERGE Network, Health Level Seven (HL7) Clinical Genomics Work Group, and Sync for Genes initiative have demonstrated the feasibility of representing sequencing results, variants, and phenotype data using FHIR in real-world and pilot settings. These efforts illustrate that while FHIR shows potential for structured genomic exchange, its broader adoption and integration into large-scale research infrastructures remains unexplored. Further, a systematic integration of genomics data into FHIR, especially across diverse consortium-level repositories, has not been developed. This represents a critical gap. Historically, research data integration has relied on common data models such as OMOP-CDM[12] and i2b2[13], as well as cloud-native platforms like Gen3[14], to manage clinical and omics data. While these frameworks have advanced interoperability within research domains, they are typically disconnected from clinical FHIR infrastructure, which limits traceability between patient care and research findings.

Several NIH funded efforts have begun mapping existing datasets into FHIR. The Gabriella Miller Kids First Data Resource Center[15] has developed a FHIR based view[16] on their dataset, which includes 36 studies and 38 thousand participants. The AnVIL Clinical Resource[17] is developing a harmonized version of the catalogue of almost 300 NHGRI datasets compatible with FHIR. The NIH INvestigation of Co-occurring conditions across the Lifespan to Understand Down syndromE (INCLUDE) Data coordinating center[18] and ImmPort[19] have also claimed to support FHIR. Finally, the NIH’s dbGap has made their data available via FHIR[20].

To extend these efforts, and discover the possible applications of integrative analysis using a common FHIR based framework for genomic research, we developed FHIR Aggregator. This dataset, based on the work of the Cancer Data Aggregator[21] integrates seven major biomedical repositories: the Cancer Data Aggregator (CDA), Genomic Data Commons (GDC), Genotype-Tissue Expression Portal (GTEx), Human Tumor Atlas Network (HTAN), the International Cancer Genome Consortium (ICGC) Cellosaurus cell line database, and the 1000 Genomes Project. Each source presents unique structural and semantic challenges, which we address through configurable transformation pipelines, terminology harmonization using systems such as LOINC, SNOMED CT, and NCIT caDSR, standardized chemical descriptors including SMILES via ChEMBL, and reusable schema graph definitions. These definitions support query path detection and adjustment, enabling analyses such as survival outcomes and cross-study mutation query workflows that are essential for scalable genomics research.

FHIR Aggregator demonstrates how modular transformation pipelines and schema-driven publishing can connect research and clinical domains. Moving heterogeneous data resources into the common FHIR schema allows the data model to support integration across genomic, clinical, and pharmacological datasets, enables study-scoped graph queries, and enforces HL7-compliant validation. This approach provides a foundation for more connected data points required for biomedical analyses that can improve care delivery, enhance research outcomes, and accelerate data-driven discovery.

## Results

We mapped seven consortium-scale, multimodal genomics datasets, including CDA, GDC, HTAN, Cellosaurus, ICGC, and 1000 Genomes, to the HL7 FHIR standard and integrated them using an open-source framework called FHIR Aggregator[22]. Each dataset was transformed independently into FHIR-compliant resources and loaded into the Google Cloud Healthcare API to enable graph-based exploration across studies. The resulting dataset includes over 1.4 million HL7 FHIR R4 resources, covering 142,334 patients, 819,251 biospecimens, 1,096,491 structured observations, and 711,166 document references. These were organized into a schema-driven graph model, with resources such as Patient, Specimen, Observation, and ResearchStudy consistently linked through standard references.

### Benefits of the FHIR schema

#### FHIR Enables Research

Integrative cancer analysis requires identifying possible samples and obtaining the files that are associated with them. This process is often time-consuming and labor-intensive, as researchers must manually search through multiple data resources and continually assess whether samples have similar enough data to be combined for analysis.

Our FHIR graph connects entities such as patients, specimens, variants, and associated files through resolvable references. This graph-oriented schema provides a foundation for cross resource cohort building, navigating across datasets and enabling more structured, analysis-ready integration of research-grade genomics data suitable for complex, multi-dimensional analyses. There are two basic ways that researchers can engage with our FHIR graph to accelerate their own research: as a data discovery tool or as a direct source for analysis.

As a data discovery tool, FHIR can be used to query across disparate datasets to build a cohort of samples that would normally require search at multiple repositories that use idiosyncratic data models. This tasks the researcher with not only finding each repository, but with learning enough about each model to run an effective search. In our FHIR implementation, the metadata from each of our resources is mapped to a single FHIR model, which results in concept level harmonization, i.e. all data relating to ‘condition’ has been combined regardless of where it was found in its original model. While we have not taken the additional step of semantically harmonizing within each concept, (i.e. female = f = F), this does not reduce the utility of search with FHIR. For instance, researchers can use wild card characters to construct flexible searches that work around this variation; a query like “birthSex = f*” will return any value in that field beginning with “f”. These inconsistencies were present in the source data but were less apparent when distributed across separate repositories. The harmonized structure provided by FHIR makes them more visible, enabling researchers to identify relevant data variation and construct queries that align more effectively with their research objectives.

Once a researcher has built a cohort, they can also use FHIR to access the underlying data for each subject/sample or to learn what data use agreement is necessary to gain access. The Global Alliance for Genomics and Health (GA4GH)[23] Data Repository Service (DRS) provides a standardized way to access data across repositories regardless of their underlying storage and management system. It provides an abstraction layer hiding the complexities of storage methods and enables users to interact with actual data content through DRS IDs vs complex storage paths. In FHIR Aggregator, DRS URIs (ex. drs://dg.4ABC:) are embedded in FHIR’s ‘DocumentReference.content.attachment.url’ fields to facilitate federated file access. This allows researchers to resolve file locations across distributed NIH platforms without duplicating storage, preserving FHIR compliance while aligning with FAIR data practices.

It is also possible to use our FHIR graph as a direct source for analysis by focusing on questions that can be answered using only metadata. For example, a researcher could use ‘US-core-patient’ fields to collect sex, race and ethnicity values to plot against ‘condition’ and ‘observation’ information such as type of disease, when and how diagnoses were made, and follow-up data to assess racial and gender disparities in cancer outcomes. Basic patient-level pharmacogenomic exploration could also be done using only FHIR data by linking observed mutations to administered medications. Using Observation resources for genomic variants and MedicationAdministration for recorded treatments, researchers can identify drugs taken by patients with specific mutations at defined timepoints. While no retrospective study is conclusive, this kind of analysis could be used as preliminary data for securing funding of treatment patterns in relation to molecular profiles or exploratory precision oncology studies.

To explore the utility of FHIR for analysis we performed survival analysis with FHIR Aggregator data (see Supplementary Analysis and Notebooks). We leveraged standardized Logical Observation Identifiers Names and Codes (LOINC) [24] coded observations of disease status and days to death to assess survival across different cancer studies including TCGA and CPTAC[25]. One simple query was used to retrieve the necessary metadata, which is converted into pandas dataframes for ease of use in analytical workflows. These queries demonstrate FHIR’s ability to search diverse biomedical data types; from genomic sequencing data to clinical treatment information, while preserving study-specific context and allowing aggregation across multiple studies..

#### Ease of Computational Setup

FHIR is traditionally seen as an enterprise service, but it is increasingly possible to easily ‘roll your own’ server and deploy it on a standard laptop. FHIR Aggregator provides open source tooling to load biomedical datasets[26], previously aggregated or prepared by the broader FHIR-Aggregator ecosystem, into a locally hosted HAPI FHIR server. This facilitates experimentation, querying, and local analysis of FHIR-formatted clinical and research datasets without relying on cloud infrastructure or enterprise level service agreements. This tool provides the connective tissue required for local deployment for analysis, validation, or development, allowing users to spin up a HAPI FHIR environment and populate it with curated FHIR resources (ex. NDJSON data). This is especially useful for research teams who require offline access, environment isolation, or reproducibility in development workflows. The overall approach supports a fully open-source FHIR pipeline: datasets are retrieved via the FHIR-Aggregator platform, then loaded into a self-managed FHIR server for subsequent queries and validation. This setup empowers researchers to explore and validate standardized FHIR resource content in a controlled, local environment, bridging cloud-based aggregation with offline usability.

There is also increasing support for cloud-based deployments that can reduce the maintenance and operational costs for service costs. The major cloud providers, Amazon Web Services, Microsoft Azure and Google Cloud Platform also provide capabilities for managed FHIR servers. These systems can be deployed as a service, so users are charged by usage rather than having to pay for a fixed virtual machine resource; these offerings also allow the capability to scale up as demands increase.

### Remaining Challenges with using FHIR for research data

While FHIR provides a critical foundation for clinical data standardization and interoperability, its application to genomics research revealed recurring design patterns and structural challenges. This reflects how FHIR’s current schema favors minimal, clinically actionable slices of molecular information, while research use cases demand richer, more integrated representations that must exhibit full scalability beyond limited gene set profile use-cases for web-based applications. Rather than replacing FHIR, our objective is to extend its utility by identifying structural gaps and proposing schema-level adaptations that preserve its core strengths while supporting research-specific needs.

#### Over Reliance on Observation Structure

A major limitation of using FHIR alone for genomics research is its lack of dedicated structures for core molecular entities such as alleles, transcripts, proteins, or pathways. For example, a variant allele and its effect, including specific consequence or predicted protein impact, must be represented using the generic Observation resource. This leads to a pattern we refer to as Observation overload, where structured molecular data is either densely packed into a single Observation using the component field or heavily fragmented across many Observations. In the first case, attributes become difficult to search or filter across studies; in the second, their biological grouping, for example an allele and its effect, are lost due to lack of explicit linkage. This weakens semantic clarity, reduces query precision, and complicates integrated analysis. The other technical effect of Observation overload is that this large collection of heterogenous records are stored in a table that is traditionally designed to store a sparse number of patient level phenotypic observations. Strategies for large data storage, such as using columnar stores[27] to efficiently pack and index information won’t be deployed. While the FHIR model offers partial workarounds, it doesn’t fully address the absence of granular relational models required for robust molecular representations that are essential for genomic research and discovery. As a result, data consumers face significant barriers when attempting to perform analysis-ready extractions from FHIR-native representations of complex omics datasets

#### Variant Representation

In clinical healthcare settings, genetic testing is typically limited to a small subset of well-characterized variants, primarily within the 1-2% of the genome that encodes proteins and current data standards reflect the constraints of clinical priorities[28]. Variants are selected based on known diagnostic or therapeutic relevance and are often identified through targeted panels or whole exome sequencing. While this approach is efficient for clinical workflows, it overlooks the broader genomic landscape, including rare, regulatory, and population-specific variants that may contribute to disease risk, heterogeneity, progression, and therapeutic resistance.

Although FHIR includes a MolecularSequence resource, it is limited to raw sequence data and explicitly excludes variants, annotations, and genotypes[29]. Genomic research requires representation of the full spectrum of variation and biological context from whole genome sequencing and multimodal assay. In FHIR, these must instead be modeled using the generic Observation resource, leading to structural complexity and query inefficiency. The vcf2fhir utility[30] illustrates this challenge. Converting VCF data to FHIR using this tool generates four Observation types per variant: Variant, Region Studied, Sequence Phase Relationship, and Diagnostic Implication. Each variant Observation contains up to 15 nested components such as allelic state, reference allele, alternate allele, genomic coordinates, copy number, and clinical annotations, all encoded with system, code, and display fields (Supplementary Table 1). This creates substantial per-record overhead that scales across tens of thousands of variants per patient in whole genome sequencing datasets. These limitations reflect a fundamental structural gap in FHIR for representing the complex molecular relationships essential to genomic research. Although efforts such as the FHIR Genomics Operations extension[31] provide valuable functionality, they do not resolve the underlying constraints of the FHIR data model for representing variant-level relationships.

#### Representing Transcriptomic Data

Similar limitations arise when representing transcriptomics data in FHIR. Auer et al. [32] proposed adapting the FHIR Genomics extension to encode gene expression profiles from microarray assays using MolecularSequence, Observation, and DiagnosticReport resources. While this demonstrates the feasibility of transcriptomic data sharing within the FHIR framework, the approach is structurally misaligned. MolecularSequence is limited to raw sequence data and explicitly excludes analytical results, making it fundamentally unsuitable for the dense, high-dimensional matrices produced by transcriptomic assays such as microarrays and RNA-Seq.

The proposed method treats expression values as isolated Observations, rather than capturing per-gene, per-sample data. This is incompatible with tasks such as differential expression, clustering, and predictive modeling which require all gene expression values per sample to be aggregated for analysis. Without structural support for dense, quantitative omics data, FHIR remains insufficient for transcriptomics use cases. This limits interoperability and constrains efforts to integrate high-resolution assays into clinical and research workflows, ultimately hindering systems-level analysis and applications in precision medicine[7,33].

#### scRNA-Seq Resolution and Cell Level schema Design

FHIR’s capacity to support high-resolution, temporally-aware single-cell workflows and lineage tracking without custom extensions or external tooling is limited. Single-cell platforms treat each cell as a distinct entity, annotated with expression profiles, clustering labels, quality metrics, and often trajectory relationships, for example CellA to CellB, which capture cellular evolution over time and are essential for modeling disease progression and differentiation. FHIR lacks a dedicated resource to represent individual cells or their lineage. As a result, expression data must be either flattened into multiple Observation entries or stored externally, for example in HDF5 or TSV files referenced by DocumentReference, however this removes native query access and strains performance at scale.

#### Specimen Hierarchy and Biopsy Lineage

While FHIR can adequately represent individual patient samples, it lacks the highly organized structures required for sub-sample tracking. Biomedical platforms define deeply nested specimen hierarchies. Biopsy, Sample, Portion, Analyte, Aliquot with files typically linked at the most granular level[34]. In practice, researchers are primarily interested in specimens with associated files, especially when comparing multiple modalities, timepoints, or assays.

FHIR uses a single Specimen resource and allows hierarchical relationships through the optional parent field. A specimen without a parent often represents a biopsy and may serve as the root of a specimen tree. However, FHIR does not enforce hierarchy; the structure must be explicitly modeled during transformation. To distinguish temporally distinct biopsies or repeated collections from the same patient, deidentified research metadata such as Days to Collection must be incorporated as secondary identifiers. Without this temporal dimension, lineage reconstruction and downstream analysis are impaired. Moreover, querying whether a specimen has a descendant aliquot that is linked to a file often requires recursive reference traversal or custom search parameters, which can significantly impact query performance. FHIR’s RESTful API is not optimized for such nested graph traversals, making retrieval of file-associated specimens inefficient unless indexed or preprocessed in advance.

#### Patient Identity Within Study Contexts

Research datasets often scope patient identifiers to a specific study (ex. StudyA-Patient01), enabling local de-identification and preventing linkage across datasets. FHIR separates the Patient resource from study participation using ResearchSubject, allowing a patient to be associated with multiple ResearchStudy instances. While this design supports reuse, it increases the risk of re-identification if Patient.identifier values are reused without pseudonymization. To maintain privacy in research settings, patients must be study-scoped or remapped at the time of ingestion.

#### Drug Response and High-Throughput Screens

As FHIR is a patient centered tool, it can be difficult to represent inherently population level research data. Pharmacogenomic platforms model dose-response experiments using structured matrices that capture drug concentration, cell viability, replicates, and compound metadata. FHIR lacks a format for this representation. In our implementation, cell lines are modeled as Patient, drug administrations as MedicationAdministration, and summary response metrics (ex. IC50) as Observation. However, full dose-response curves, replicate tracking, and multidose designs cannot be natively represented, requiring custom encodings of Observations which leads to Observation overload.

#### Competing Schema Standards

As FHIR continues to evolve, version adoption has become uneven across platforms. FHIR R4 remains the widely supported version in production systems, while FHIR R5 introduces incremental improvements in resource structure, linkage semantics, and terminology expressiveness. FHIR R6 is anticipated to be released in late 2026.

To address this, the FHIR Aggregator includes a version-aware submission pipeline that converts internally generated R5 resources into R4[35]. At the time of implementation, platforms such as the Google Cloud Healthcare API only supported R4 and added R5 later in July 2025.

The transformation process implemented in the submission pipeline addresses key differences between the two versions (Table 1). Unsupported fields in R4 are either removed or mapped to custom extensions, and implicit R5 relationships are made explicit using R4 constructs like basedOn, partOf, and context. The pipeline also ensures conformance with R4 value sets, replacing R5-specific codes with the closest allowable alternatives. Resources such as ServiceRequest, DiagnosticReport, and DocumentReference are restructured to preserve clinical intent within the constraints of the R4 model.

**Table 1.** Summary of major FHIR R5 changes and their corresponding FHIR R4 compatibility approaches. This down-conversion strategy allowed developers to model data in R5 while maintaining compatibility with existing R4-based infrastructure. It serves as an example of a practical bridge between current deployment constraints and forward-looking modeling needs, ensuring semantic integrity and interoperability across evolving FHIR versions.

| Domain | R5 Characteristics | R4 Compatibility Approach |
| --- | --- | --- |
| Resource Fields | New elements (ex. DiagnosticReport.media.reference) | Removed or replaced with custom extensions |
| Workflow Modeling | More flexible linkage semantics between resources | Explicit basedOn, partOf, and context references added |
| Data Types | Enriched data types (ex. canonical, markdown) | Re-encoded using R4-supported base types |
| **Extensions** | Native fields in R5 reduce the need for extensions | Semantically equivalent extensions reintroduced in R4 |
| **Terminology** | Updated or trial-use value sets | Mapped to the nearest R4-supported code systems |

## Discussion

Transforming genomics research data into HL7 FHIR presents significant challenges due to fundamental differences in schema structures, terminology systems, and design priorities across data sources. Platforms such as GDC and HTAN reflect divergent modeling strategies, with GDC focusing on case-level structure and HTAN centering around file granularity. Aligning these with FHIR’s resource-based model required custom transformation pipelines that preserve both semantics and study-specific context.

A key lesson from this work is the tension between schema modularity and modeling depth. FHIR offers a robust, reference-driven structure that promotes data consistency and validation. However, it lacks native representations for essential bioinformatics concepts such as molecular variants, alleles, expression patterns, and cell-level lineage. As a result, these concepts must often be represented using the generic Observation resource, leading to what we describe as Observation overload. This occurs when densely packed component fields obscure relational meaning, or when fragmented Observations fail to preserve essential biological groupings. The result is reduced query precision and limited analytical utility unless these representations are carefully constrained and documented.

The schema-level gaps reflect a broader divergence in priorities between clinical and research systems. Clinical infrastructures emphasize completeness, consistency, and alignment with workflows such as billing, diagnosis, and care delivery. In contrast, research systems must accommodate evolving study designs, derived measurements, high-throughput assay outputs, and entity structures specific to genomics and experimental biology. These elements must often be retrieved efficiently to support scalable computational workflows. In this context, FHIR frequently serves only as the surface layer. It provides a standardized entry point but lacks the schema depth required to express high-dimensional models such as inferred phenotypes, biomarker associations, or lineage-based cell transitions. Accurately modeling these molecular and assay-level relationshipships requires domain-specific schemas that require precise semantic structure while retaining performance at scale.

Privacy modeling also differs. Research datasets typically rely on study-scoped patient identifiers and must avoid direct linkage to clinical records to reduce the risk of reidentification. Deidentifying temporal information is equally crucial. Instead of exposing actual dates, research systems often use relative time measures such as days to diagnosis or days to death. These values can be computed from clinical timestamps and then discarded, preserving temporal context while minimizing disclosure risk. FHIR’s structural separation of Patient and ResearchSubject has proven essential in supporting these research-specific requirements.

Deployability is another important factor. Although FHIR is typically associated with large-scale enterprise systems, our framework demonstrates that it can be adapted for lightweight research use. Using containerized components, schema templates, and simplified client interfaces, we show that researchers can deploy and query study-scoped FHIR data on local machines. This lowers technical barriers and enables reproducible, FAIR-aligned workflows without requiring institutional-scale infrastructure.

The transformation process itself is not easily automated. While reusable mappings and modular pipelines provide a strong foundation, schema transformation in genomics remains context-sensitive and often requires manual decisions about resource modeling, terminology mapping, and reference logic. These decisions are informed by biological expertise and study-specific design constraints. Rather than depending on a single universal transformer, effective workflows require a structured approach that separates concerns across high-level schema entities, field-level mappings, and controlled vocabularies. Future developments may explore assisted transformation using inference or pattern recognition, but human oversight remains essential for preserving interpretability and fidelity across heterogeneous datasets.

In practical terms, what worked well was a layered strategy: resource-specific transformation logic stored in modular META folders, GraphDefinition templates that encode study structure, and study-scoped filtering that optimizes both performance and relevance. This approach enabled us to accommodate structural differences across datasets while supporting integrated, analysis-ready querying across resources that were never designed to interoperate.

## Conclusions

The NCPI FHIR Aggregator illustrates how HL7 FHIR, when extended with modular tooling and schema-aware design, can support the integration and exploration of complex biomedical research data. By harmonizing datasets from seven major repositories including clinical records, molecular profiles, pharmacological screens, and cell line metadata. This framework enables standardized analysis workflows that align with study design and research intent.

Through subgraph templates and validated resource structures, the system facilitates consistent cohort extraction, graph traversal, and federated file referencing. These capabilities make it possible to query across previously siloed domains and perform analyses that link genomic variation to clinical outcomes or treatment history. The ability to deploy the entire system locally or in the cloud further enhances accessibility for both exploratory use and production-grade pipelines.

Ongoing development will expand coverage of study-specific subgraphs, improve support for structured multi-omic assays, and introduce schema-driven methods for building machine learning-ready views of the data. While we anticipate greater use of AI-assisted schema transformer agents in our future work, the current architecture remains grounded in modularity, validation, and expert-informed mappings. This ensures that the framework remains flexible and transparent across use cases.

Rather than positioning FHIR as a complete solution for genomics, this work demonstrates how its data model requires extension to effectively bridge research and clinical genomics. The NCPI FHIR Aggregator serves as a proof of concept by supporting consistent, schema-driven integration across clinical records, molecular profiles, medication administrations, and cell line metadata, that illustrates both what is feasible today and what will require additional extensions for true multi-omic interoperability. In doing so, FHIR Aggregator provides a foundation that has the potential to support a more connected research data ecosystem and facilitate new possibilities for translational research, multi-institutional collaboration, and precision medicine discovery.

## Supporting information

Supplementary Analysis and Notebooks

## Access to Data and Software

*Code and documentation are available at:* https://github.com/FHIR-Aggregator *and* https://fhir-aggregator.org/

## Materials and methods

The NCPI FHIR Aggregator implements a modular architecture that transforms diverse biomedical data sources into a unified FHIR-compliant repository. **Figure 2** illustrates the complete data flow and architectural components.

**Figure 2:**
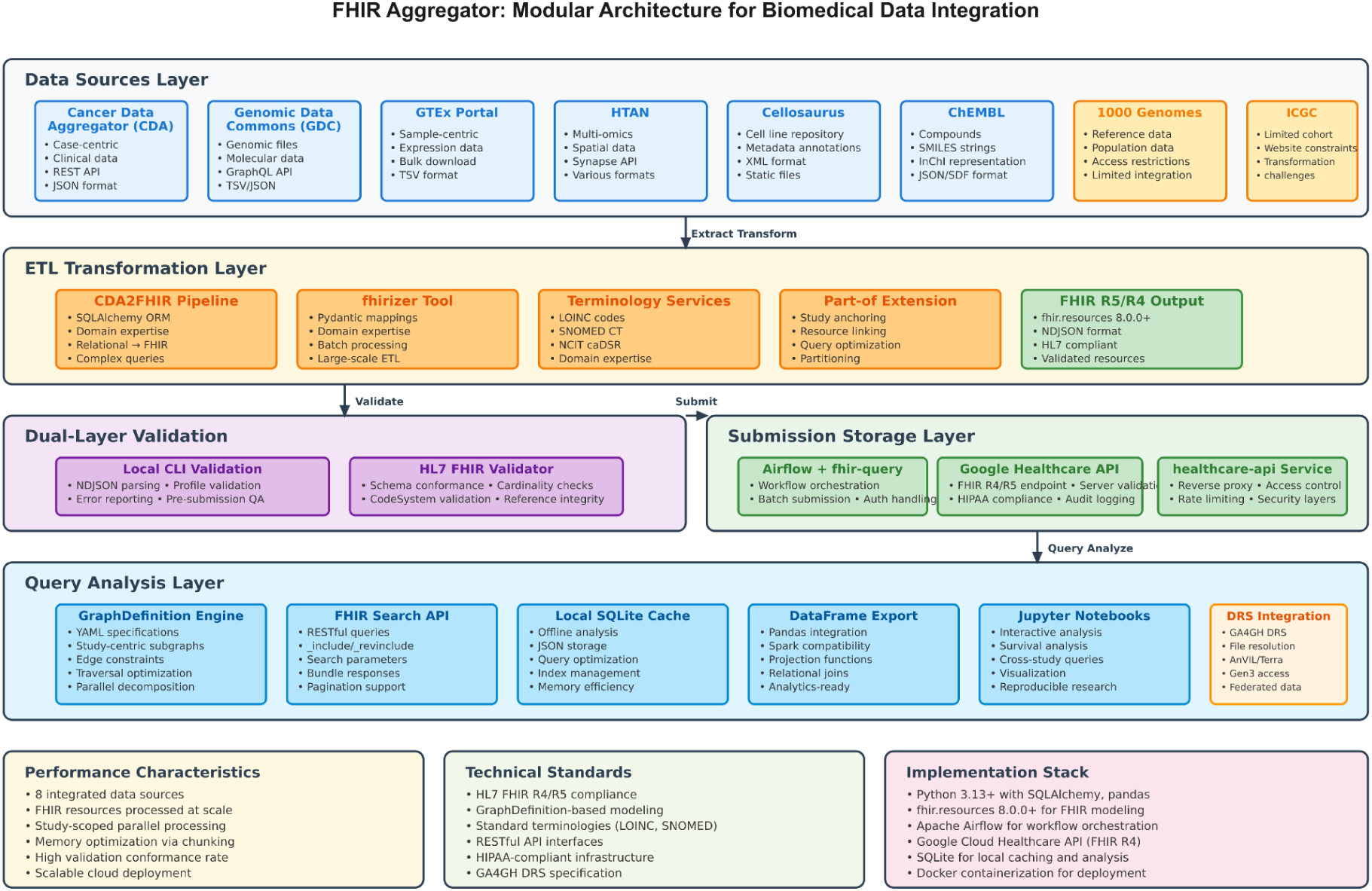
FHIR Aggregator modular architecture for biomedical data integration. The system implements a layered architecture spanning data extraction from eight biomedical sources, ETL transformation using domain expertise and Pydantic mappings (CDA2FHIR and fhirizer python client), dual-layer validation, Airflow-orchestrated submission to Google Healthcare API, and graph-based query interfaces with planned DRS integration for federated access. GraphDefinition-based subgraphs enable study-centric analysis while standardized terminologies ensure semantic consistency across heterogeneous data sources.

The architecture consists of five modular layers: data source integration, ETL transformation, validation, storage/submission, and query/analysis each tuned for scalability and performance. It operates through seven phases: (1) data extraction from heterogeneous sources, (2) schema transformation using SQLAlchemy ETL or Pydantic-based mappings, (3) local FHIR resource validation, (4) batch submission to Google Healthcare API via Airflow workflows, (5) server-side validation and storage, (6) graph-based query execution using GraphDefinition templates, and (7) local analysis using pandas dataframes. Reusable GraphDefinition templates enforce consistent traversal patterns across study designs. This design enables modular development, maintains FHIR compliance, and supports end-to-end data integrity.

### Schema Design and Subgraph Modeling

The NCPI FHIR Aggregator schema defines the underlying structure and data model used to organize FHIR resources such as Patient, Specimen, Observation, and Condition. By enforcing standardized attributes, relationships, and data types, it enables consistent data exchange and simplifies integration across heterogeneous platforms.

To support study-specific analytics, FHIR Aggregator employs HL7 GraphDefinition resources to define reusable subgraphs that reflect real-world study designs. These subgraphs link core entities such as patients, biospecimens, diagnoses, treatments and enable efficient traversal for clinically meaningful cohort extraction. Each subgraph is authored in YAML format and validated for structural and referential correctness.

A key transformation step embeds “part-of-study” relationships using FHIR extensions to explicitly associate each resource with its originating ResearchStudy. This enables efficient filtering, partitioning, and graph traversal scoped by study context. The overall schema structure used for subgraph modeling is shown in **Figure 1**. These standardized definitions promote harmonized traversal semantics across cancer types and study designs, ultimately increasing statistical power and supporting reproducible, cross-study analysis.

### Graph Representation and Flattening for Analysis

Each GraphDefinition describes a typed, directed graph G = (V, E), where nodes vᵢ ∈ V represent FHIR resources and edges e ∈ E encode typed references (ex. Observation.subject to Patient). For analytical workflows, we define a projection function X: V → ℝᵈ, where X(vᵢ) extracts attributes into a row vector. The resulting matrix A = [X(v₁), X(v₂), …, X(v□)] enables transformation into tabular formats compatible with pandas, Spark, and SQL-based systems.

Graph traversal adheres to GraphDefinition constraints, employing depth-first strategies for nested references and breadth-first traversal for cohort-based retrieval anchored by study resources. “Part-of-study” extension relationships introduced during transformation enable performance optimizations, including: (1) pruning traversal via pre-filtering by study, (2) parallel processing across studies, (3) reduced memory footprint via study-level chunking, and (4) accelerated study-specific analyses. These optimizations support both lightweight local use (ex. SQLite) and scalable distributed execution.

### Data Ingestion and Schema Transformation

Data ingestion serves as the primary transformation layer, where heterogeneous source schemas are mapped to standardized FHIR resources through diverse approaches tailored to each data source. The CDA2FHIR pipeline uses SQLAlchemy-based transformation for complex relational mappings, while large-scale transformations of GDC, HTAN, Cellosaurus cell lines, and ICGC cohorts are handled through the fhirizer python client, which leverages computational biology expertise and Pydantic class representation of mapping dictionaries for efficient FHIR resource generation. At the time of implementation, ICGC integration was limited due to new website and resource transformation constraints.

The transformation process incorporates standardized medical terminologies to ensure semantic consistency: LOINC codes for laboratory observations and measurements, SNOMED CT for clinical concepts and procedures, and NCIT caDSR for body structure codes and condition classifications. Patient observations critical for survival analysis, such as days to death, are mapped using appropriate LOINC codes to enable consistent querying across studies. This terminology standardization addresses schema alignment challenges including mapping different field structures, normalizing clinical vocabularies, and maintaining valid relationships between related resources.

### Client Architecture

The system provides a Python client library for programmatic access to aggregated data, supporting both local and remote FHIR server deployments. Authentication and authorization are handled through Google Cloud’s reverse proxy infrastructure, enabling secure access to sensitive biomedical datasets while maintaining FHIR compliance.

Cloud deployment is managed through containerized infrastructure with SSL termination and reverse proxy configurations. The deployment stack includes a SWAG reverse proxy for SSL handling, secure proxies to Google Healthcare API, and static content serving, all orchestrated through Docker Compose for reproducible deployments across different environments.

### FHIR Resource Submission and Healthcare API

After transformation and validation, FHIR resources are submitted to Google Cloud Healthcare API through the fhir-query submission pipeline. This process handles authentication, batch resource creation, and error handling during the upload to the managed FHIR store. The healthcare-api component manages the reverse proxy configuration and access controls, enabling secure programmatic access to the submitted resources while maintaining HIPAA compliance and audit trails.

Overall, this modular design separates concerns between data transformation (CDA2FHIR, fhirizer), graph modeling (graph-definitions), resource submission (fhir-query), infrastructure management (healthcare-api), cloud deployment (cloud), and client access (fhir-aggregator-client), supporting independent development and deployment of each component.

### Validation and Quality Assurance

Dual-layer validations are incorporated to ensure FHIR compliance and data quality. First, a CLI validation tool processes NDJSON files in the META folder, validating transformed resources against FHIR profiles and reporting structural or semantic errors with precise line-by-line feedback. This validation checks referential integrity by ensuring resource references point to existing resources, and catches transformation errors early in the pipeline.

Second, the Google FHIR API provides server-side validation during resource submission, enforcing FHIR conformance and rejecting non-compliant resources. This dual approach ensures both local development validation and production-grade compliance.

### Graph Definition Schema, Query Interface and Local Analysis

Study-specific subgraphs are defined using YAML-based GraphDefinition resources. For instance, a cholangiocarcinoma-specific graph outlines the relationships among Patients diagnosed with cholangiocarcinoma and their associated Specimens, Observations, and ResearchStudy resources, capturing traversal paths from study enrollment through specimen collection to downstream molecular observations. These declarative subgraph definitions enable consistent traversal patterns across studies sharing the same diagnosis constraints, supporting reproducible analytics and enhancing downstream analysis statistical power.

The NCPI FHIR Aggregator includes a locally hosted CLI tool that enables graph-based traversal of interconnected FHIR resources, accessible to users across a range of technical backgrounds. Key features include GraphDefinition-driven traversals using R5/R4 GraphDefinition objects, local SQLite storage for offline use, seamless conversion to pandas dataframes for analysis, and a reusable library of YAML-based GraphDefinition templates.

### Pipeline Orchestration

The transformation architecture employs specialized tools suited to each data source’s scale and characteristics. CDA transformations use SQLAlchemy for complex relational mappings, while large-scale data sources (GDC, HTAN, Cellosaurus cell lines, ICGC limited cohorts) are processed through the fhirizer python client, which provides efficient batch transformation capabilities. Integration challenges vary by source - ICGC’s limited cohort reflects constraints in website accessibility and resource transformation complexity, while other sources enable more comprehensive data integration.

### Subgraph Design

YAML-based GraphDefinition specifications prove effective for capturing study-specific data relationships. The declarative approach enables non-technical stakeholders to review and modify graph structures while ensuring technical validity. Disease-specific graphs like CDA Cholangiocarcinoma GraphDefinition demonstrate how the framework adapts to different research study design contexts while maintaining consistent traversal semantics.

### Validation and Quality Control

The dual-layer validation approach proves essential for maintaining data quality across heterogeneous sources. CLI-based validation of NDJSON files enables rapid iteration during development, providing immediate feedback on transformation errors with precise error locations and JSON context. The Google FHIR API’s server-side validation serves as a final quality gate, ensuring only compliant resources reach the production environment. This layered approach catches both structural FHIR violations and semantic inconsistencies that could compromise downstream analysis.

## Supplementary Figures and Tables

**Figure S1.**
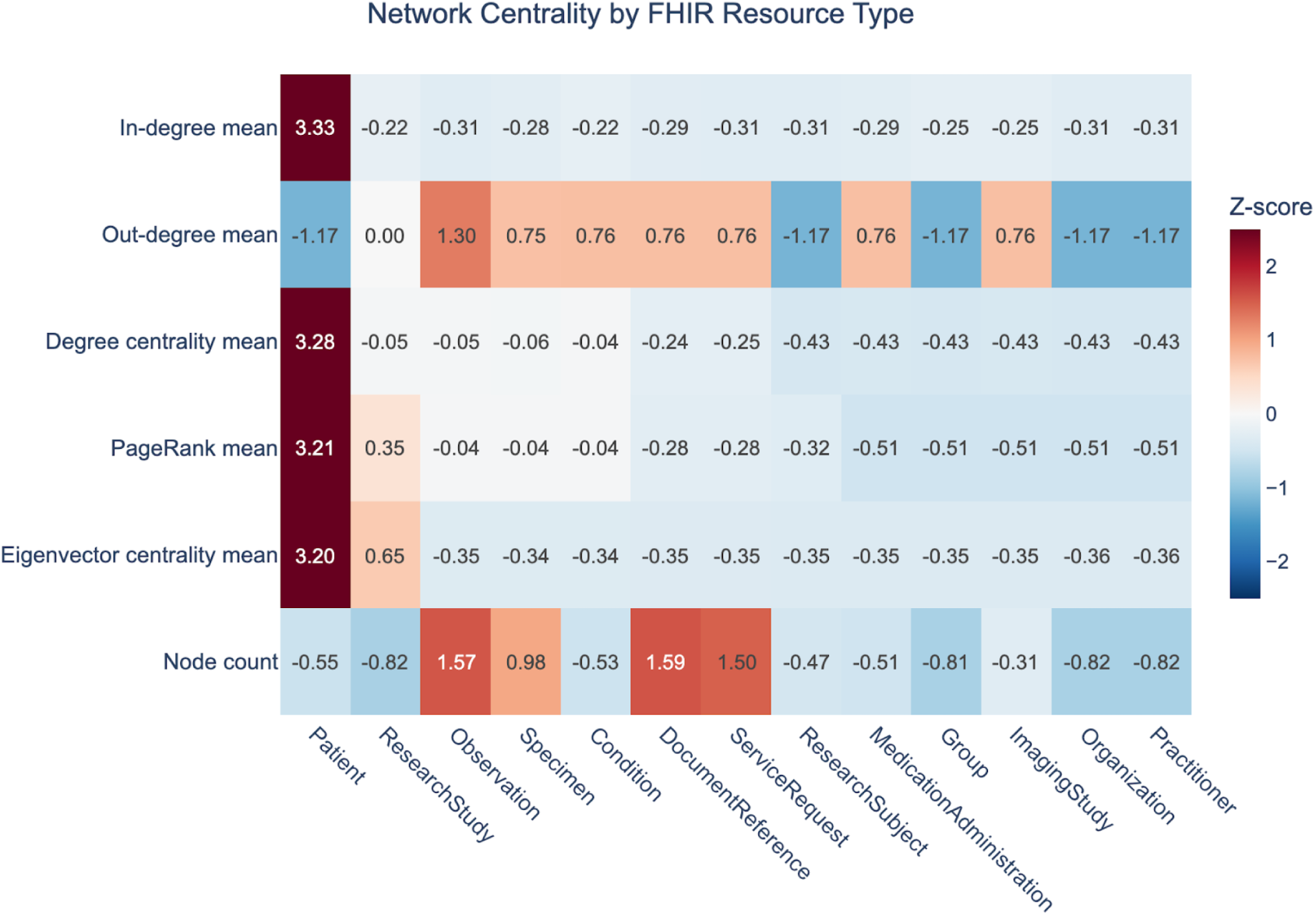
Z-score normalized network centrality metrics across FHIR resource types. Each cell represents the standardized (z-score) value of a FHIR resource type across five graph metrics: in-degree, out-degree, degree centrality, PageRank, and eigenvector centrality. Higher z-scores (red) indicate above-average connectivity or influence within the resource graph, while lower scores (blue) indicate less central roles. The Patient entity exhibits the highest centrality across most metrics, while Observation shows elevated out-degree, consistent with its role linking patients, specimens, and metadata. These results quantitatively confirm structural patterns inherent to the FHIR schema, such as the central role of Patient and the connector role of Observation in cross-resource linking. The analysis was performed using harmonized FHIRized datasets for CDA, GDC, GTEx, HTAN, ICGC, CELLOSAURUS, and 1000Genome, each including some or all of the following resource files: Patient.ndjson, Observation.ndjson, Specimen.ndjson, ResearchStudy.ndjson, ResearchSubject.ndjson, ServiceRequest.ndjson, and DocumentReference.ndjson; and, where applicable, Condition.ndjson, MedicationAdministration.ndjson, Organization.ndjson, Practitioner.ndjson, and ImagingStudy.ndjson.

**Supplemental Table 1.** FHIR Observation types and component structure generated by vcf2fhir. Structure of FHIR Observations produced during vcf2fhir conversion of VCF files. Each Observation type is identified by LOINC code and contains components mapping to specific FHIR paths. Components marked with asterisk (*) are included only when clinical annotations are provided. Source (https://github.com/elimuinformatics/vcf2fhir)

| 1. Variant Observation (Observation.code = LOINC 69548-6) |  |  |  |
| --- | --- | --- | --- |
| LOINC Code | Component Description | FHIR Path | Required |
| 48004-6 | DNA change (c.HGVS) | Observation.component | Yes |
| 48013-7 | Genomic reference sequence ID | Observation.component | Yes |
| 48002-0 | Genomic source class | Observation.component | Optional |
| 53034-5 | Allelic state | Observation.component | Optional |
| 69547-8 | Genomic ref allele | Observation.component | Yes |
| 69551-0 | Genomic alt allele | Observation.component | Optional |
| 81254-5 | Genomic allele start-end | Observation.component | Yes |
| 92822-6 | Genomic coordinate system | Observation.component | Yes |
| 48019-4 | DNA change type (variant type) | Observation.component | Optional |
| 81258-6 | Sample allelic frequency | Observation.component | Optional |
| 82155-3 | Copy number | Observation.component | Optional |
| 48018-6 | Gene studied* | Observation.component | Optional |
| 51958-7 | Transcript reference sequence* | Observation.component | Optional |
| 48005-3 | Amino acid change (p.HGVS)* | Observation.component | Optional |
| 53037-8 | Clinical significance* | Observation.component | Optional |
| 2. Region Studied Observation (Observation.code = LOINC 53041-0) |  |  |  |
| LOINC Code | Component Description | FHIR Path | Required |
| 92822-6 | Genomic coordinate system | Observation.component | Yes |
| 48013-7 | Genomic reference sequence ID | Observation.component | Yes |
| 51959-5 | Range(s) of DNA sequence examined | Observation.component | Yes |
| uncallable-region | Uncallable regions (custom) | Observation.component | Optional |
| 3. Sequence Phase Relationship Observation (Observation.code = LOINC 82120-7) |  |  |  |
| LOINC Code | Component Description | FHIR Path | Required |
| N/A | Allelic phase (Cis/Trans) | Observation.valueCodeableConcept | Yes |
| 4. Diagnostic Implication Observation (Observation.code = LOINC 51968-6) |  |  |  |
| LOINC Code | Component Description | FHIR Path | Required |
| 48018-6 | Gene studied | Observation.component | Yes |
| 53037-8 | Clinical significance | Observation.component | Yes |
| 81259-4 | Associated phenotype | Observation.component | Optional |

