## Supplementary Analysis and Notebooks for "FHIR as a Unifying Format for Genomic Research Data Tracking, Aggregation, and Integration"

### FHIR-Aggregator: A Catalog of Research Data¶

The FHIR Aggregator acts as a centralized repository for diverse healthcare data, organized using the FHIR (Fast Healthcare Interoperability Resources) standard. It provides researchers access to a wide range of information, including:

- Clinical data: Patient demographics, conditions, medications, observations, and procedures.
- Research studies: Information about research projects, participants, and study protocols.
- OMICS data associated with Specimens

#### Overview¶

This notebook leverages **FHIR GraphDefinition** objects to define and execute graph-based traversals across multiple interconnected FHIR resource graphs. The data retrieved is written to a **local SQLite database** for persistence and later transformed into **analyst-friendly dataframes** for analysis using tools like Python's pandas library.

##### Motivation¶

**FHIR Search** provides a robust querying framework but comes with significant limitations:

1. **Deep Chaining Limits**:
Chaining searches (e.g., `Patient -> Observation -> Encounter -> Procedure`) often hits server depth limitations.

2. **Inefficient Query Execution**:
Searching deeply related resources requires multiple chained requests, leading to performance issues and unnecessary round trips.

3. **Lack of Explicit Traversals**:
Relationships in FHIR are implicit in references (e.g., `Observation.subject` pointing to `Patient`). This implicit structure requires manual composition of queries, which is prone to errors.

4. **Differences between servers**:
Relationships in FHIR are implicit in references (e.g., `Observation.subject` pointing to `Patient`). This implicit structure requires manual composition of queries, which is prone to errors. **Key differences:**

1. Supported Search Parameters:

The FHIR specification defines a core set of search parameters for each resource type. However, servers might implement only a subset of these parameters or extend them with custom ones. This means a search that works on one server might not be valid on another. 2. Chaining and Search Depth:

Servers have different limits on how deeply you can chain search parameters (e.g., searching for Patients with Observations linked to specific Encounters). Some servers might restrict chaining depth, impacting the complexity of queries you can execute. 3. Search Parameter Behavior:

Even for commonly supported parameters, the specific behavior might differ. For instance, how date ranges are handled or how string searches are performed can vary, leading to unexpected results when switching between servers. 4. Terminology Services:

Servers might use different terminology services for coding systems, impacting how searches using codes are resolved. This can lead to discrepancies in results if the terminology is not aligned. 5. Performance and Optimization:

Search performance can vary significantly due to server infrastructure, indexing strategies, and data volume. Certain search patterns might be optimized on one server but inefficient on another. 6. Data Models and Link Population:

Data Models: FHIR servers may implement different profiles or extensions on top of the base FHIR specification. This can affect the structure of resources and the availability of certain data elements relevant to research. Link Population: The extent to which servers populate links (references) between resources can vary. Some servers might aggressively pre-fetch linked resources, while others may require explicit chaining or separate requests to retrieve them. This can significantly impact search performance and the complexity of queries needed to access related data for analysis.

#### Solution¶

###### fq (fhir-query): Your FHIR Querying Assistant¶

By using **FHIR GraphDefinition**, we declaratively define resource relationships and efficiently retrieve data. Once retrieved, the data is stored locally and can be transformed into dataframes for advanced analysis.

The fhir-aggregator-client tool runs an R5 GraphDefinition against a FHIR server.

##### Key Features¶

- **GraphDefinition-Driven Traversals**: Use GraphDefinition objects to define explicit relationships between resources and automate traversal logic.
- **Local SQLite Storage**: Persist the retrieved FHIR data in a local SQLite database for querying and offline analysis.
- **Analyst-Friendly Dataframes**: Convert stored FHIR resources into pandas dataframes for ease of use in analytical workflows.
- **Reusable Graph Definitions**: Maintain a library of GraphDefinition YAML files that can be reused across different workflows and projects. Researchers and Data submitters can publish GraphDefinition files to help others navigate their data.

##### Installation¶

The fq utility, short for "fhir-query," is a command-line tool specifically designed to simplify the process of interacting with FHIR servers. It provides researchers with a convenient way to:

1. Retrieve the vocabulary of a FHIR server: With the vocabulary command, fq fetches and summarizes the key data elements (CodeableConcepts and Extensions) used within the FHIR data. This creates a central vocabulary Dataframe that helps researchers identify important data elements and their usage within the server.

2. Execute queries to retrieve FHIR resources: Researchers can then use fq to execute FHIR queries using a readable syntax. This helps to retrieve and filter data from the FHIR Server based on various search parameters and criteria.

```
!pip install fhir-aggregator-client --no-cache-dir --quiet
!pip freeze | grep fhir_aggregator_client
```

```
Preparing metadata (setup.py) ... done
Building wheel for halo (setup.py) ... done
fhir_aggregator_client==0.2.2
```

###### Verify the tools were installed¶

```
!fq
```

```
Creating directory: /root/.fhir-aggregator
Usage: fq [OPTIONS] COMMAND [ARGS]...

  FHIR-Aggregator utilities.

Options:
  --version  Show the version and exit.
  --help     Show this message and exit.

Commands:
  ls          List all the installed GraphDefinitions.
  run         Run GraphDefinition queries.
  results     Work with the results of a GraphDefinition query.
  vocabulary  FHIR-Aggregator's key Resources and CodeSystems.
```

#### Useage¶

##### List the installed GraphDefinition files¶

```
!fq ls
```

```
| id                           |
description
                                                                                                                                                                        |
|----------------------------
+-------------------------------------------------------------------------------------------------------------------------------------------------------------------------
-------------------------------------------------------------------------------------------------------------------------------------------------------------------------
-------------------------------------------------------------------------------------------------------------------------------------------------------------------------
|
| research-study-part-of      | (FHIR-Aggregator) Retrieve a ResearchStudy and children. Uses part-of-study extension. fhir-query '/ResearchStudy?
identifier=TCGA-
BRCA'
                         |
| cholangiocarcinoma-graph    | (FHIR-Aggregator) Condition to ResearchStudy and children focusing on Observations [NCIT_C156418,NCIT_C156419,NCIT_C164934].
fhir-query '/Condition?
code:code=70179006'
      |
| file2patients               | (FHIR-Aggregator) Retrieve a ResearchStudy and it's related entities. DocumentReference resources are filtered by a study
design criterion, e.g.: - category:code = {data_category} (default: Transcriptome Profiling), {workflow_type} (default: STAR - Counts),
{experimental_strategy} (default: RNA-Seq), {access} (default: open) - type:code = {file_type} (default: TSV) while the Patient link can be filtered via list
of comma separated patient identifiers {patientFilter}. |
| research-study-link-iterate | (dbGAP) Retrieve ResearchStudy and children. Uses HAPI's deep linking. fhir-query '/ResearchStudy?
_id=phs001232'
                                                         |
| patient-survival-graph      | (FHIR-Aggregator) Retrieve Patient and Observations [NCIT_C156418,NCIT_C156419]. fhir-query '/ResearchStudy?identifier=TCGA-
BRCA'
                               |
| condition-graph             | (FHIR-Aggregator) Condition to ResearchStudy and children. fhir-query '/Condition?
code:code=70179006'
                                                                         |
```

##### Run a GraphDefiniton¶

```
%env FHIR_BASE= https://google-fhir.fhir-aggregator.org
```

```
env: FHIR_BASE=https://google-fhir.fhir-aggregator.org
```

```
!fq run cholangiocarcinoma-graph '/Condition?code:code=70179006'
```

```
cholangiocarcinoma-graph is valid FHIR R5 GraphDefinition
ℹ Fetching https://google-fhir.fhir-aggregator.org/Condition?code:code=70179006
ℹ Processing Condition with 389 resources
ℹ Processing 1 links for Condition in parallel.
ℹ Processing link: Patient/_id={ref} with 354 Condition(s)
✔ Processed link: Patient/_id={ref}
ℹ Processing Patient with 389 resources
ℹ Processing 10 links for Patient in parallel.
ℹ Processing link: ResearchSubject/individual={ref}&_include=ResearchSubject:study with 354 Patient(s)
```

```
ℹ Processing link: Group/member={ref} with 354 Patient(s)
ℹ Processing link: Specimen/subject={ref} with 354 Patient(s)
ℹ Processing link: Observation/subject={ref}&code=NCIT_C156418,NCIT_C156419,NCIT_C164934&_count=1000&_total=accurate with 354 Patient(s)
ℹ Processing link: Procedure/subject={ref} with 354 Patient(s)
ℹ Processing link: DocumentReference/subject={ref}&_count=1000&_total=accurate with 354 Patient(s)
✖ Could not find any resources for Patient->Group link: {'params': 'member={ref}&_count=1000&_total=accurate', 'sourceId': 'Patient', 'targetId': 'Group',
'path': 'Patient.id'}
ℹ Processing link: ImagingStudy/subject={ref}&_count=1000&_total=accurate with 354 Patient(s)
ℹ Processing link: MedicationAdministration/subject={ref}&_count=1000&_total=accurate with 354 Patient(s)
ℹ Processing link: Encounter/subject={ref}&_count=1000&_total=accurate with 354 Patient(s)
✔ Processed link: ResearchSubject/individual={ref}&_include=ResearchSubject:study
✔ Processed link: Group/member={ref}
✔ Processed link: Observation/subject={ref}&code=NCIT_C156418,NCIT_C156419,NCIT_C164934&_count=1000&_total=accurate
✔ Processed link: Procedure/subject={ref}
✔ Processed link: ImagingStudy/subject={ref}&_count=1000&_total=accurate
✔ Processed link: DocumentReference/subject={ref}&_count=1000&_total=accurate
✔ Processed link: Encounter/subject={ref}&_count=1000&_total=accurate
✔ Processed link: MedicationAdministration/subject={ref}&_count=1000&_total=accurate
✔ Processed link: Specimen/subject={ref}
ℹ Processing ResearchSubject with 354 resources
ℹ Processing Specimen with 411 resources
ℹ Processing ResearchStudy with 2715 resources
ℹ Processing 1 links for ResearchSubject in parallel.
ℹ Processing 1 links for Specimen in parallel.
ℹ Processing 1 links for ResearchStudy in parallel.
✔ Processed link: ResearchStudy/
ℹ Processing link: ServiceRequest/specimen={ref} with 2715 Specimen(s)
ℹ Processing link: DocumentReference/subject={ref}&_count=1000&_total=accurate with 18 ResearchStudy(s)
✔ Processed link: DocumentReference/subject={ref}&_count=1000&_total=accurate
✔ Processed link: ServiceRequest/specimen={ref}
Aggregated Results: {'Condition': 389, 'Patient': 354, 'ResearchStudy': 18, 'ResearchSubject': 411, 'Specimen': 2715}
database available at: /root/.fhir-aggregator/fhir-graph.sqlite
```

```
# !fq run  condition-graph '/Condition?code:text=cholangiocarcinoma'
```

##### Analyse Results¶

The graph represents relationships between different FHIR resources.Examples of FHIR resources include Patient, Condition, Observation, Procedure, etc.

Each node is labled as: <resource_type>/<count> the number of records of that type retrieved.

The edges in the graph are weighted. The thicker the line, the more connections there are between nodes.

```
# Create a graph of the results
!fq results visualize
```

```
Wrote: fhir-graph.html
```

```
# Read the locally stored HTML file containing a graph visualization and displaying it within the Jupyter notebook.

from IPython.display import HTML
with open('fhir-graph.html', 'r') as file:
    html_content = file.read()

# Set the display height (in pixels)
display(HTML("<div style='height: 800px;'>{}</div>".format(html_content)))
```

##### Create a dataframe of results¶

```
!fq results dataframe
```

```
Saved fhir-graph.tsv
```

```
import pandas as pd
```

```
df = pd.read_csv('fhir-graph.tsv', sep='\t')
```

```
df
```

| | specimen_identifier | specimen_id | specimen_type | specimen_part_of_study | patient_identifier | patient_subject_id | patient_subje |
|---|---|---|---|---|---|---|---|
| **0** | FM-AD.AD4664.e91d151b-d460-5f1d-afc4-01cc8b45d74e | fbe21d14-b473-5fcd-adbf-dbe463c2511b | Tumor | ResearchStudy/ b86ee080-2f2f-54c6-b6a8-c1674bb9... | AD4664 | FM.AD4664 | 68981 |
| **1** | FM-AD.AD4664.AD4664_slide | e33cc270-35ca-57b8-901e-1d6cee1b8f82 | Tumor | ResearchStudy/ b86ee080-2f2f-54c6-b6a8-c1674bb9... | AD4664 | FM.AD4664 | 68981 |
| **2** | FM-AD.AD4664.AD4664_sample | b8b85c83-a762-58fd-ace1-4ba515e7c38f | Tumor | ResearchStudy/ b86ee080-2f2f-54c6-b6a8-c1674bb9... | AD4664 | FM.AD4664 | 68981 |
| **3** | FM-AD.AD4664.AD4664_aliquot | ec378c89-6c10-50f5-888b-233681973ce5 | Tumor | ResearchStudy/ b86ee080-2f2f-54c6-b6a8-c1674bb9... | AD4664 | FM.AD4664 | 68981 |
| **4** | FM-AD.AD4664.24b66eb0-bed8-50a0-8051-ebb181c80b0e | 7c57e79b-7504-5490-aab5-d42b94cac0c6 | Tumor | ResearchStudy/ b86ee080-2f2f-54c6-b6a8-c1674bb9... | AD4664 | FM.AD4664 | 68981 |
| **...** | ... | ... | ... | ... | ... | ... | ... |
| **2710** | FM-AD.AD5125.8a71532f-b6bc-5f0a-9ddb-c9cb93037d4e | 7b60af59-6865-5e96-8521-01b12057831c | Tumor | ResearchStudy/ b86ee080-2f2f-54c6-b6a8-c1674bb9... | AD5125 | FM.AD5125 | 69494 |
| **2711** | FM-AD.AD5125.AD5125_aliquot | 39e64af9-54ac-5aff-9e31-fcc3a2deb700 | Tumor | ResearchStudy/ b86ee080-2f2f-54c6-b6a8-c1674bb9... | AD5125 | FM.AD5125 | 69494 |
| **2712** | FM-AD.AD5125.e9677cdc-53c0-54f7-88d7-f02c6a7cb73a | 0b8af889-b048-5d21-b993-9051a9e8521b | Tumor | ResearchStudy/ b86ee080-2f2f-54c6-b6a8-c1674bb9... | AD5125 | FM.AD5125 | 69494 |
| **2713** | FM-AD.AD5125.AD5125_slide | 40cec0fc-3ac6-5ca3-9d44-153e63b01749 | Tumor | ResearchStudy/ b86ee080-2f2f-54c6-b6a8-c1674bb9... | AD5125 | FM.AD5125 | 69494 |
| **2714** | FM-AD.AD5125.54aa9c6a-7d7f-5728-a520-66200c5dd492 | 7ea86835-d52d-5663-ae35-2793b7454ff1 | Tumor | ResearchStudy/ b86ee080-2f2f-54c6-b6a8-c1674bb9... | AD5125 | FM.AD5125 | 69494 |

2715 rows × 30 columns

##### Other servers¶

You can use the `fq` tool with other FHIR servers. For example, this query retrieves a study from `dbGAP`

```
# delete the previous results, start with a fresh database
!rm ~/.fhir-aggregator/fhir-graph.sqlite
!fq run  --fhir-base-url https://dbgap-api.ncbi.nlm.nih.gov/fhir-jpa-pilot/x1  research-study-link-iterate  '/ResearchStudy?_id=phs001232'
```

```
research-study-link-iterate is valid FHIR R5 GraphDefinition
ℹ Fetching https://dbgap-api.ncbi.nlm.nih.gov/fhir-jpa-pilot/x1/ResearchStudy?_id=phs001232
ℹ Processing ResearchStudy with 1 resources
ℹ Processing 1 links for ResearchStudy in parallel.
ℹ Processing link: Patient/_has:ResearchSubject:individual:study={ref}
&_revinclude=Group:member&_revinclude=ResearchSubject:individual&_revinclude=Specimen:subject&_revinclude=Observation:subject&_revinclude=DocumentReference:su
bject&_count=1000&_total=accurate with 1 ResearchStudy(s)
```

```
# use the same commands to analyse results
!fq results visualize
```

```
# create a graph of the results

from IPython.display import HTML
with open('fhir-graph.html', 'r') as file:
    html_content = file.read()

# Set the display height (in pixels)
display(HTML("<div style='height: 800px;'>{}</div>".format(html_content)))
```

```
# create a dataframe of results
!fq results dataframe
```

```
Saved fhir-graph.tsv
```

```
import pandas as pd

df = pd.read_csv('fhir-graph.tsv', sep='\t')

df
```

| | specimen_identifier | specimen_phs001232-SampleIdentifier | specimen_id | specimen_type | patient_identifier | patient_id | patient_active | patient_gender | patient_meta | pat |
|---|---|---|---|---|---|---|---|---|---|---|
| **0** | SAME123424 | f9964f35-00a8-4f0c-8ba2-6f848ea0fb0f | dgs-5063489 | Fibroblasts | NaN | 1826993 | True | male | phs001232 | NaN |
| **1** | SAME123417 | 6ba8fe4f-693d-40c9-8c69-981cc7b6ba87 | dgs-2211200 | Blood | NaN | 1826993 | True | male | phs001232 | NaN |
| **2** | SAME1839057 | 084ecc87-1588-4bb3-94d7-33942e3f0561 | dgs-4274478 | Blood | NaN | 1827029 | True | male | phs001232 | NaN |
| **3** | SAME1839246 | 08fcbd02-5729-4c9a-9a63-4ce48c400fe4 | dgs-2211148 | Blood | NaN | 1827029 | True | male | phs001232 | NaN |
| **4** | SAME124161 | db37d721-4b6a-4de7-8e37-38d3a00f9aca | dgs-2211247 | Blood | NaN | 1827062 | True | male | phs001232 | NaN |
| **...** | ... | ... | ... | ... | ... | ... | ... | ... | ... | ... |
| **6408** | SAME122976 | a5679b33-ff69-48e7-9f82-6bbd0a9b4049 | dgs-5063129 | Blood | NaN | 4296843 | True | male | phs001232 | NaN |
| **6409** | SAME1839098 | ced2fb56-6535-4e52-b755-dd16263925f1 | dgs-5063313 | Blood | NaN | 4296875 | True | female | phs001232 | NaN |
| **6410** | SAME1839147 | e5ede4ac-e8a3-447d-a654-0c6319af9c5c | dgs-5063407 | Blood | NaN | 4296515 | True | male | phs001232 | NaN |
| **6411** | SAME123105 | f8240565-ff65-4742-a299-99c01bab31ab | dgs-5063482 | Blood | NaN | 4297305 | True | male | phs001232 | NaN |
| **6412** | SAME123917 | 98af2a6c-7f17-4612-832d-05d1d104e1ea | dgs-5063082 | Blood | NaN | 4296730 | True | female | phs001232 | NaN |

6413 rows × 21 columns

### FHIR-Aggregator survival anaysis¶

In this notebook we will show how to retrieve data from breast cancer patients in TCGA and compare the Kaplan-Meier curves of two cohorts. The cohorts are white and african american pateints that are 50 years or younger.

#### Install necessay packages¶

```
!pip install lifelines -q
```

```
  Preparing metadata (setup.py) ... done
   ━━━━━━━━━━━━━━━━━━━━━━━━━━━━━━━━━━━━━━━━ 349.3/349.3 kB 5.3 MB/s eta 0:00:00
   ━━━━━━━━━━━━━━━━━━━━━━━━━━━━━━━━━━━━━━━━ 117.2/117.2 kB 8.2 MB/s eta 0:00:00
  Building wheel for autograd-gamma (setup.py) ... done
```

```
!pip install fhir-aggregator-client --no-cache-dir --quiet
!pip freeze | grep fhir_aggregator_client
```

```
  Preparing metadata (setup.py) ... done
     ━━━━━━━━━━━━━━━━━━━━━━━━━━━━━━━━━━━━━━━━ 49.2/49.2 kB 3.3 MB/s eta 0:00:00
   ━━━━━━━━━━━━━━━━━━━━━━━━━━━━━━━━━━━━━━━━ 2.3/2.3 MB 36.2 MB/s eta 0:00:00
   ━━━━━━━━━━━━━━━━━━━━━━━━━━━━━━━━━━━━━━━━ 756.0/756.0 kB 53.8 MB/s eta 0:00:00
   ━━━━━━━━━━━━━━━━━━━━━━━━━━━━━━━━━━━━━━━━ 1.6/1.6 MB 127.3 MB/s eta 0:00:00
  Building wheel for halo (setup.py) ... done
fhir_aggregator_client==0.2.1
```

#### Use FHIR-Aggregator to retrieve the necessary data¶

##### Export TCGA-BRCA data to a local database¶

```
# run query against released data
# !rm /root/.fhir-aggregator/fhir-graph.sqlite
%env  FHIR_BASE=https://google-fhir.fhir-aggregator.org
!fq run patient-survival-graph    '/ResearchStudy?identifier=TCGA-BRCA'
```

```
env: FHIR_BASE=https://google-fhir.fhir-aggregator.org
Creating directory: /root/.fhir-aggregator
patient-survival-graph is valid FHIR R5 GraphDefinition
i Fetching https://google-fhir.fhir-aggregator.org/ResearchStudy?identifier=TCGA-BRCA
i Processing ResearchStudy with 1 resources
i Processing 2 links for ResearchStudy in parallel.
i Processing link: Patient/part-of-study={ref}&_count=1000&_total=accurate with 1 ResearchStudy(s)
i Processing link: Observation/part-of-study={ref}&code=NCIT_C156418,NCIT_C156419&_count=1000&_total=accurate with 1 ResearchStudy(s)
✔ Processed link: Patient/part-of-study={ref}&_count=1000&_total=accurate
✔ Processed link: Observation/part-of-study={ref}&code=NCIT_C156418,NCIT_C156419&_count=1000&_total=accurate
Aggregated Results: {'Observation': 1231, 'Patient': 1098, 'ResearchStudy': 1}
database available at: /root/.fhir-aggregator/fhir-graph.sqlite
```

##### Create a tsv file from the extracted data¶

```
# The previous query included a Specimen,  the dataframe type defaults to Specimen
# Since the optimized query only has Patient, we as for a Patient dataframe type
# Note: default output is in the current directory and is a TSV
!fq results dataframe Patient
```

```
WARNING:root:ResearchSubject, which maps patient to study, not found. Useful for multi study queries.  patient: 2e75d38a-da90-5ff1-b469-906e4132ab9c
Saved fhir-graph.tsv
```

#### Surviving analysis¶

After retrieveing the data, we then use the python library lifelines to plor Kaplan-Meier plots of two groups (white and african american) of Breat cancer patients that are 50 years old or younger.

```
import pandas as pd
import numpy as np
import matplotlib.pyplot as plt
from lifelines import KaplanMeierFitter
kmf = KaplanMeierFitter()


# read the data into a dataframe
df = pd.read_csv('fhir-graph.tsv', sep='\t')

# get days to death data in the necessary format
df['days_to_death'] = (
    df['observation_days_between_diagnosis_and_death']
    .str.replace(' days', '', regex=False)
    .replace('', np.nan)
    .astype(float)
)
# get age data in the necessary format
df['age_at_diagnosis'] = (
    df['observation_days_between_birth_and_diagnosis']
    .str.replace(' days', '', regex=False)
    .replace('', np.nan)
    .astype(float)
)

# group by patient_id
df_unique = df.drop_duplicates(subset=['patient_id'])
```

Select Breast cancer patients that are white, african american, and 50 years old or younger.

```
df_cohort = df_unique[ (df_unique['age_at_diagnosis'] >= -50*365 )
                      & (df_unique['patient_us_core_race'].isin(['black or african american','white']) )
                      & (df_unique['patient_us_core_ethnicity'] == 'not hispanic or latino')   ]
```

Get the necessary data for `lifelines` package.

```
# Fill in NAs in days_to_death with the max from the days to death
T = df_cohort['days_to_death'].fillna(df_cohort['days_to_death'].max())

# Convert the vital status to numbers
E = df_cohort['patient_deceasedBoolean'].astype(bool)
```

Plot the survivial curves

```
fig=plt.figure(figsize=(13, 8), dpi= 80)
#plt.style.use('seaborn-colorblind')
ax = plt.subplot(111,
                 title = "Survival Curve")

for r in  df_cohort['patient_us_core_race'].sort_values().unique() :
  if (r != None):
    cohort = df_cohort['patient_us_core_race'] == r
    kmf.fit(T.loc[cohort], E.loc[cohort], label=r)
    kmf.plot(ax=ax, )
  else:
```

```
    print("")

ax.set_ylabel("Percent Survival")
ax.set_xlabel("Days")
```

```
Text(0.5, 0, 'Days')
```

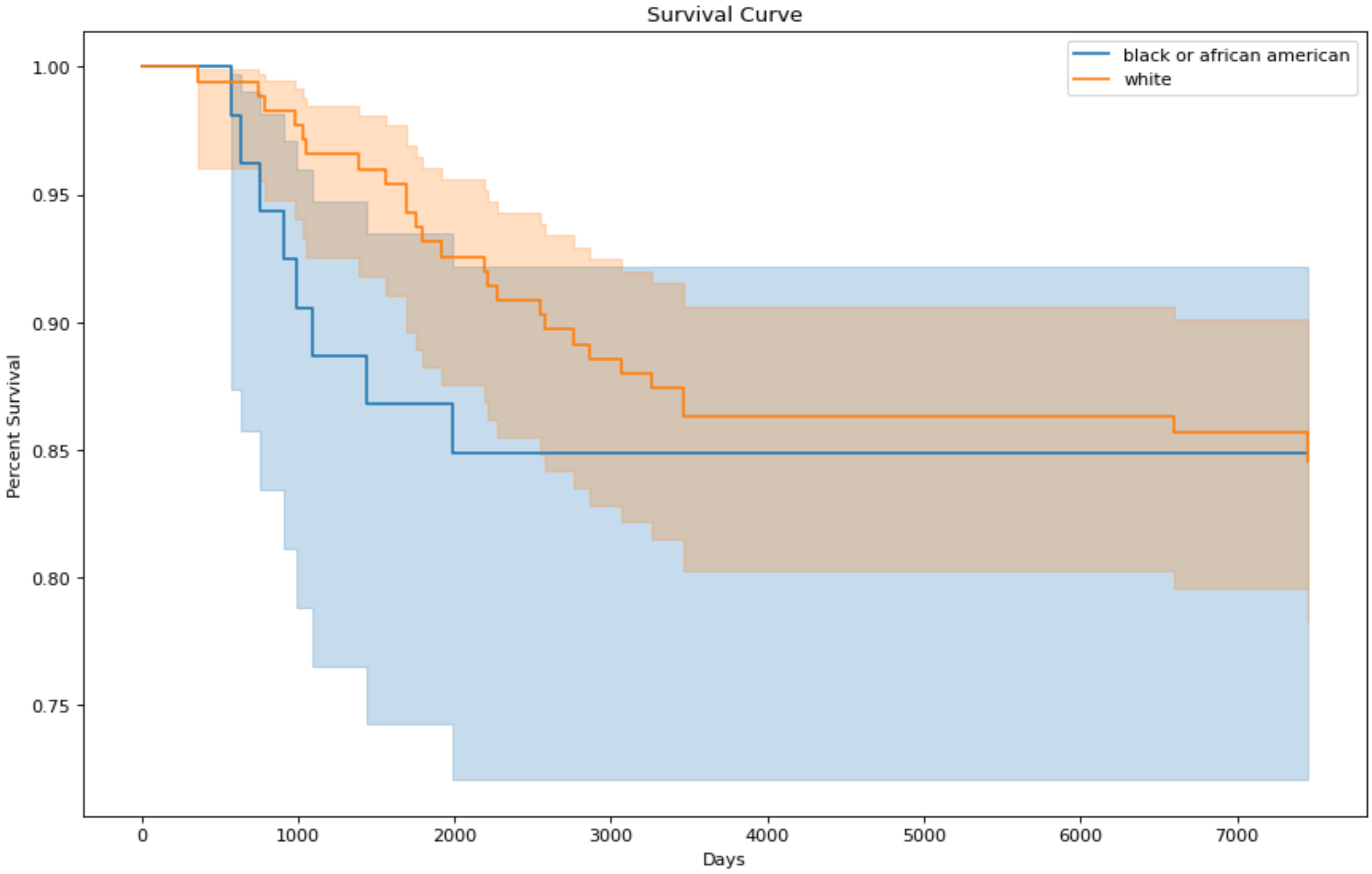

### FHIR-Aggregator: A Catalog of Research Data¶

The FHIR Aggregator acts as a centralized repository for diverse healthcare data, organized using the FHIR (Fast Healthcare Interoperability Resources) standard. It provides researchers access to a wide range of information, including:

- Clinical data: Patient demographics, conditions, medications, observations, and procedures.
- Research studies: Information about research projects, participants, and study protocols.
- OMICS data associated with Specimens

#### Specify the endpoint¶

- We need to select the FHIR Server's URL https://google-fhir.fhir-aggregator.org
  - This line of code tells the notebook, "Remember this address: https://google-fhir.fhir-aggregator.org, and label it FHIR_BASE. We'll use it later to talk to a server that stores healthcare data."
  - By setting this environment variable, the URL to the FHIR Aggregator server is conveniently stored for later use within the notebook. This way you won't need to repeat the URL every time it's needed.
- From there we have access to search the data in the server using FHIR queries

```
%env FHIR_BASE= https://google-fhir.fhir-aggregator.org
```

```
env: FHIR_BASE=https://google-fhir.fhir-aggregator.org
```

### Example FHIR query¶

Now that you have the endpoint, if you are comfortable with FHIR, that is all you need. For example:

This query returns the official identifier for all ResearchStudy resources.

- $FHIR_BASE is the environment variable we set earlier, which holds the FHIR server's base URL. It's expanded to the actual URL during execution.
- /ResearchStudy is the FHIR resource type we are interested in (in this case, "ResearchStudy").
- ?_elements=identifier is a FHIR search parameter that limits the returned data to only include the 'identifier' element of the ResearchStudy resources.

```
# Install the jq json formatter tool
!apt-get install -yq jq > /dev/null
!jq --version

! curl -s $FHIR_BASE'/ResearchStudy?_elements=identifier&identifier.use=official' | jq -rc '.entry[] | [ (.resource.identifier[] | .value), .fullUrl]' | sort
```

```
jq-1.6
parse error: Invalid numeric literal at line 1, column 7
```

- Let's craft the code to query the FHIR server and load the results into a Pandas DataFrame.

```
import requests
import pandas as pd
import json

# Assuming FHIR_BASE is already set as an environment variable
fhir_base_url = %env FHIR_BASE
```

```
# Define the API endpoint
endpoint = f"{fhir_base_url}/ResearchStudy?_elements=identifier&identifier.use=official"

# Make the request
response = requests.get(endpoint)

# Check for successful response
if response.status_code == 200:
    # Parse the JSON response
    data = response.json()

    # Extract identifiers
    identifiers = []
    for entry in data.get('entry', []):
        resource = entry.get('resource', {})
        for identifier in resource.get('identifier', []):
            # add the url RearchStudy to the dataframe
            identifier['url'] = entry.get('fullUrl')
            identifiers.append(identifier)

    # Create a Pandas DataFrame
    print(f"Found {len(identifiers)} ResearchStudy identifiers. Use the 'url' field to retrieve the data.")
    df = pd.DataFrame(identifiers)
    display(df)  # Display the DataFrame
else:
    print(f"Error: Request failed with status code {response.status_code}")
```

```
Found 151 ResearchStudy identifiers. Use the 'url' field to retrieve the data.
```

| | system | use | value | url |
|---|---|---|---|---|
| **0** | https://aced-idp.org/ICGC-LUCA_KR | official | ICGC-LUCA_KR | https://google-fhir.fhir-aggregator.org/Resear... |
| **1** | https://gtexportal.org/home/downloads/adult-gt... | NaN | GTEX_V10 | https://google-fhir.fhir-aggregator.org/Resear... |
| **2** | https://https://ftp.1000genomes.ebi.ac.uk/vol1... | NaN | 1KG | https://google-fhir.fhir-aggregator.org/Resear... |
| **3** | https://data.humantumoratlas.org | official | WUSTL | https://google-fhir.fhir-aggregator.org/Resear... |
| **4** | https://data.humantumoratlas.org | official | CHOP | https://google-fhir.fhir-aggregator.org/Resear... |
| **...** | ... | ... | ... | ... |
| **146** | https://gdc.cancer.gov/project | official | TARGET-ALL-P3 | https://google-fhir.fhir-aggregator.org/Resear... |
| **147** | https://gdc.cancer.gov/project | official | TCGA-LUAD | https://google-fhir.fhir-aggregator.org/Resear... |
| **148** | https://gdc.cancer.gov/project | official | TCGA-COAD | https://google-fhir.fhir-aggregator.org/Resear... |
| **149** | https://gdc.cancer.gov/project | official | CGCI-BLGSP | https://google-fhir.fhir-aggregator.org/Resear... |
| **150** | https://gdc.cancer.gov/dbgap_accession_number | secondary | phs000527 | https://google-fhir.fhir-aggregator.org/Resear... |

151 rows × 4 columns

### Vocabulary dataframe¶

#### The Vocabulary DataFrame: A Researcher's Guide to Data Elements¶

Imagine you have a vast collection of FHIR data, containing medical records, research studies, and various observations. Within this data, there are numerous CodeableConcepts and Extensions that provide structure and meaning to the information. However, as a researcher, it's crucial to have a clear overview of these key data elements and how they're used.

**This is where the Vocabulary DataFrame comes in.**

The Vocabulary DataFrame is essentially a summary table that catalogs the important CodeableConcepts and Extensions found within the FHIR dataset. It acts as a central inventory, providing researchers with valuable insights into the structure and content of the data.

**Here's how it helps:**

**Identifying Key Data Elements**: The DataFrame lists all the significant CodeableConcepts and Extensions used within the data, giving researchers a comprehensive view of the elements present.

**Understanding Code Systems and Terminology**: It provides information about the code systems and terminologies associated with each CodeableConcept (e.g., SNOMED CT, LOINC), helping researchers interpret the coded data.

**Exploring Data Structure and Usage**: The DataFrame reveals where these CodeableConcepts and Extensions are used within different FHIR resources and elements (e.g., Condition.code, Observation.valueCodeableConcept). This helps researchers understand how the data is structured and how these elements relate to each other.

**Navigating to Specific Data**: It often includes FHIR queries that can be used to directly access the data associated with each CodeableConcept or Extension, making it easier to locate specific information.

**Facilitating Data Analysis**: By providing a structured inventory of key data elements, the Vocabulary DataFrame simplifies data exploration, analysis, and the formulation of research questions.

**Analogy**:

Think of the Vocabulary DataFrame as a library catalog. Just as a catalog helps you find books based on author, title, or subject, the DataFrame helps you find specific data elements within the FHIR dataset based on their code system, display, or where they are used.

**Example**:

The Vocabulary DataFrame contains columns like:

`research_study_identifiers`: Linking the CodeableConcept/extension to the study. `path`: Showing the FHIR element where the code is used (e.g., Condition.code). `system`: Indicating the code system (e.g., [http://snomed.info/sct](http://snomed.info/sct)). `display`: Providing a human-readable label for the code (e.g., Diabetes mellitus). `url`: Linking to a FHIR query to retrieve more information.

This structure empowers researchers to:

- Quickly identify all the conditions documented in studies using SNOMED CT codes.
- Explore the range of medications recorded in the dataset.
- Locate specific observations related to tumor grades.

**In essence, the Vocabulary DataFrame serves as a valuable tool for researchers, providing a structured overview of the key data elements within the FHIR dataset, enabling them to effectively explore, analyze, and understand the available information. **

### Retrieve vocabularies used on commonly used resources¶

When a study is submitted to the site, we survey the data and create an Observation of the data in the Study. These summary Observations are published on the server.

These summaries can inform researchers who need to formulate queries. e.g. for all studies:

- what are the conditions?
- what condition stages?
- what are the tumor grades ?
- what are the medications?
- what are the document types?

We can query the data using a FHIR query.

```
# get the native FHIR vocabulary Observation
! curl -s $FHIR_BASE'/Observation?code=vocabulary' | jq . | head -50
```


```
{
  "entry": [
    {
      "fullUrl": "https://google-fhir.fhir-aggregator.org/Observation/4fc6bcaa-bb07-51e9-99b2-a85f679d81ab",
      "resource": {
        "code": {
          "coding": [
            {
              "code": "vocabulary",
              "display": "Vocabulary",
              "system": "http://fhir-aggregator.org/fhir/CodeSystem/vocabulary"
            }
          ]
        },
        "component": [
          {
            "code": {
              "coding": [
                {
                  "code": "ServiceRequest.category",
                  "display": "ServiceRequest.category",
                  "system": "http://fhir-aggregator.org/fhir/CodeSystem/vocabulary/path"
                },
                {
                  "code": "108252007",
                  "display": "Laboratory procedure",
                  "system": "http://snomed.info/sct"
                }
              ]
            },
            "valueInteger": 15063
          },
          {
            "code": {
              "coding": [
                {
                  "code": "ServiceRequest.code",
                  "display": "ServiceRequest.code",
                  "system": "http://fhir-aggregator.org/fhir/CodeSystem/vocabulary/path"
                },
                {
```

```
                "code": "15220000",
                "display": "Laboratory test",
                "system": "http://snomed.info/sct"
              }
            ]
          },
          "valueInteger": 15063
        },
        {
```


#### Install the query tool¶

##### fq (fhir-query): Your FHIR Querying Assistant¶

The fq utility, short for "fhir-query," is a command-line tool specifically designed to simplify the process of interacting with FHIR servers. It provides researchers with a convenient way to:

1. Retrieve the vocabulary of a FHIR server: With the vocabulary command, fq fetches and summarizes the key data elements (CodeableConcepts and Extensions) used within the FHIR data. This creates a central vocabulary Dataframe that helps researchers identify important data elements and their usage within the server.
2. Execute queries to retrieve FHIR resources: Researchers can then use fq to execute FHIR queries using a readable syntax. This helps to retrieve and filter data from the FHIR Server based on various search parameters and criteria.

##### jq: Your JSON Navigator¶

The jq utility is a lightweight and flexible command-line JSON processor. It's like a Swiss Army knife for working with JSON data, and it is used here to help organize the responses from the FHIR server in a readable way.

```
!pip install fhir-aggregator-client --no-cache-dir --quiet
!pip freeze | grep fhir_aggregator_client
```

```
  Preparing metadata (setup.py) ... done
     ━━━━━━━━━━━━━━━━━━━━━━━━━━━━━━━━━━━━━━━━ 47.6/47.6 kB 115.6 MB/s eta 0:00:00
   ━━━━━━━━━━━━━━━━━━━━━━━━━━━━━━━━━━━━━━━━ 2.2/2.2 MB 130.0 MB/s eta 0:00:00
   ━━━━━━━━━━━━━━━━━━━━━━━━━━━━━━━━━━━━━━━━ 756.0/756.0 kB 189.8 MB/s eta 0:00:00
   ━━━━━━━━━━━━━━━━━━━━━━━━━━━━━━━━━━━━━━━━ 1.6/1.6 MB 310.1 MB/s eta 0:00:00
  Building wheel for halo (setup.py) ... done
fhir_aggregator_client==0.1.8
```

#### Verify the tools were installed¶

```
!fq
```

```
Creating directory: /root/.fhir-aggregator
Usage: fq [OPTIONS] COMMAND [ARGS]...

  FHIR-Aggregator utilities.

Options:
  --help  Show this message and exit.

Commands:
  ls          List all the installed GraphDefinitions.
  run         Run GraphDefinition queries.
  results     Work with the results of a GraphDefinition query.
  vocabulary  FHIR-Aggregator's key Resources and CodeSystems.
```

#### Vocabulary Dataframe¶

As a convenience, the fhir-aggregator-client's vocabulary command will query this data and save it in a local dataframe.

This generates a tab-separated file named vocabulary.tsv, which serves as an inventory of the server's data elements and their usage, including example FHIR queries to retrieve those values.

```
!fq vocabulary vocabulary.tsv --fhir-base-url $FHIR_BASE
```

```
✔ Wrote 21846 vocabularies to vocabulary.tsv
```

This dataframe provides a catalog of the data elements present in the FHIR server, along with other useful information.

Create a dataframe from the vocabulary tsv. Note that the `url` field is a FHIR query that will return the resources that match that vocabulary. The `documentation` column links to the data dictionary documentation for that field.

| Column | Description |
|---|---|
| research_study_identifiers | A comma-separated list of identifiers that uniquely identify the research studies where this data element is found. This helps to link the data element back to specific studies. |
| path | The FHIR resource and element where the code is used (e.g., Condition.code, Observation.valueCodeableConcept). This shows the structural context of the data element within FHIR resources. |
| documentation | A URL linking to the official FHIR data dictionary documentation for the specific data element. This provides detailed information about the meaning and usage of the element according to the FHIR standard. |
| code | The actual code value within a CodeableConcept. This is the coded representation of the data element (e.g., a SNOMED CT code for a specific disease). |
| display | A human-readable label or description associated with the code. This makes it easier to understand the meaning of the code without needing to look up code system definitions. |
| system | The code system or terminology from which the code originates (e.g., http://snomed.info/sct, http://loinc.org). This helps to identify the source and context of the code. |
| extension_url | If the data element is an Extension, this column contains the URL that defines the Extension. Extensions provide a way to add custom data elements to FHIR resources. |
| count | The number of times this specific code value was found within the FHIR data. This gives an indication of the prevalence or frequency of the data element. |
| low | If the data element is numeric and has a defined range, this column represents the lower bound of the range. This is helpful for understanding the possible values for numeric data elements. |
| high | If the data element is numeric and has a defined range, this column represents the upper bound of the range. Similar to low, this helps to understand the potential values. |
| research_study_title | The title of the research study associated with this data element. This provides a more descriptive context for understanding where the data element is used. |
| research_study_description | A brief description of the research study associated with the data element. This offers additional context for the data element's usage. |
| observation | The ID of the Observation resource which is used to store the vocabulary. This provides a way to see where this data element was extracted from in the FHIR server. |
| research_study | The ID of the research study where the vocabulary is used. This allows you to retrieve the specific study that contains the vocabulary |
| url | A readily available FHIR query that can be used to retrieve the resources that contain the itemized data for this code. This makes it easy to access the data for analysis. |

```
import pandas as pd
df = pd.read_csv('vocabulary.tsv', sep='\t').fillna('')
df.loc[df['research_study_identifiers'] == 'GTEX_V10']
```

| | research_study_identifiers | path | documentation | code | display | system | extension_url | count | low | high | url | researc |
|---|---|---|---|---|---|---|---|---|---|---|---|---|
| **15** | GTEX_V10 | ServiceRequest.category | https://hl7.org/fhir/R4B/servicerequest-defini... | 108252007 | Laboratory procedure | http://snomed.info/sct | | 1.0 | | | https://google-fhir.fhir-aggregator.org/Servic... | GTEX Ana Adult Sar Subject M |
| **16** | GTEX_V10 | ServiceRequest.code | https://hl7.org/fhir/R4B/servicerequest-defini... | 15220000 | Laboratory test | http://snomed.info/sct | | 1.0 | | | https://google-fhir.fhir-aggregator.org/Servic... | GTEX Ana Adult Sar Subject M |
| **17** | GTEX_V10 | DocumentReference.type | https://hl7.org/fhir/R4B/documentreference-def... | file | file | https://gtexportal.org/api/v2/dataset/fileList | | 49.0 | | | https://google-fhir.fhir-aggregator.org/Docume... | GTEX Ana Adult Sar Subject M |
| **18** | GTEX_V10 | Specimen.type | https://hl7.org/fhir/R4B/specimen-definitions.... | WES | WES | https://terminology.hl7.org/CodeSystem-v3-Spec... | | 979.0 | | | https://google-fhir.fhir-aggregator.org/Specim... | GTEX Ana Adult Sar Subject M |
| **19** | GTEX_V10 | Specimen.type | https://hl7.org/fhir/R4B/specimen-definitions.... | OMNI | OMNI | https://terminology.hl7.org/CodeSystem-v3-Spec... | | 450.0 | | | https://google-fhir.fhir-aggregator.org/Specim... | GTEX Ana Adult Sar Subject M |
| **...** | ... | ... | ... | ... | ... | ... | ... | ... | ... | ... | ... | ... |
| **96** | GTEX_V10 | Patient.extension | https://hl7.org/fhir/R4B/patient-definitions.h... | 1956 - 1965 | 1956 - 1965 | | https://hl7.org/fhir/extensions/SearchParamete... | 5.0 | | | https://google-fhir.fhir-aggregator.org/Patien... | GTEX Ana Adult Sar Subject M |
| **97** | GTEX_V10 | Patient.extension | https://hl7.org/fhir/R4B/patient-definitions.h... | 1976 - 1985 | 1976 - 1985 | | https://hl7.org/fhir/extensions/SearchParamete... | 2.0 | | | https://google-fhir.fhir-aggregator.org/Patien... | GTEX Ana Adult Sar Subject M |
| **98** | GTEX_V10 | Patient.extension | https://hl7.org/fhir/R4B/patient-definitions.h... | 1966 - 1975 | 1966 - 1975 | | https://hl7.org/fhir/extensions/SearchParamete... | 7.0 | | | https://google-fhir.fhir-aggregator.org/Patien... | GTEX Ana Adult Sar Subject M |
| **99** | GTEX_V10 | Patient.extension | https://hl7.org/fhir/R4B/patient-definitions.h... | 1986 - 1995 | 1986 - 1995 | | https://hl7.org/fhir/extensions/SearchParamete... | 2.0 | | | https://google-fhir.fhir-aggregator.org/Patien... | GTEX Ana Adult Sar Subject M |
| **100** | GTEX_V10 | Patient.extension | https://hl7.org/fhir/R4B/patient-definitions.h... | 1946 - 1955 | 1946 - 1955 | | https://hl7.org/fhir/extensions/SearchParamete... | 2.0 | | | https://google-fhir.fhir-aggregator.org/Patien... | GTEX Ana Adult Sar Subject M |

86 rows × 15 columns

##### Accessing Native FHIR Data with Pre-formatted URLs¶

The vocabulary dataframe generated by the fq vocabulary command provides a convenient way to access the native FHIR data associated with each data element. This is achieved through the pre-formatted URLs present in the url column of the dataframe.

Here's how researchers can use these URLs:

**Identify the Data Element of Interest**

Researchers should first identify the specific data element they want to explore further. This can be done by examining the rows in the vocabulary dataframe and focusing on the columns like path, display, and system to locate the desired element.

**Retrieve the Pre-formatted URL**

Once the data element is identified, researchers should locate the corresponding pre-formatted URL in the url column of the same row. This URL is specifically crafted to retrieve FHIR resources containing the itemized data for that code.

**Execute the URL as a FHIR Query**

Researchers can execute this pre-formatted URL using tools like curl within a terminal or a Jupyter Notebook cell. The curl command will send a request to the FHIR server and retrieve the relevant FHIR resources. Here's how to execute the pre-formatted url with curl:

```
!curl -s $FHIR_BASE/$URL | jq .
```

Replace the following:

- `$FHIR_BASE` - FHIR server's base URL.
- `$URL` - The url obtained from the url field.

**Example**

Assume a researcher has found a pre-formatted url of DocumentReference/?category=H%26E&part-of-study=ResearchStudy/b7015858-983c-5e51-bcd6-af788edd5056 in the url column for H&E data:

`Note` Use the [documentation](documentation) field to see the FHIR data dictionary for the field

```
!curl -s $FHIR_BASE'/DocumentReference/?category=H%26E&part-of-study=ResearchStudy/b7015858-983c-5e51-bcd6-af788edd5056&_count=0' | echo There are
`jq .total` DocumentReferences of that category in this study.
```

```
There are 22 DocumentReferences of that category in this study.
```

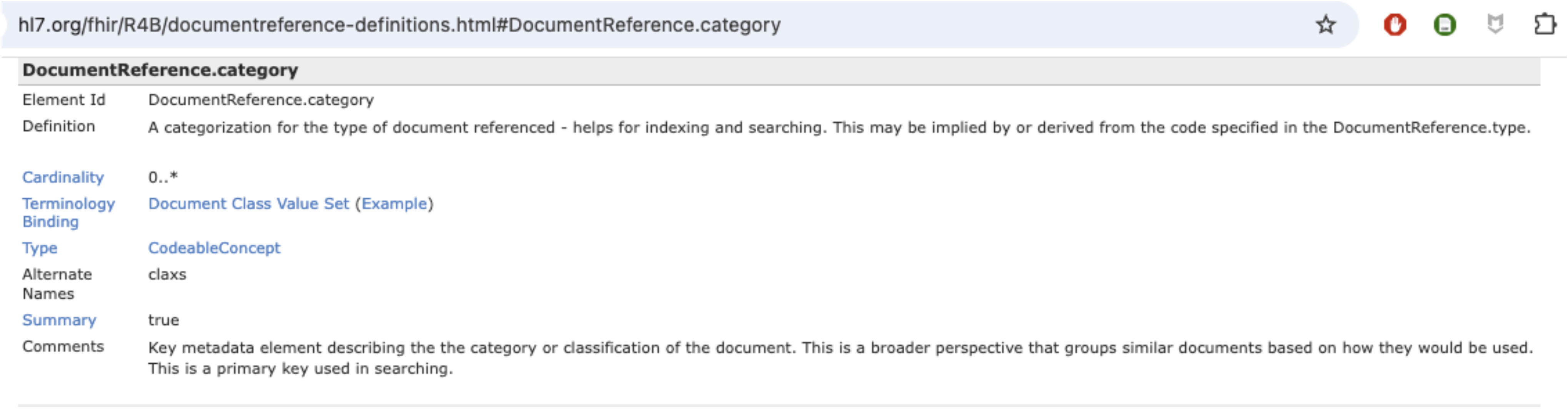
hl7.org/fhir/R4B/documentreference-definitions.html#DocumentReference.category

**DocumentReference.category**

| | |
|---|---|
| Element Id | DocumentReference.category |
| Definition | A categorization for the type of document referenced - helps for indexing and searching. This may be implied by or derived from the code specified in the DocumentReference.type. |
| Cardinality | 0..* |
| Terminology Binding | Document Class Value Set (Example) |
| Type | CodeableConcept |
| Alternate Names | claxs |
| Summary | true |
| Comments | Key metadata element describing the the category or classification of the document. This is a broader perspective that groups similar documents based on how they would be used. This is a primary key used in searching. |

#### Visualize the CodeableConcept used in each resource¶

FHIR CodeableConcepts are essential for representing and exchanging healthcare information using codes and terminologies. They ensure interoperability, provide semantic meaning, offer flexibility, and enhance human understanding of healthcare data.

```
combined_counts = df[['research_study_identifiers', 'path', 'system',  'display', 'count']]
# Filter combined_counts to ignore rows with 'extension' in 'path'
combined_counts = combined_counts[~combined_counts['path'].str.contains('extension')]

combined_counts
```

| | research_study_identifiers | path | system | display | count |
|---|---|---|---|---|---|
| **0** | ICGC-LUCA_KR | ServiceRequest.category | http://snomed.info/sct | Laboratory procedure | 15063.0 |
| **1** | ICGC-LUCA_KR | ServiceRequest.code | http://snomed.info/sct | Laboratory test | 15063.0 |
| **2** | ICGC-LUCA_KR | DocumentReference.type | https://aced-idp.org/ICGC-LUCA_KR | TGZ | 10120.0 |
| **3** | ICGC-LUCA_KR | DocumentReference.category | https://aced-idp.org/ICGC-LUCA_KR/data_type | Aligned Reads QC | 3265.0 |
| **4** | ICGC-LUCA_KR | DocumentReference.securityLabel | http://terminology.hl7.org/CodeSystem/v3-Confi... | normal | 15063.0 |
| **...** | ... | ... | ... | ... | ... |
| **21829** | HCMI-CMDC | MedicationAdministration.medicationCodeableCon... | https://gdc.cancer.gov/therapeutic_agents | Dabrafenib | 3.0 |
| **21830** | HCMI-CMDC | MedicationAdministration.medicationCodeableCon... | https://gdc.cancer.gov/therapeutic_agents | Monalizumab | 1.0 |
| **21831** | HCMI-CMDC | MedicationAdministration.medicationCodeableCon... | https://gdc.cancer.gov/therapeutic_agents | Andecaliximab | 1.0 |
| **21832** | HCMI-CMDC | MedicationAdministration.medicationCodeableCon... | https://gdc.cancer.gov/therapeutic_agents | Vinblastine | 2.0 |
| **21833** | HCMI-CMDC | MedicationAdministration.medicationCodeableCon... | https://gdc.cancer.gov/therapeutic_agents | Bavituximab | 1.0 |

20771 rows × 5 columns

```
import seaborn as sns
import matplotlib.pyplot as plt
import pandas as pd

# Create heatmap for each path


# Get unique paths
unique_paths = combined_counts['path'].unique()
combined_counts = combined_counts.infer_objects(copy=False)

# Create a heatmap for each path
for path in unique_paths:
    # Filter data for the current path
    filtered_data = combined_counts[combined_counts['path'] == path]

    # Pivot the filtered data for the heatmap
    heatmap_data = filtered_data.pivot_table(index='research_study_identifiers',
                                        columns=['display'],
                                        values='count',
                                        fill_value=0.0)

    # Create the heatmap
    plt.figure(figsize=(12, 10))  # Adjust figsize as needed

    # sns.heatmap(heatmap_data, annot=True, annot_kws={"size": 6}, fmt=".0f", cmap='viridis', cbar=False)
```

```
   # Create a mask to identify zero values
   mask = heatmap_data == 0

# Custom function to format annotations
   def heatmap_annot(val, **kwargs):
       if val == 0:  # Hide annotations for zero values
           return ""
       else:
           return f"{val:.0f}"  # Format non-zero values as integers


   # Generate heatmap with the custom annotation function
   ax = sns.heatmap(heatmap_data, annot=True, fmt=".0f", cmap='viridis',
               cbar_kws={'extend': 'both', 'extendrect': True},
               vmin=0,  # Set minimum value for color scale
               vmax=heatmap_data.max().max() / 2,  # Adjust maximum value
               annot_kws={"size": 6})  # Reduce font size


   # Apply the custom annotation function to each cell
   for text in ax.texts:
       text.set_text(heatmap_annot(float(text.get_text())))

   plt.title(f'Heatmap for Path: {path}')  # Set title with path name
   plt.xlabel('Display')
   plt.ylabel('Research Study Identifiers')
   plt.xticks(rotation=45, ha='right')
   plt.tight_layout()
   plt.show()
```

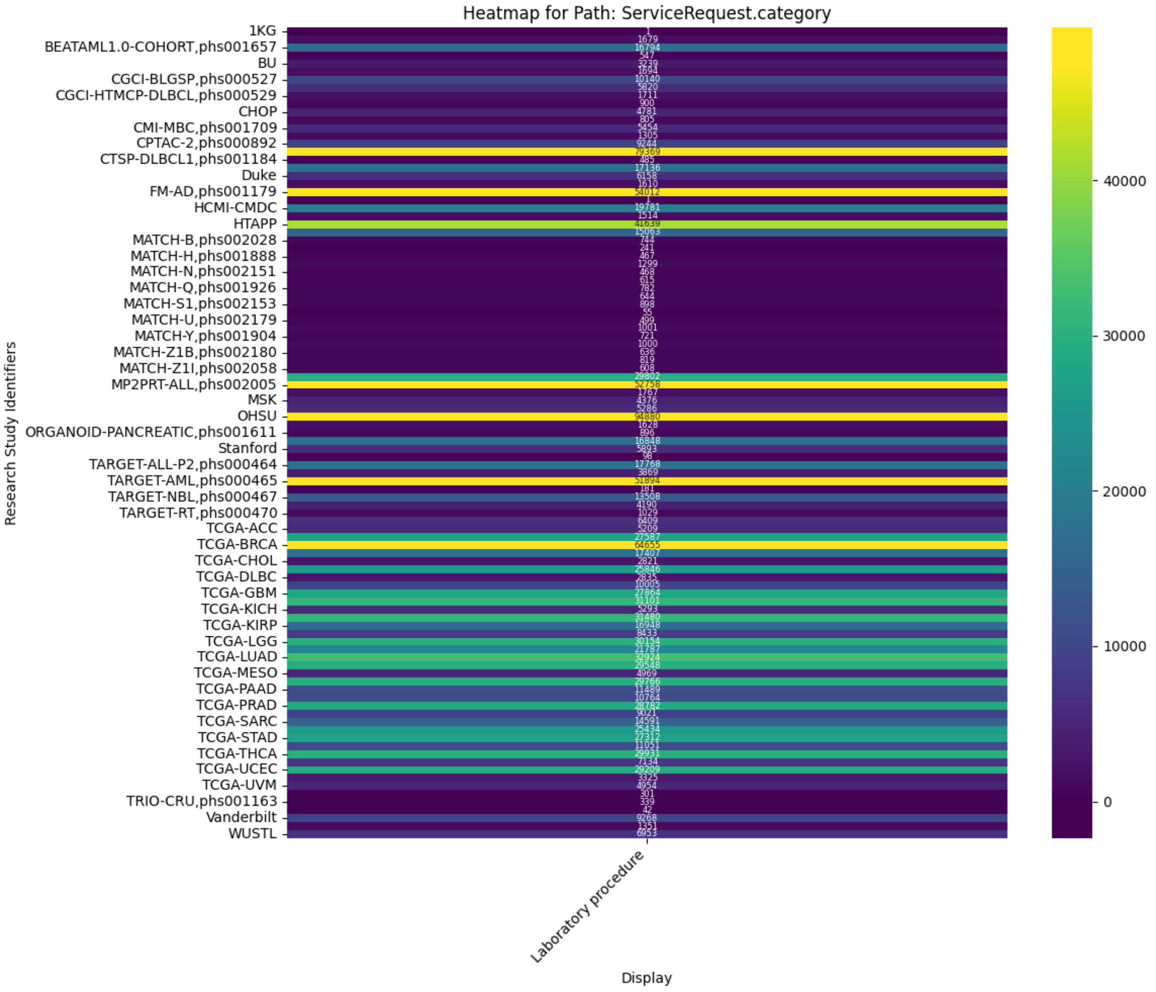
Heatmap for Path: ServiceRequest.category
Research Study Identifiers
Display
Laboratory procedure
1KG
BEATAML1.0-COHORT,phs001657
BU
CGCI-BLGSP,phs000527
CGCI-HTMCP-DLBCL,phs000529
CHOP
CMI-MBC,phs001709
CPTAC-2,phs000892
CTSP-DLBCL1,phs001184
Duke
FM-AD,phs001179
HCMI-CMDC
HTAPP
MATCH-B,phs002028
MATCH-H,phs001888
MATCH-N,phs002151
MATCH-Q,phs001926
MATCH-S1,phs002153
MATCH-U,phs002179
MATCH-Y,phs001904
MATCH-Z1B,phs002180
MATCH-Z1I,phs002058
MP2PRT-ALL,phs002005
MSK
OHSU
ORGANOID-PANCREATIC,phs001611
Stanford
TARGET-ALL-P2,phs000464
TARGET-AML,phs000465
TARGET-NBL,phs000467
TARGET-RT,phs000470
TCGA-ACC
TCGA-BRCA
TCGA-CHOL
TCGA-DLBC
TCGA-GBM
TCGA-KICH
TCGA-KIRP
TCGA-LGG
TCGA-LUAD
TCGA-MESO
TCGA-PAAD
TCGA-PRAD
TCGA-SARC
TCGA-STAD
TCGA-THCA
TCGA-UCEC
TCGA-UVM
TRIO-CRU,phs001163
Vanderbilt
WUSTL
40000
30000
20000
10000
0

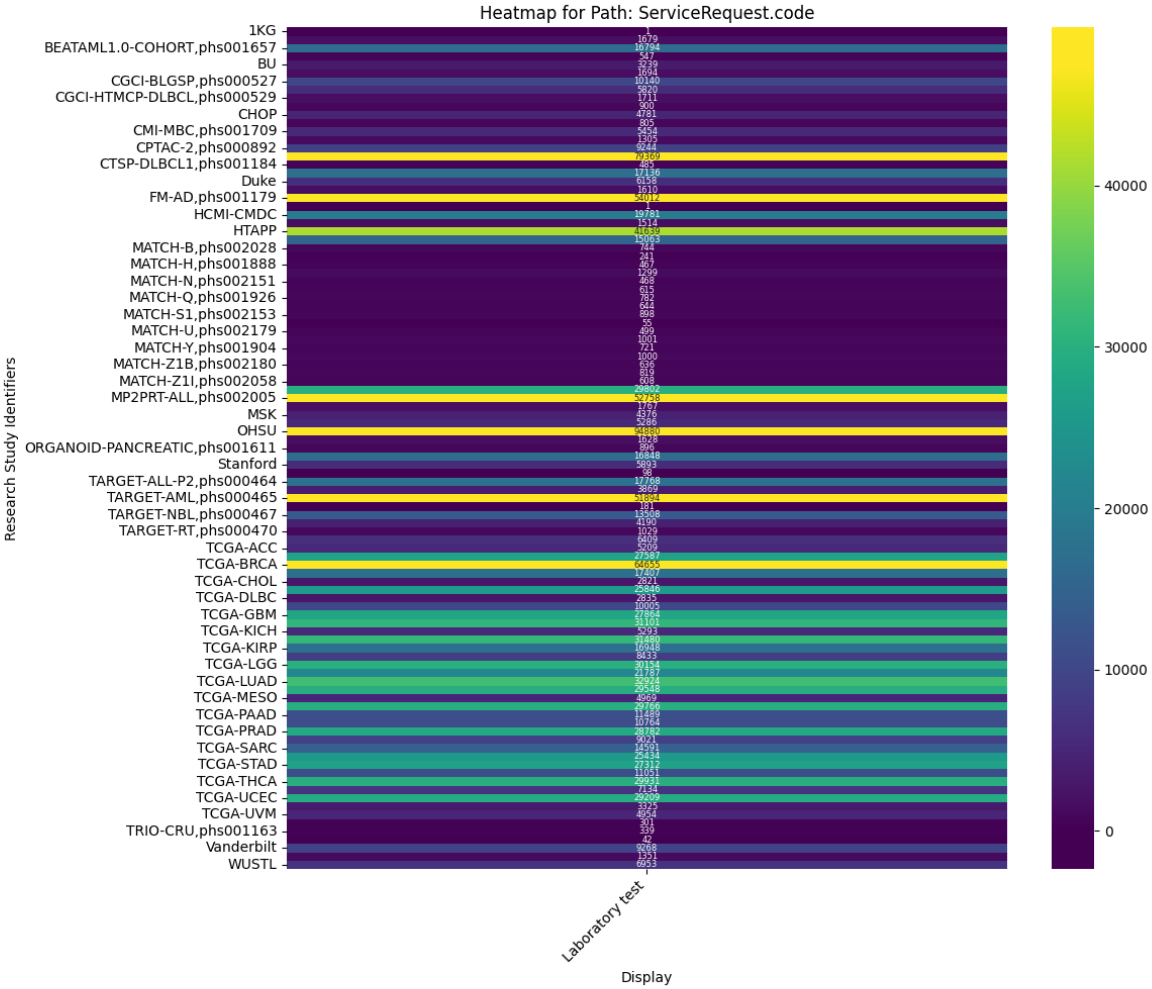
Heatmap for Path: ServiceRequest.code
Research Study Identifiers
1KG
BEATAML1.0-COHORT,phs001657
BU
CGCI-BLGSP,phs000527
CGCI-HTMCP-DLBCL,phs000529
CHOP
CMI-MBC,phs001709
CPTAC-2,phs000892
CTSP-DLBCL1,phs001184
Duke
FM-AD,phs001179
HCMI-CMDC
HTAPP
MATCH-B,phs002028
MATCH-H,phs001888
MATCH-N,phs002151
MATCH-Q,phs001926
MATCH-S1,phs002153
MATCH-U,phs002179
MATCH-Y,phs001904
MATCH-Z1B,phs002180
MATCH-Z1I,phs002058
MP2PRT-ALL,phs002005
MSK
OHSU
ORGANOID-PANCREATIC,phs001611
Stanford
TARGET-ALL-P2,phs000464
TARGET-AML,phs000465
TARGET-NBL,phs000467
TARGET-RT,phs000470
TCGA-ACC
TCGA-BRCA
TCGA-CHOL
TCGA-DLBC
TCGA-GBM
TCGA-KICH
TCGA-KIRP
TCGA-LGG
TCGA-LUAD
TCGA-MESO
TCGA-PAAD
TCGA-PRAD
TCGA-SARC
TCGA-STAD
TCGA-THCA
TCGA-UCEC
TCGA-UVM
TRIO-CRU,phs001163
Vanderbilt
WUSTL
Laboratory test
Display
40000
30000
20000
10000
0

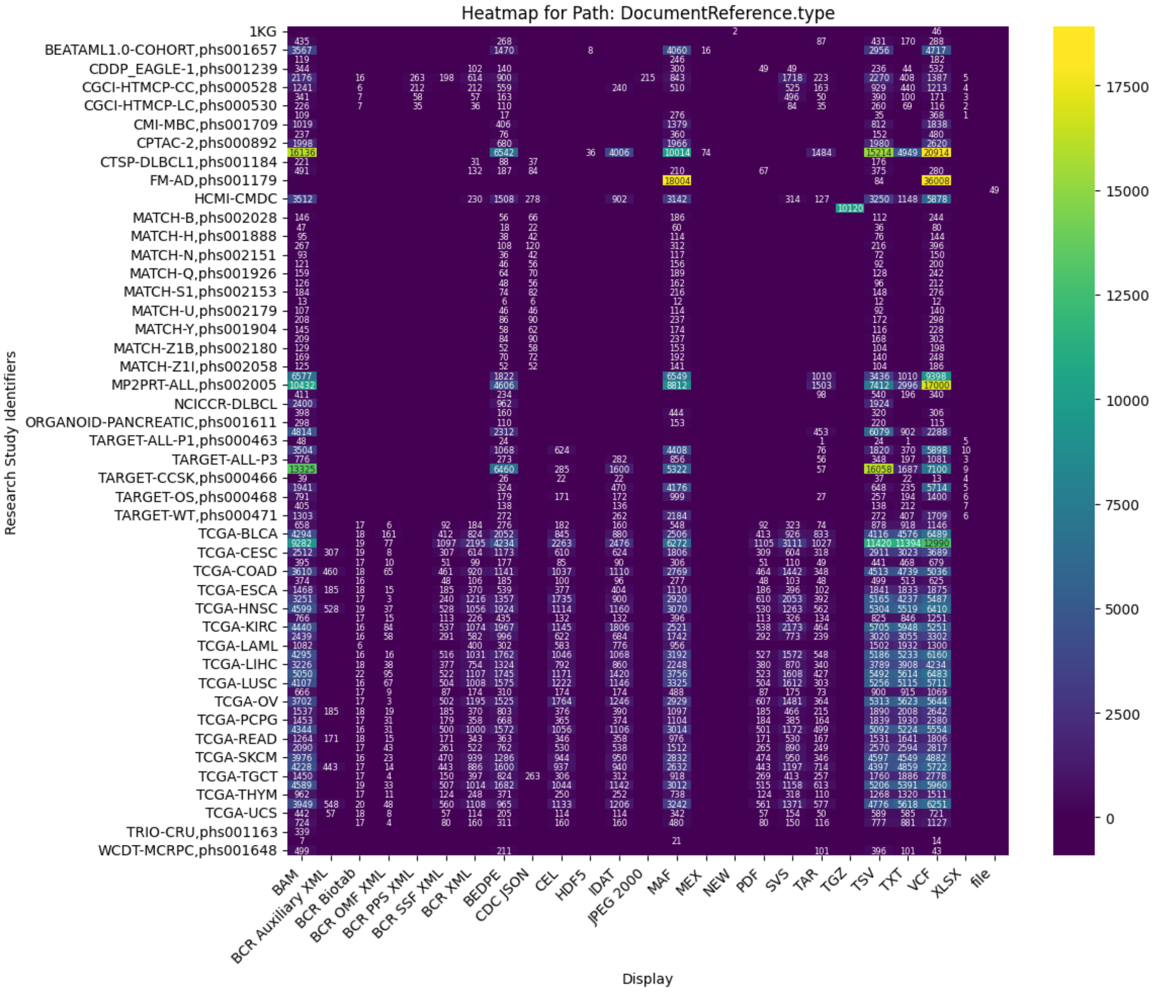

Heatmap for Path: DocumentReference.type
Research Study Identifiers
1KG
BEATAML1.0-COHORT,phs001657
CDDP_EAGLE-1,phs001239
CGCI-HTMCP-CC,phs000528
CGCI-HTMCP-LC,phs000530
CMI-MBC,phs001709
CPTAC-2,phs000892
CTSP-DLBCL1,phs001184
FM-AD,phs001179
HCMI-CMDC
MATCH-B,phs002028
MATCH-H,phs001888
MATCH-N,phs002151
MATCH-Q,phs001926
MATCH-S1,phs002153
MATCH-U,phs002179
MATCH-Y,phs001904
MATCH-Z1B,phs002180
MATCH-Z1I,phs002058
MP2PRT-ALL,phs002005
NCICCR-DLBCL
ORGANOID-PANCREATIC,phs001611
TARGET-ALL-P1,phs000463
TARGET-ALL-P3
TARGET-CCSK,phs000466
TARGET-OS,phs000468
TARGET-WT,phs000471
TCGA-BLCA
TCGA-CESC
TCGA-COAD
TCGA-ESCA
TCGA-HNSC
TCGA-KIRC
TCGA-LAML
TCGA-LIHC
TCGA-LUSC
TCGA-OV
TCGA-PCPG
TCGA-READ
TCGA-SKCM
TCGA-TGCT
TCGA-THYM
TCGA-UCS
TRIO-CRU,phs001163
WCDT-MCRPC,phs001648
BAM
BCR Auxiliary XML
BCR Biotab
BCR OMF XML
BCR PPS XML
BCR SSF XML
BCR XML
BEDPE
CDC JSON
CEL
HDF5
IDAT
JPEG 2000
MAF
MEX
NEW
PDF
SVS
TAR
TGZ
TSV
TXT
VCF
XLSX
file
Display
17500
15000
12500
10000
7500
5000
2500
0

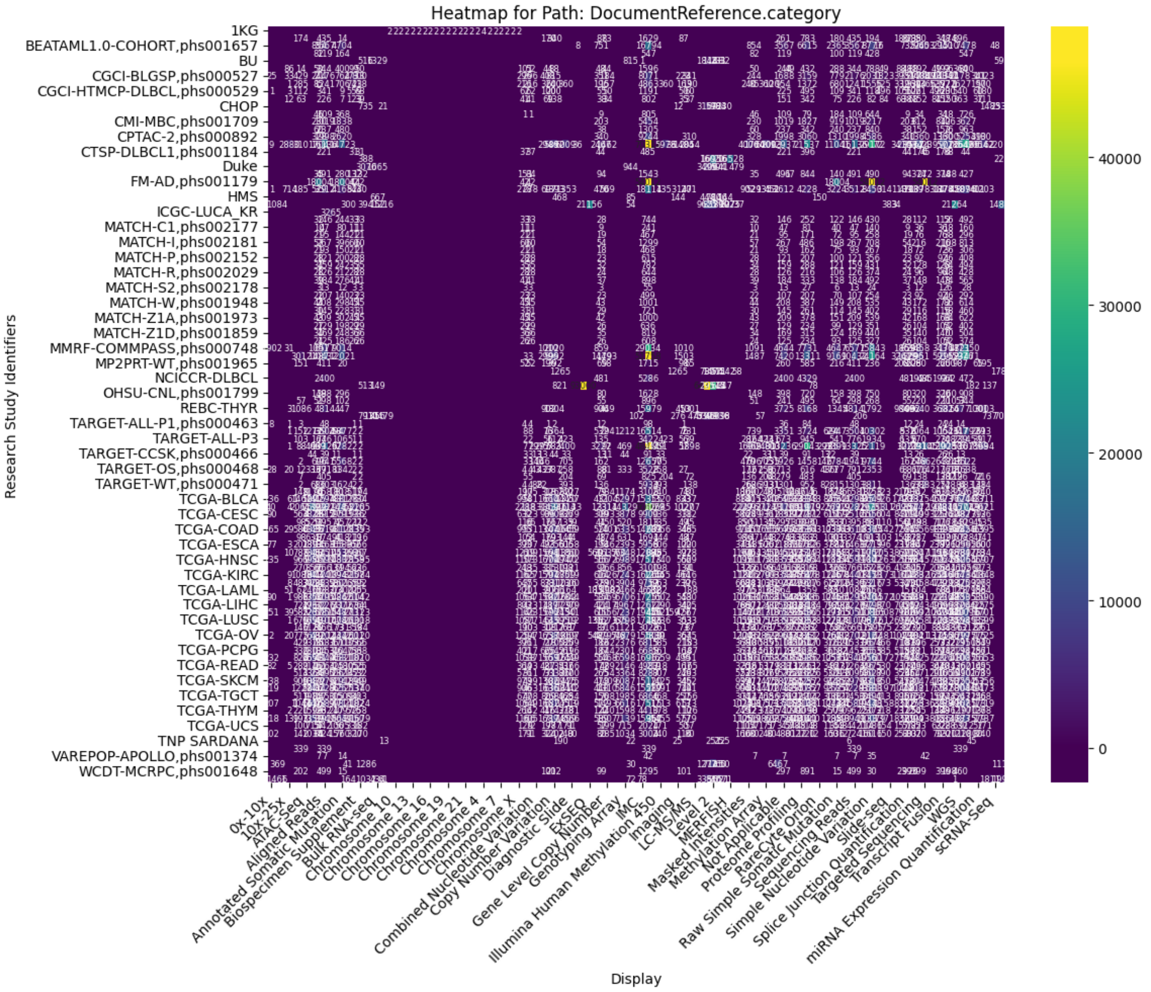
Heatmap for Path: DocumentReference.category
Research Study Identifiers
Display
1KG
BEATAML1.0-COHORT,phs001657
BU
CGCI-BLGSP,phs000527
CGCI-HTMCP-DLBCL,phs000529
CHOP
CMI-MBC,phs001709
CPTAC-2,phs000892
CTSP-DLBCL1,phs001184
Duke
FM-AD,phs001179
HMS
ICGC-LUCA_KR
MATCH-C1,phs002177
MATCH-I,phs002181
MATCH-P,phs002152
MATCH-R,phs002029
MATCH-S2,phs002178
MATCH-W,phs001948
MATCH-Z1A,phs001973
MATCH-Z1D,phs001859
MMRF-COMMPASS,phs000748
MP2PRT-WT,phs001965
NCICCR-DLBCL
OHSU-CNL,phs001799
REBC-THYR
TARGET-ALL-P1,phs000463
TARGET-ALL-P3
TARGET-CCSK,phs000466
TARGET-OS,phs000468
TARGET-WT,phs000471
TCGA-BLCA
TCGA-CESC
TCGA-COAD
TCGA-ESCA
TCGA-HNSC
TCGA-KIRC
TCGA-LAML
TCGA-LIHC
TCGA-LUSC
TCGA-OV
TCGA-PCPG
TCGA-READ
TCGA-SKCM
TCGA-TGCT
TCGA-THYM
TCGA-UCS
TNP SARDANA
VAREPOP-APOLLO,phs001374
WCDT-MCRPC,phs001648
0x-10x
10x-25x
ATAC-Seq
Aligned Reads
Annotated Somatic Mutation
Biospecimen Supplement
Bulk RNA-seq
Chromosome 10
Chromosome 13
Chromosome 16
Chromosome 19
Chromosome 21
Chromosome 4
Chromosome 7
Chromosome X
Combined Nucleotide Variation
Copy Number Variation
Diagnostic Slide
EXSEQ
Gene Level Copy Number
Genotyping Array
IMC
Illumina Human Methylation 450
Imaging
LC-MS/MS
Level 2
MERFISH
Masked Intensities
Methylation Array
Not Applicable
Proteome Profiling
RareCyte Orion
Raw Simple Somatic Mutation
Sequencing Reads
Simple Nucleotide Variation
Slide-seq
Splice Junction Quantification
Targeted Sequencing
Transcript Fusion
WGS
miRNA Expression Quantification
scRNA-Seq
40000
30000
20000
10000
0

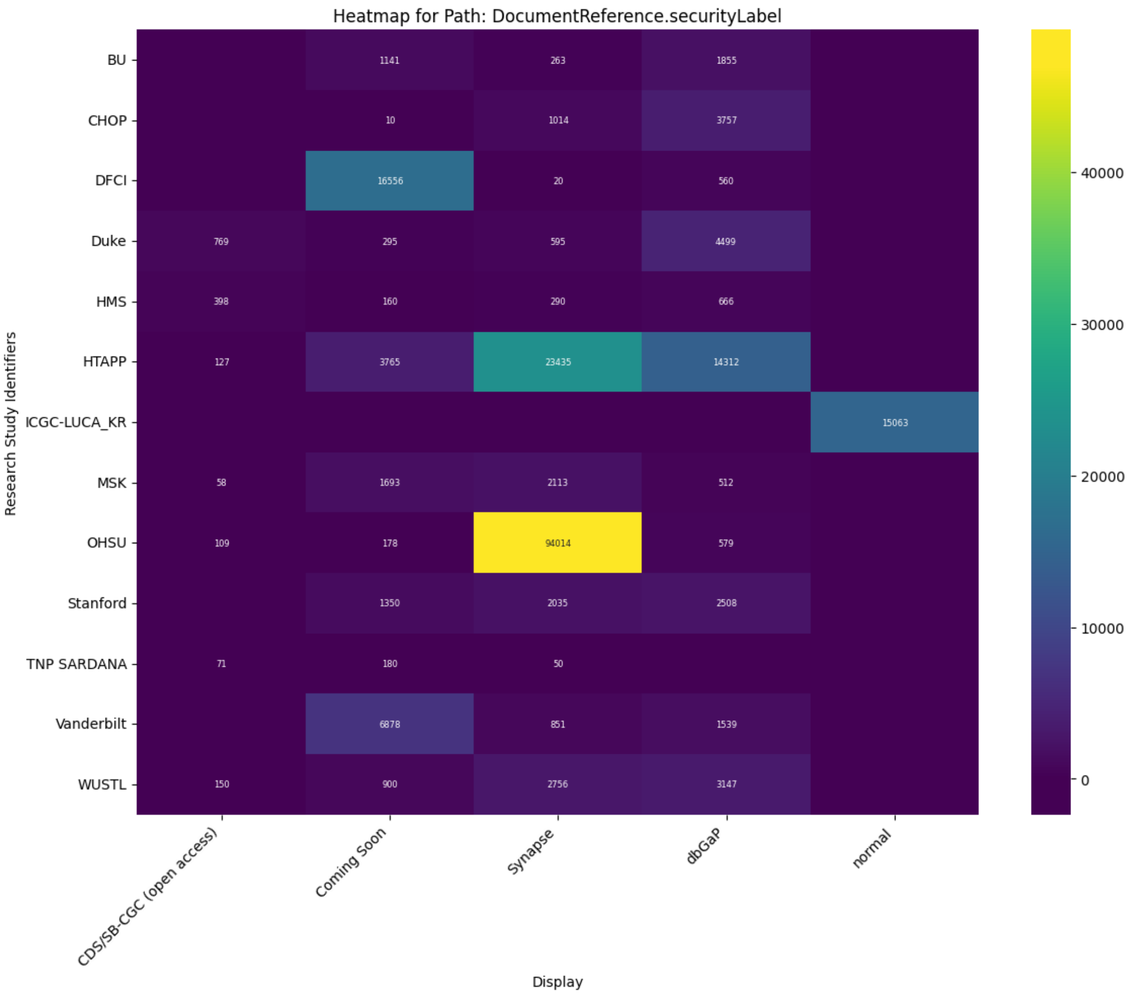
Heatmap for Path: DocumentReference.securityLabel
Research Study Identifiers
BU
CHOP
DFCI
Duke
HMS
HTAPP
ICGC-LUCA_KR
MSK
OHSU
Stanford
TNP SARDANA
Vanderbilt
WUSTL
1141
263
1855
10
1014
3757
16556
20
560
769
295
595
4499
398
160
290
666
127
3765
23435
14312
15063
58
1693
2113
512
109
178
94014
579
1350
2035
2508
71
180
50
6878
851
1539
150
900
2756
3147
CDS/SB-CGC (open access)
Coming Soon
Synapse
dbGaP
normal
Display
40000
30000
20000
10000
0

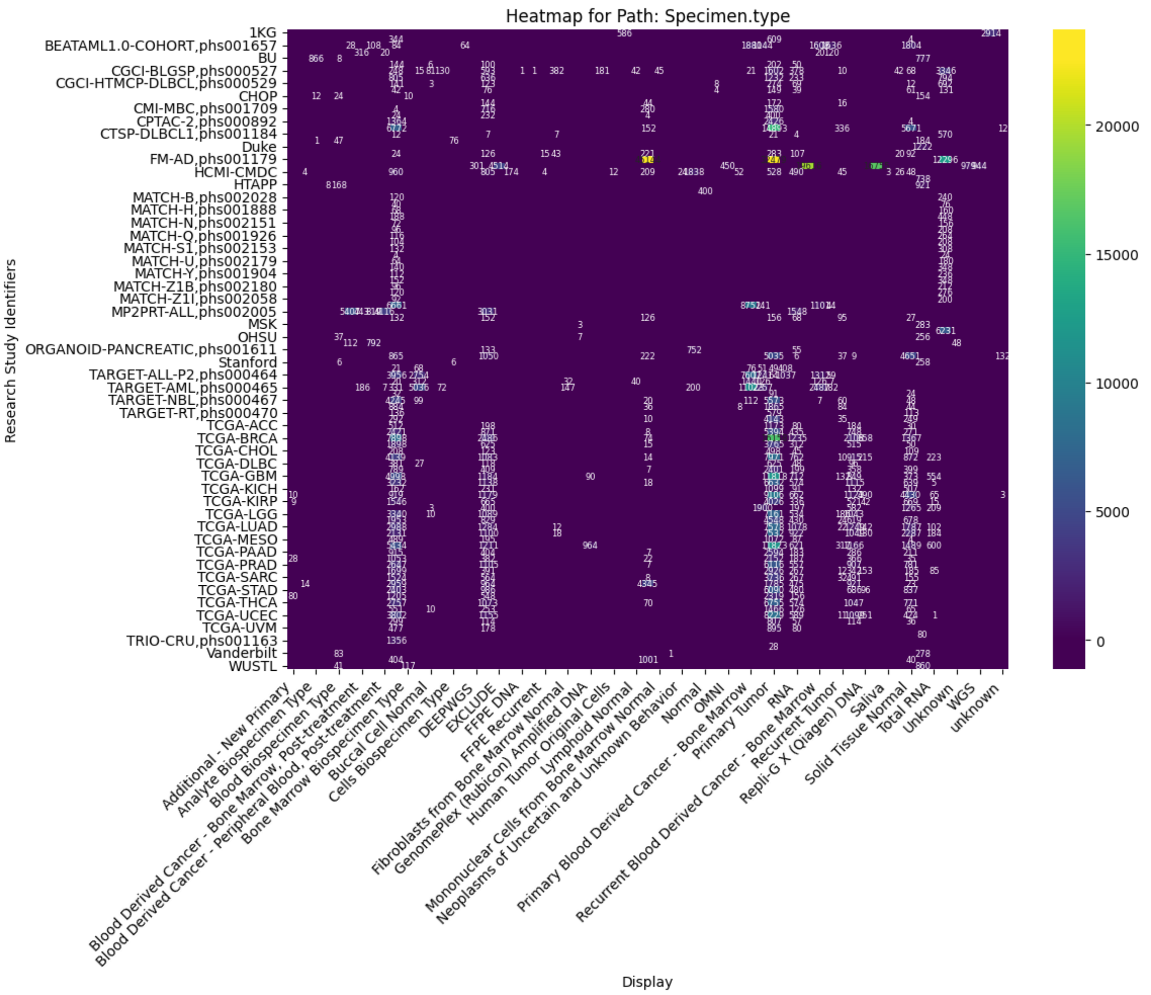

Heatmap for Path: Specimen.type
Research Study Identifiers
1KG
BEATAML1.0-COHORT,phs001657
BU
CGCI-BLGSP,phs000527
CGCI-HTMCP-DLBCL,phs000529
CHOP
CMI-MBC,phs001709
CPTAC-2,phs000892
CTSP-DLBCL1,phs001184
Duke
FM-AD,phs001179
HCMI-CMDC
HTAPP
MATCH-B,phs002028
MATCH-H,phs001888
MATCH-N,phs002151
MATCH-Q,phs001926
MATCH-S1,phs002153
MATCH-U,phs002179
MATCH-Y,phs001904
MATCH-Z1B,phs002180
MATCH-Z1I,phs002058
MP2PRT-ALL,phs002005
MSK
OHSU
ORGANOID-PANCREATIC,phs001611
Stanford
TARGET-ALL-P2,phs000464
TARGET-AML,phs000465
TARGET-NBL,phs000467
TARGET-RT,phs000470
TCGA-ACC
TCGA-BRCA
TCGA-CHOL
TCGA-DLBC
TCGA-GBM
TCGA-KICH
TCGA-KIRP
TCGA-LGG
TCGA-LUAD
TCGA-MESO
TCGA-PAAD
TCGA-PRAD
TCGA-SARC
TCGA-STAD
TCGA-THCA
TCGA-UCEC
TCGA-UVM
TRIO-CRU,phs001163
Vanderbilt
WUSTL
Additional - New Primary
Analyte Biospecimen Type
Blood Biospecimen Type
Blood Derived Cancer - Bone Marrow, Post-treatment
Blood Derived Cancer - Peripheral Blood, Post-treatment
Bone Marrow Biospecimen Type
Buccal Cell Normal
Cells Biospecimen Type
DEEPWGS
EXCLUDE
FFPE DNA
FFPE Recurrent
Fibroblasts from Bone Marrow Normal
GenomePlex (Rubicon) Amplified DNA
Human Tumor Original Cells
Lymphoid Normal
Mononuclear Cells from Bone Marrow Normal
Neoplasms of Uncertain and Unknown Behavior
Normal
OMNI
Primary Blood Derived Cancer - Bone Marrow
Primary Tumor
RNA
Recurrent Blood Derived Cancer - Bone Marrow
Recurrent Tumor
Repli-G X (Qiagen) DNA
Saliva
Solid Tissue Normal
Total RNA
Unknown
WGS
unknown
Display
20000
15000
10000
5000
0

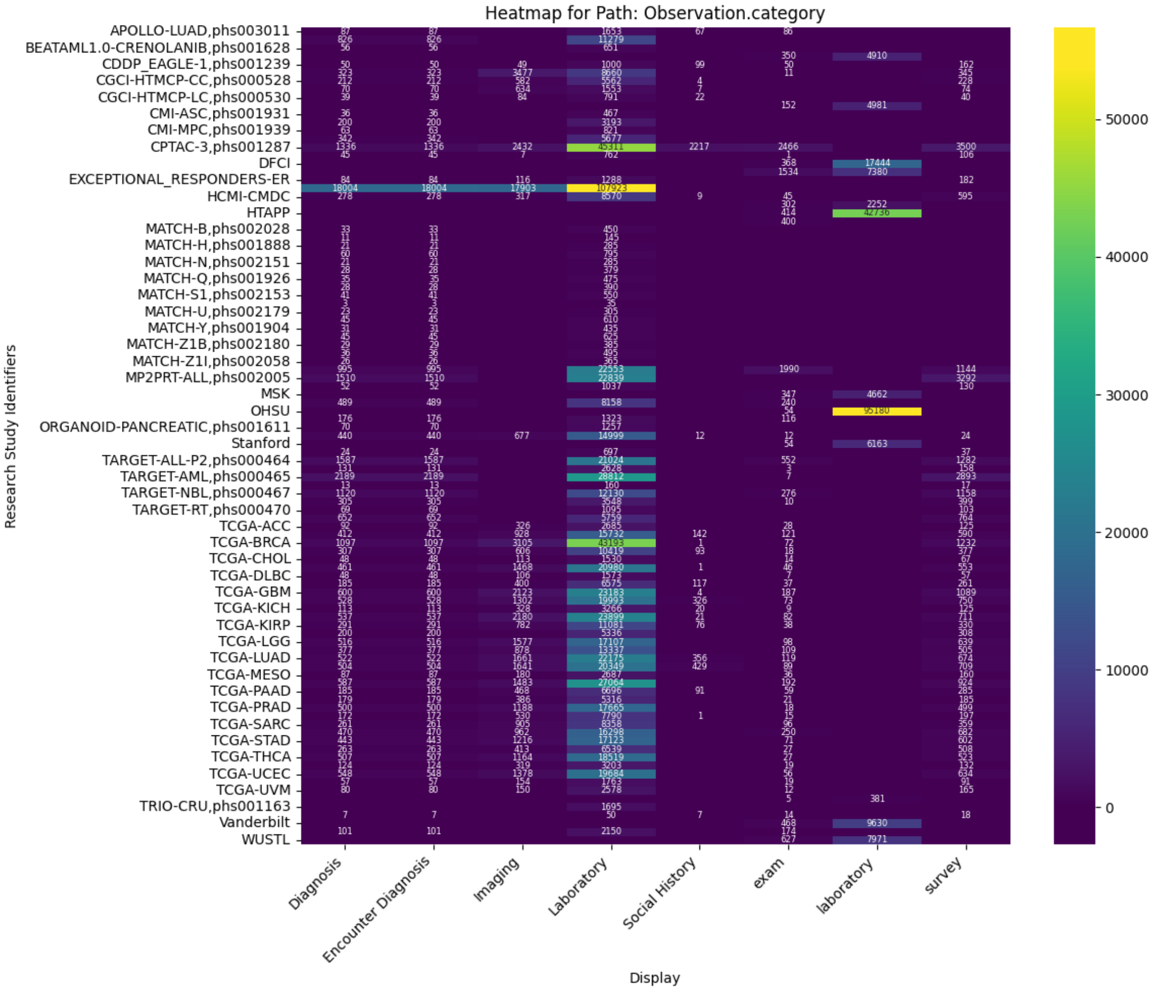
Heatmap for Path: Observation.category
Research Study Identifiers
Display
Diagnosis
Encounter Diagnosis
Imaging
Laboratory
Social History
exam
laboratory
survey
50000
40000
30000
20000
10000
0

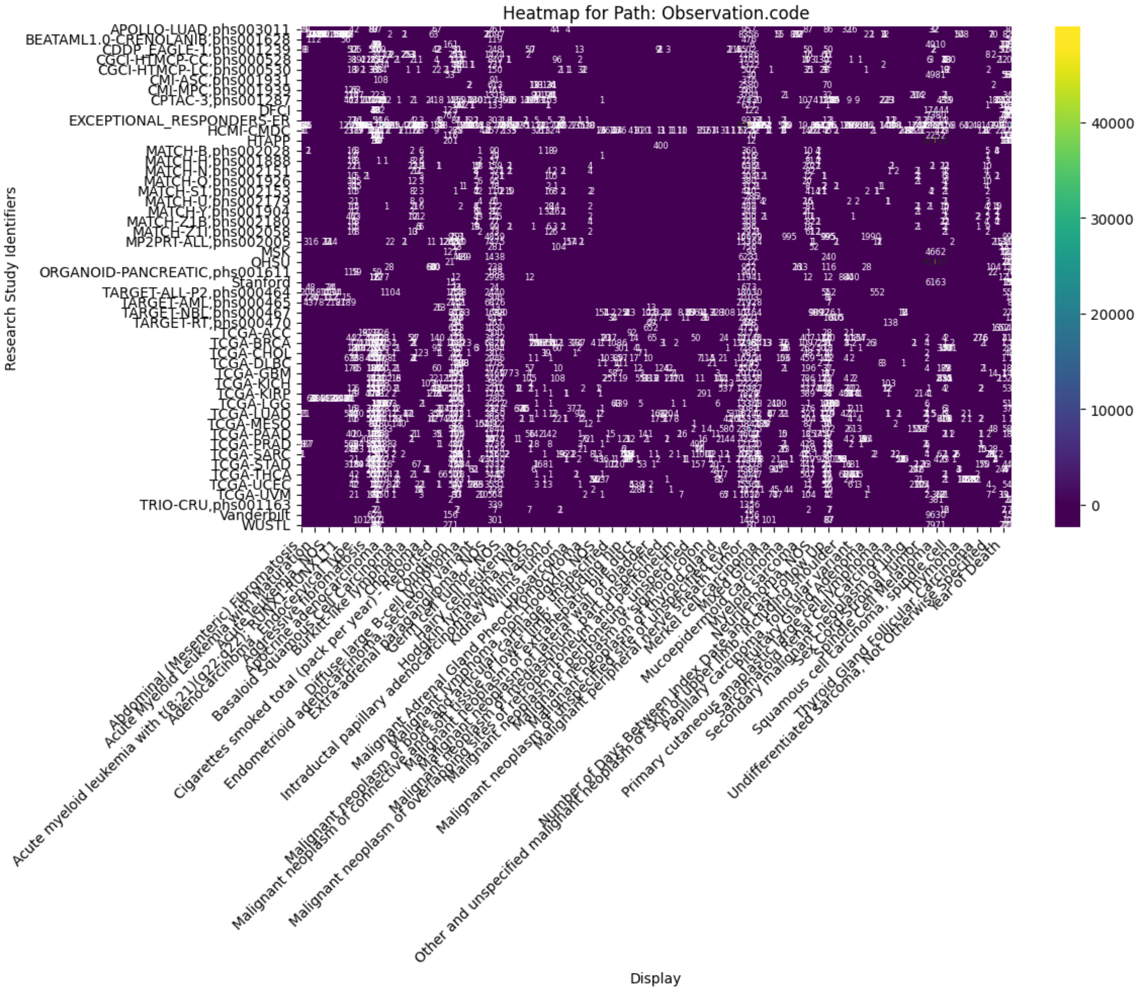
Heatmap for Path: Observation.code
Research Study Identifiers
Display
APOLLO-LUAD,phs003011
BEATAML1.0-CRENOLANIB,phs001628
CPTAC-3,phs001287
DFCI
EXCEPTIONAL_RESPONDERS-ER
HCMI-CMDC
HTAPP
MP2PRT-ALL,phs002005
MSK
OHSU
ORGANOID-PANCREATIC,phs001611
Stanford
TARGET-ALL-P2,phs000464
TARGET-RT,phs000470
TCGA-ACC
TCGA-BRCA
TCGA-CHOL
TCGA-DLBC
TCGA-GBM
TCGA-KICH
TCGA-KIRP
TCGA-LGG
TCGA-LUAD
TCGA-MESO
TCGA-PAAD
TCGA-PRAD
TCGA-SARC
TCGA-STAD
TCGA-THCA
TCGA-UCEC
TCGA-UVM
TRIO-CRU,phs001163
Vanderbilt
WUSTL
40000
30000
20000
10000
0
Abdominal (Mesenteric) Fibromatosis
Cigarettes smoked total (pack per year)
Number of Days Between Index Date and Diagnosis
Other and unspecified malignant neoplasm of skin of upper limb, including shoulder
Undifferentiated Sarcoma, Not Otherwise Specified
Year of Death

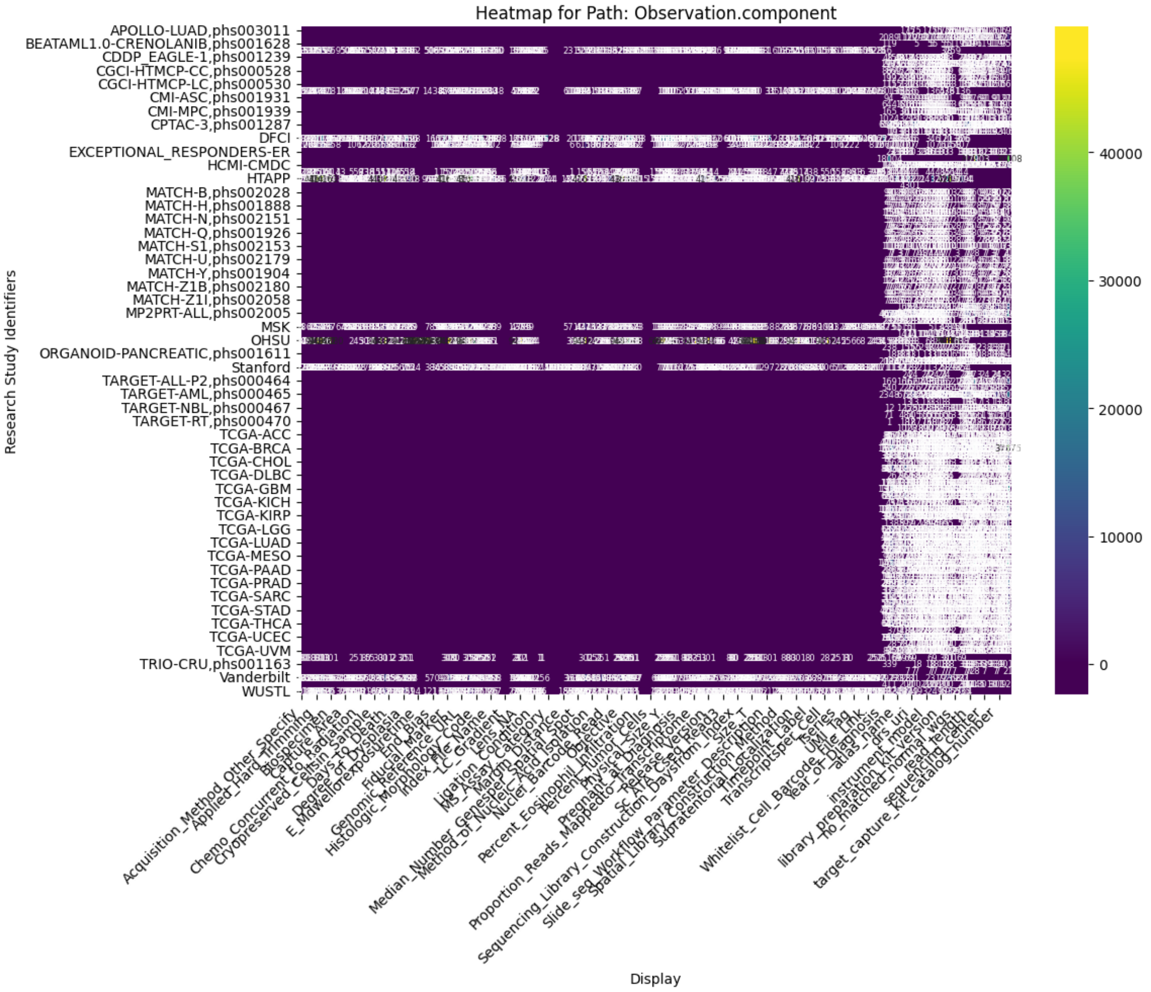

Heatmap for Path: Observation.component
Research Study Identifiers
APOLLO-LUAD,phs003011
BEATAML1.0-CRENOLANIB,phs001628
CDDP_EAGLE-1,phs001239
CGCI-HTMCP-CC,phs000528
CGCI-HTMCP-LC,phs000530
CMI-ASC,phs001931
CMI-MPC,phs001939
CPTAC-3,phs001287
DFCI
EXCEPTIONAL_RESPONDERS-ER
HCMI-CMDC
HTAPP
MATCH-B,phs002028
MATCH-H,phs001888
MATCH-N,phs002151
MATCH-Q,phs001926
MATCH-S1,phs002153
MATCH-U,phs002179
MATCH-Y,phs001904
MATCH-Z1B,phs002180
MATCH-Z1I,phs002058
MP2PRT-ALL,phs002005
MSK
OHSU
ORGANOID-PANCREATIC,phs001611
Stanford
TARGET-ALL-P2,phs000464
TARGET-AML,phs000465
TARGET-NBL,phs000467
TARGET-RT,phs000470
TCGA-ACC
TCGA-BRCA
TCGA-CHOL
TCGA-DLBC
TCGA-GBM
TCGA-KICH
TCGA-KIRP
TCGA-LGG
TCGA-LUAD
TCGA-MESO
TCGA-PAAD
TCGA-PRAD
TCGA-SARC
TCGA-STAD
TCGA-THCA
TCGA-UCEC
TCGA-UVM
TRIO-CRU,phs001163
Vanderbilt
WUSTL
Acquisition_Method_Other_Specify
Applied_Hard_Trimming
Biospecimen
Capture_Area
Chemo_Concurrent_to_Radiation
Cryopreserved_Cells_in_Sample
Days_to_Death
Degree_of_Dysplasia
Histologic_Morphology_Code
Index_File_Name
LC_Gradient
Lens_NA
Ligation
MS_Assay_Category
Median_Number_Genes_per_Spatial_Spot
Method_of_Nucleic_Acid_Isolation
Nuclei_Barcode_Read
Percent_Eosinophil_Infiltration
Percent_Tumor_Cells
Pregnant_at_Diagnosis
Proportion_Reads_Mapped_to_Transcriptome
Sequencing_Library_Construction_Days_from_Index_T
Slide_seq_Workflow_Parameter_Description
Spatial_Library_Construction_Method
Supratentorial_Localization
Timepoint_Label
Whitelist_Cell_Barcode_File_Link
Year_of_Diagnosis
atlas_name
drs_uri
instrument_model
library_preparation_kit_version
no_matched_normal_wgs
read_length
sequencing_center
target_capture_kit_catalog_number
Display
40000
30000
20000
10000
0

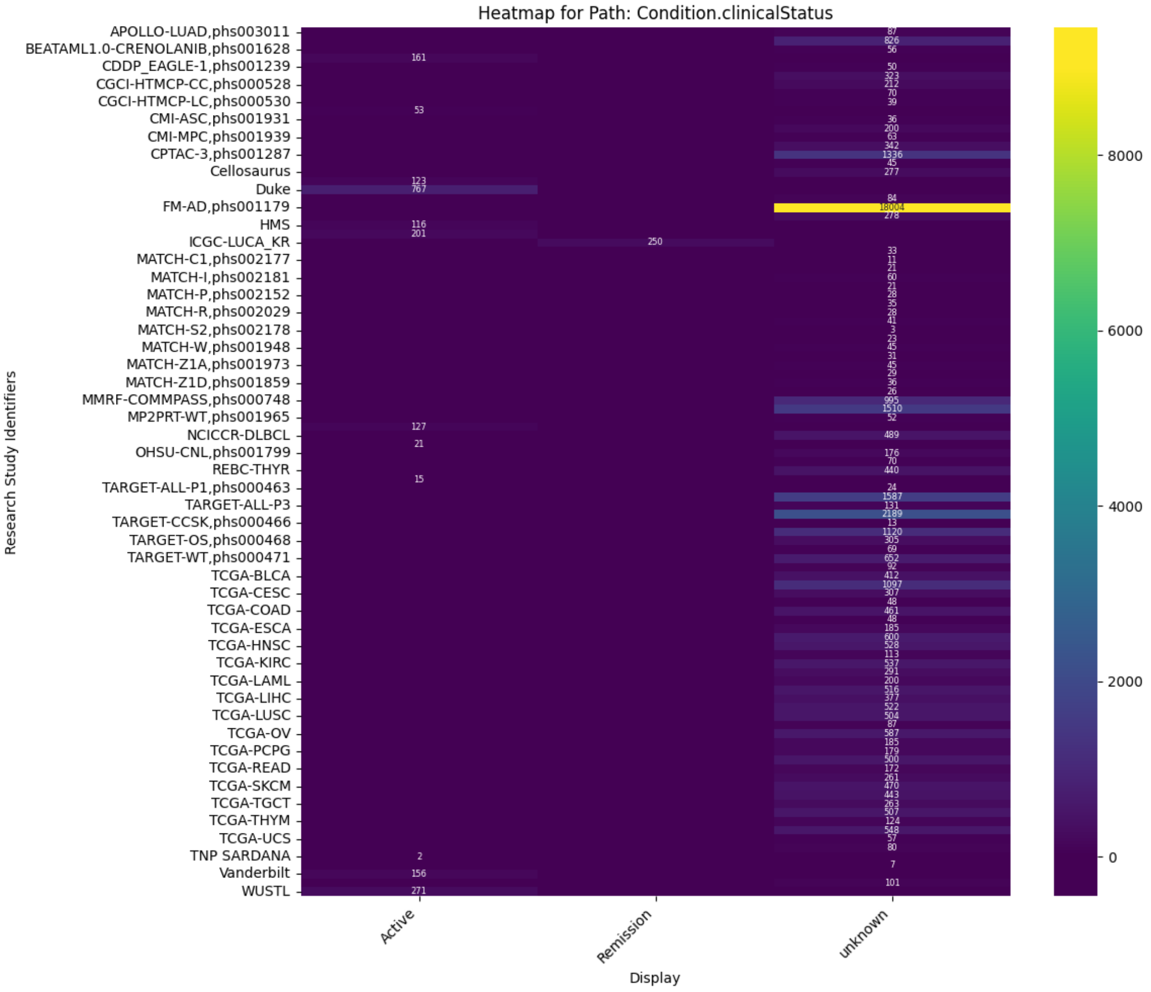
Heatmap for Path: Condition.clinicalStatus
Research Study Identifiers
Display
Active
Remission
unknown

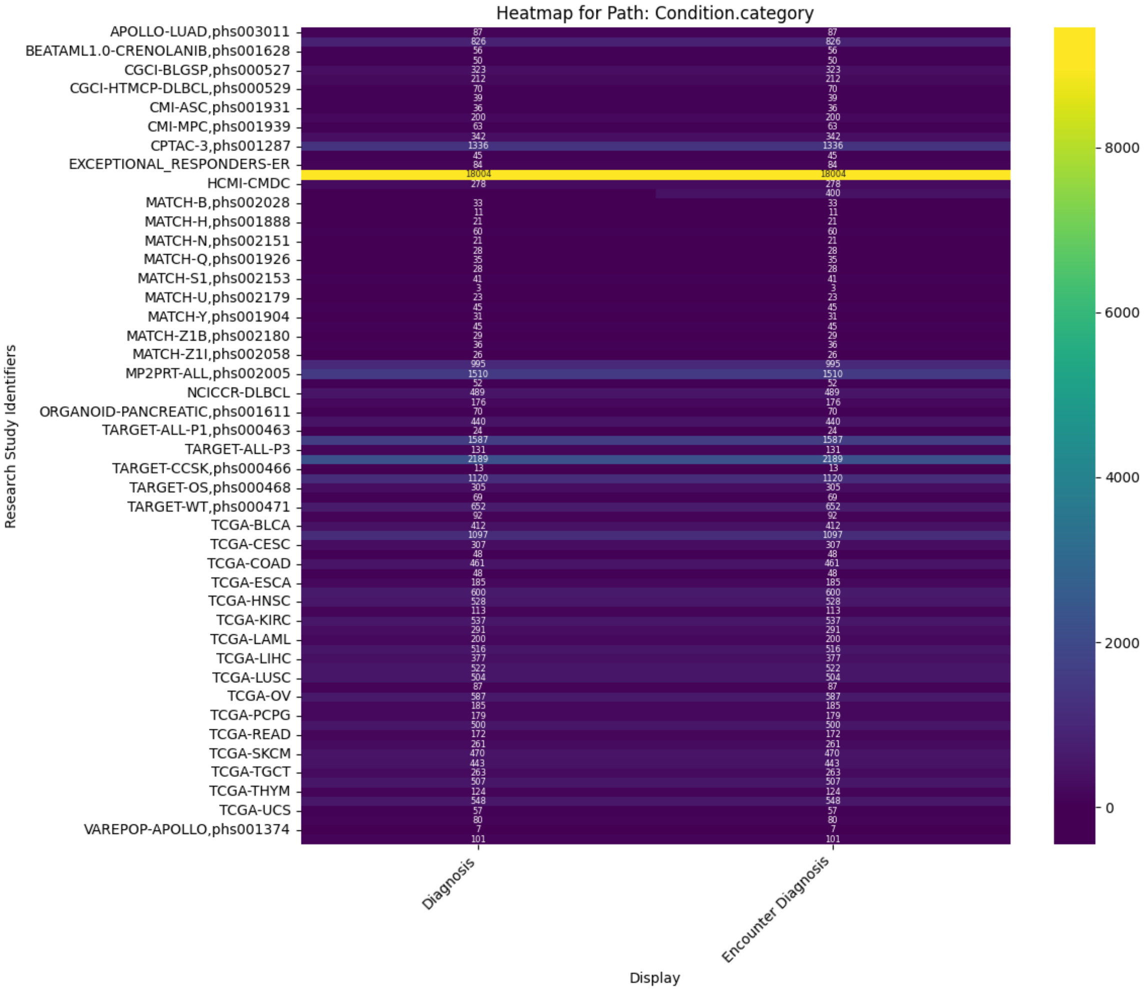
Heatmap for Path: Condition.category
Research Study Identifiers
Display
Diagnosis
Encounter Diagnosis

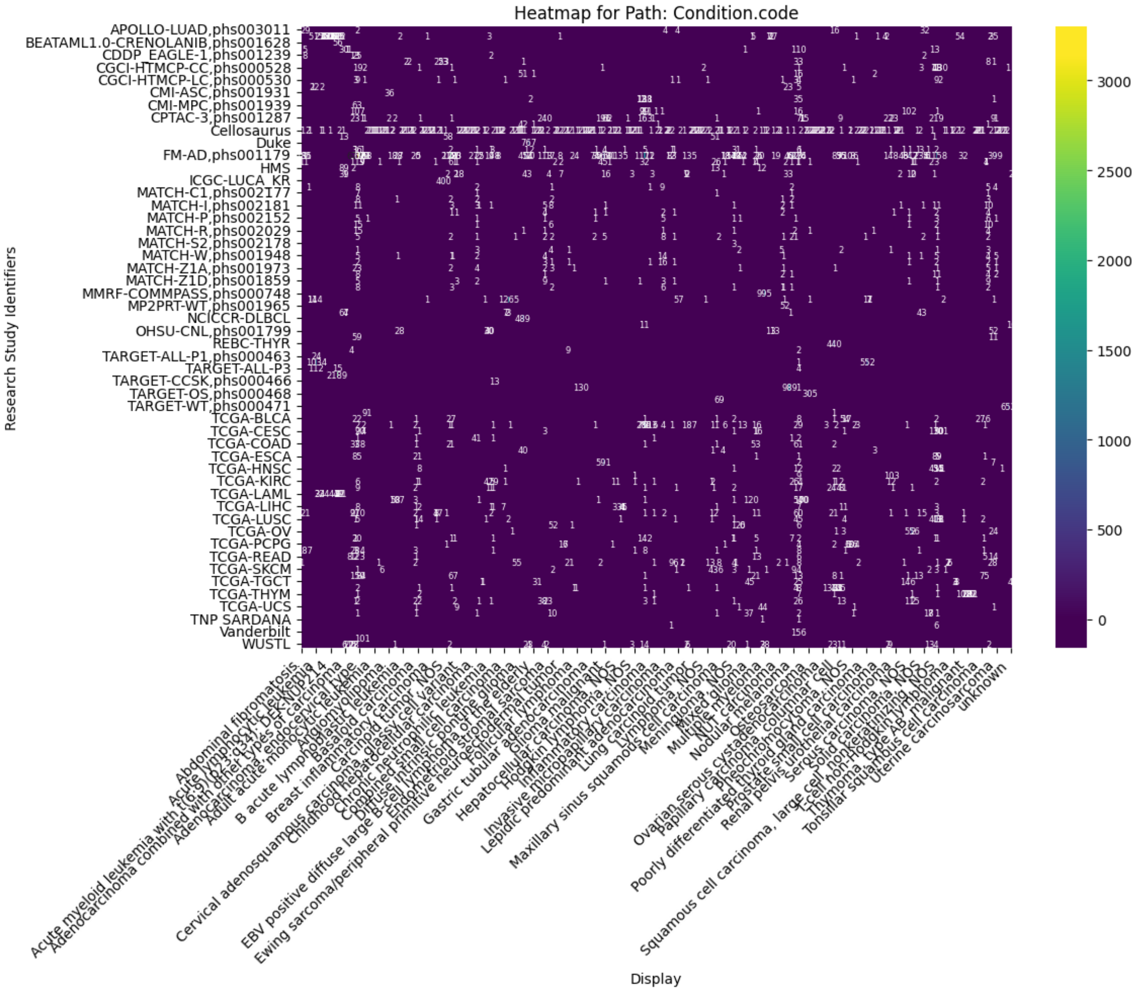
Heatmap for Path: Condition.code
APOLLO-LUAD,phs003011
BEATAML1.0-CRENOLANIB,phs001628
CDDP_EAGLE-1,phs001239
CGCI-HTMCP-CC,phs000528
CGCI-HTMCP-LC,phs000530
CMI-ASC,phs001931
CMI-MPC,phs001939
CPTAC-3,phs001287
Cellosaurus
Duke
FM-AD,phs001179
HMS
ICGC-LUCA_KR
MATCH-C1,phs002177
MATCH-I,phs002181
MATCH-P,phs002152
MATCH-R,phs002029
MATCH-S2,phs002178
MATCH-W,phs001948
MATCH-Z1A,phs001973
MATCH-Z1D,phs001859
MMRF-COMMPASS,phs000748
MP2PRT-WT,phs001965
NCICCR-DLBCL
OHSU-CNL,phs001799
REBC-THYR
TARGET-ALL-P1,phs000463
TARGET-ALL-P3
TARGET-CCSK,phs000466
TARGET-OS,phs000468
TARGET-WT,phs000471
TCGA-BLCA
TCGA-CESC
TCGA-COAD
TCGA-ESCA
TCGA-HNSC
TCGA-KIRC
TCGA-LAML
TCGA-LIHC
TCGA-LUSC
TCGA-OV
TCGA-PCPG
TCGA-READ
TCGA-SKCM
TCGA-TGCT
TCGA-THYM
TCGA-UCS
TNP SARDANA
Vanderbilt
WUSTL
Research Study Identifiers
Abdominal fibromatosis
unknown
Display
3000
2500
2000
1500
1000
500
0

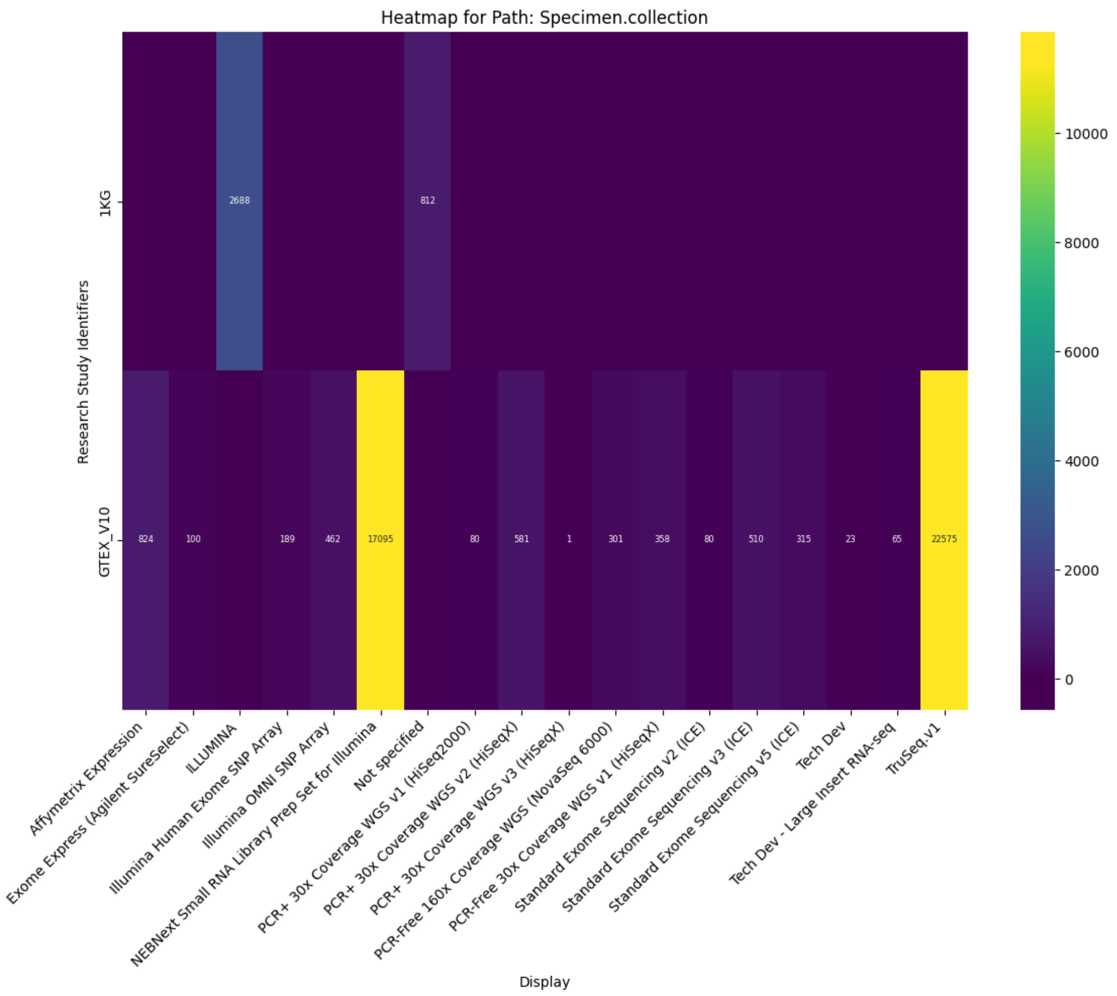

Heatmap for Path: Specimen.collection
Research Study Identifiers
1KG
GTEX_V10
2688
812
824
100
189
462
17095
80
581
1
301
358
80
510
315
23
65
22575
Affymetrix Expression
Exome Express (Agilent SureSelect)
ILLUMINA
Illumina Human Exome SNP Array
Illumina OMNI SNP Array
NEBNext Small RNA Library Prep Set for Illumina
Not specified
PCR+ 30x Coverage WGS v1 (HiSeq2000)
PCR+ 30x Coverage WGS v2 (HiSeqX)
PCR+ 30x Coverage WGS v3 (HiSeqX)
PCR-Free 160x Coverage WGS (NovaSeq 6000)
PCR-Free 30x Coverage WGS v1 (HiSeqX)
Standard Exome Sequencing v2 (ICE)
Standard Exome Sequencing v3 (ICE)
Standard Exome Sequencing v5 (ICE)
Tech Dev
Tech Dev - Large Insert RNA-seq
TruSeq.v1
Display
10000
8000
6000
4000
2000
0

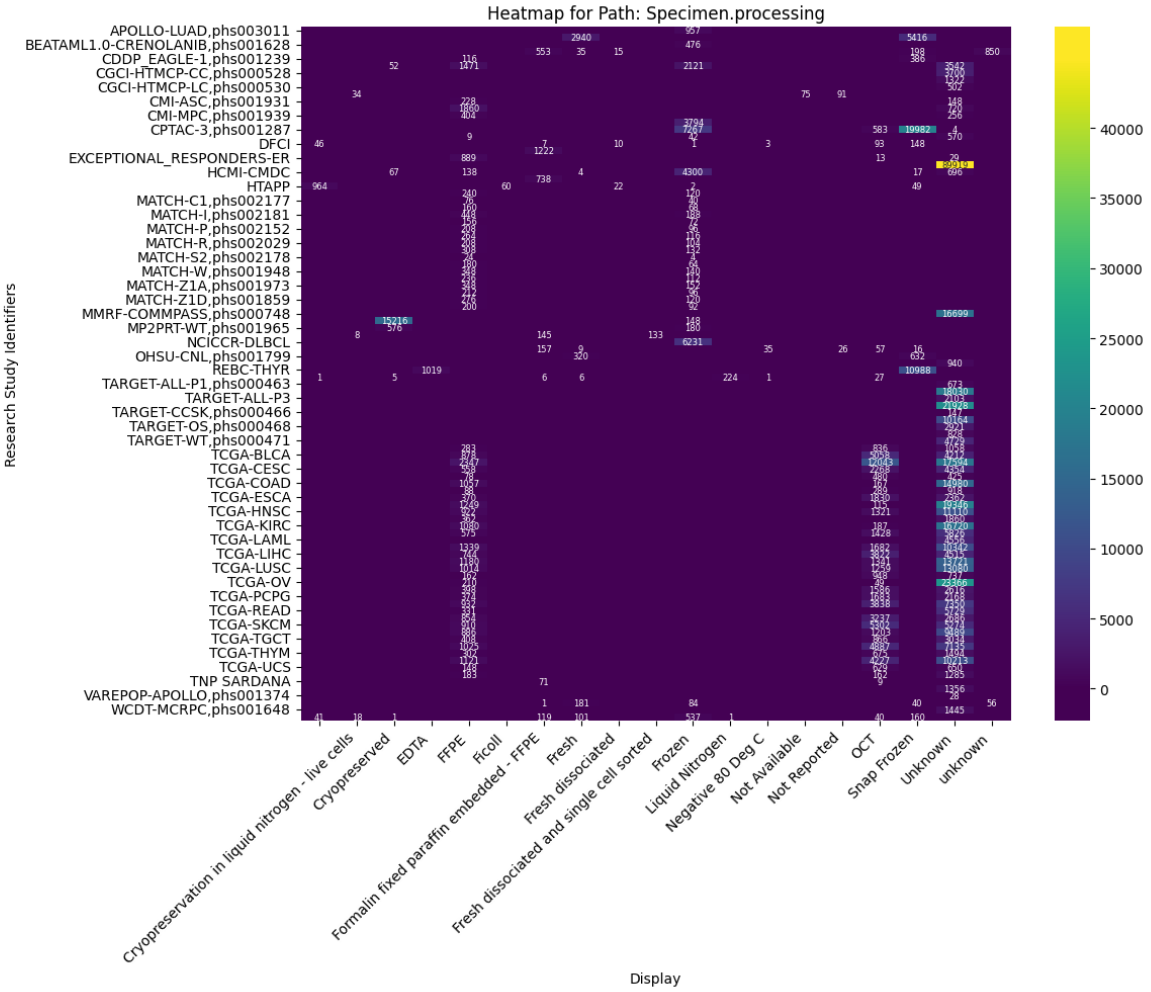
Heatmap for Path: Specimen.processing
Research Study Identifiers
APOLLO-LUAD,phs003011
BEATAML1.0-CRENOLANIB,phs001628
CDDP_EAGLE-1,phs001239
CGCI-HTMCP-CC,phs000528
CGCI-HTMCP-LC,phs000530
CMI-ASC,phs001931
CMI-MPC,phs001939
CPTAC-3,phs001287
DFCI
EXCEPTIONAL_RESPONDERS-ER
HCMI-CMDC
HTAPP
MATCH-C1,phs002177
MATCH-I,phs002181
MATCH-P,phs002152
MATCH-R,phs002029
MATCH-S2,phs002178
MATCH-W,phs001948
MATCH-Z1A,phs001973
MATCH-Z1D,phs001859
MMRF-COMMPASS,phs000748
MP2PRT-WT,phs001965
NCICCR-DLBCL
OHSU-CNL,phs001799
REBC-THYR
TARGET-ALL-P1,phs000463
TARGET-ALL-P3
TARGET-CCSK,phs000466
TARGET-OS,phs000468
TARGET-WT,phs000471
TCGA-BLCA
TCGA-CESC
TCGA-COAD
TCGA-ESCA
TCGA-HNSC
TCGA-KIRC
TCGA-LAML
TCGA-LIHC
TCGA-LUSC
TCGA-OV
TCGA-PCPG
TCGA-READ
TCGA-SKCM
TCGA-TGCT
TCGA-THYM
TCGA-UCS
TNP SARDANA
VAREPOP-APOLLO,phs001374
WCDT-MCRPC,phs001648
Cryopreservation in liquid nitrogen - live cells
Cryopreserved
EDTA
FFPE
Ficoll
Formalin fixed paraffin embedded - FFPE
Fresh
Fresh dissociated
Fresh dissociated and single cell sorted
Frozen
Liquid Nitrogen
Negative 80 Deg C
Not Available
Not Reported
OCT
Snap Frozen
Unknown
unknown
Display
40000
35000
30000
25000
20000
15000
10000
5000
0

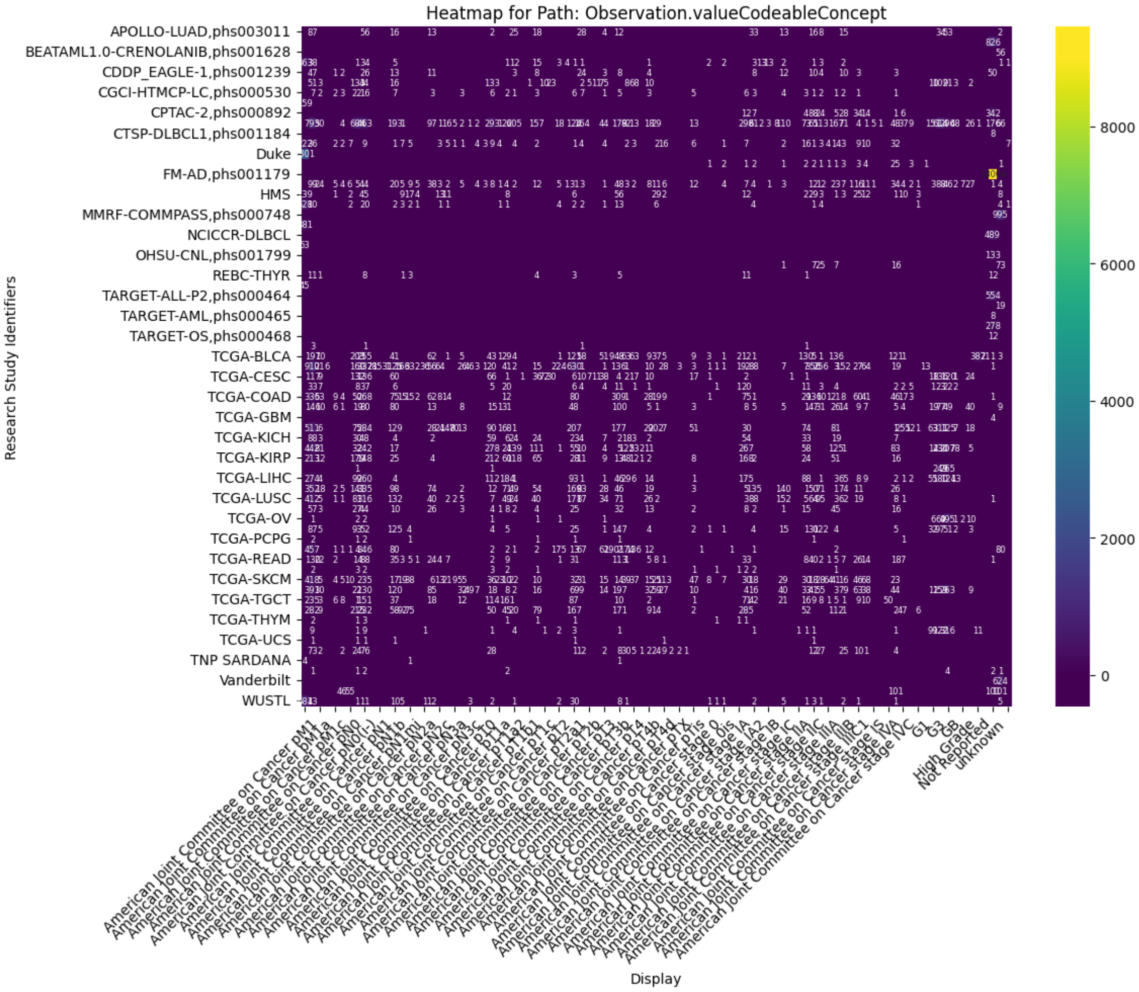
Heatmap for Path: Observation.valueCodeableConcept
Research Study Identifiers
APOLLO-LUAD,phs003011
BEATAML1.0-CRENOLANIB,phs001628
CDDP_EAGLE-1,phs001239
CGCI-HTMCP-LC,phs000530
CPTAC-2,phs000892
CTSP-DLBCL1,phs001184
Duke
FM-AD,phs001179
HMS
MMRF-COMMPASS,phs000748
NCICCR-DLBCL
OHSU-CNL,phs001799
REBC-THYR
TARGET-ALL-P2,phs000464
TARGET-AML,phs000465
TARGET-OS,phs000468
TCGA-BLCA
TCGA-CESC
TCGA-COAD
TCGA-GBM
TCGA-KICH
TCGA-KIRP
TCGA-LIHC
TCGA-LUSC
TCGA-OV
TCGA-PCPG
TCGA-READ
TCGA-SKCM
TCGA-TGCT
TCGA-THYM
TCGA-UCS
TNP SARDANA
Vanderbilt
WUSTL
Display
High Grade
Not Reported
unknown
8000
6000
4000
2000
0

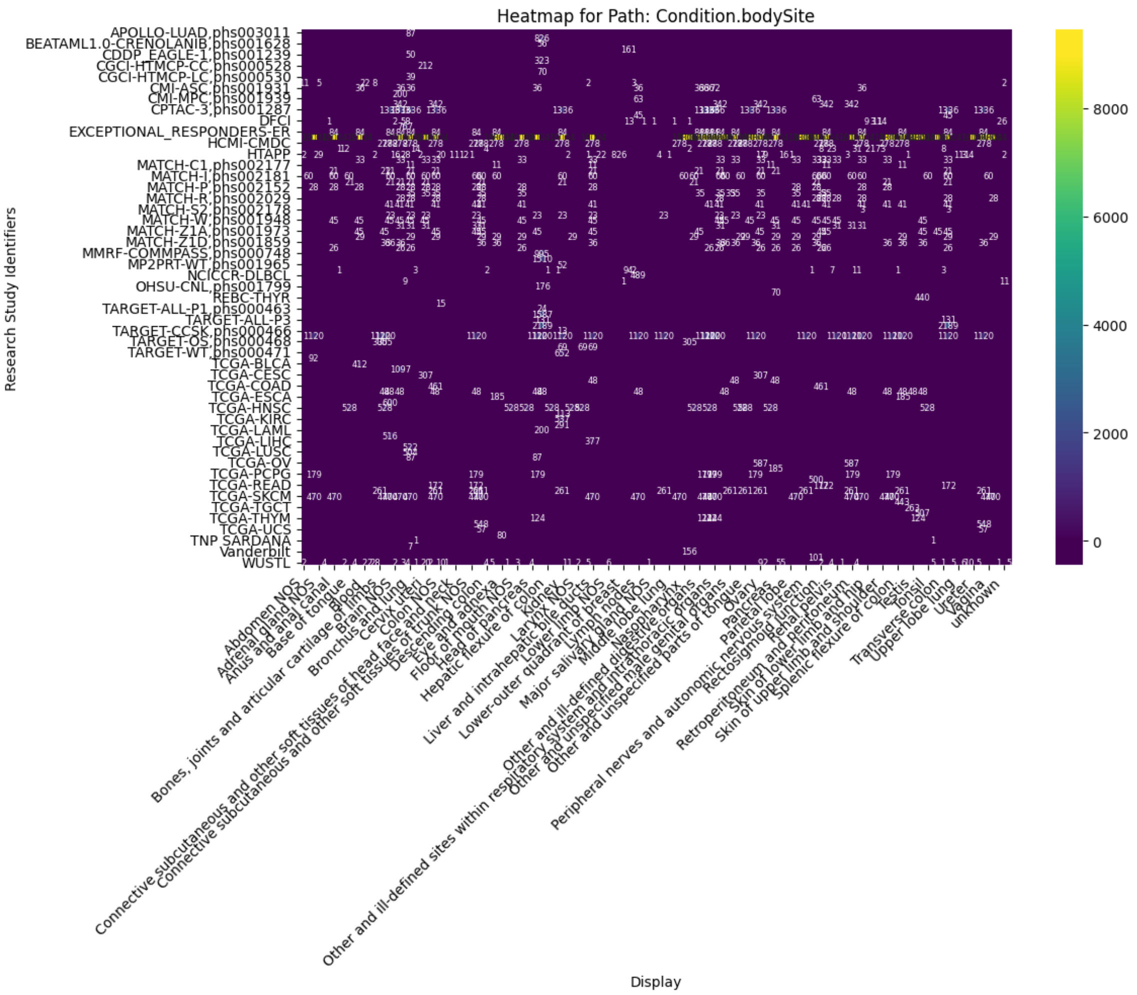
Heatmap for Path: Condition.bodySite
Research Study Identifiers
Display
APOLLO-LUAD,phs003011
BEATAML1.0-CRENOLANIB,phs001628
CDDP_EAGLE-1,phs001239
CGCI-HTMCP-CC,phs000528
CGCI-HTMCP-LC,phs000530
CMI-ASC,phs001931
CMI-MPC,phs001939
CPTAC-3,phs001287
DFCI
EXCEPTIONAL_RESPONDERS-ER
HCMI-CMDC
HTAPP
MATCH-C1,phs002177
MATCH-I,phs002181
MATCH-P,phs002152
MATCH-R,phs002029
MATCH-S2,phs002178
MATCH-W,phs001948
MATCH-Z1A,phs001973
MATCH-Z1D,phs001859
MMRF-COMMPASS,phs000748
MP2PRT-WT,phs001965
NCICCR-DLBCL
OHSU-CNL,phs001799
REBC-THYR
TARGET-ALL-P1,phs000463
TARGET-ALL-P3
TARGET-CCSK,phs000466
TARGET-OS,phs000468
TARGET-WT,phs000471
TCGA-BLCA
TCGA-CESC
TCGA-COAD
TCGA-ESCA
TCGA-HNSC
TCGA-KIRC
TCGA-LAML
TCGA-LIHC
TCGA-LUSC
TCGA-OV
TCGA-PCPG
TCGA-READ
TCGA-SKCM
TCGA-TGCT
TCGA-THYM
TCGA-UCS
TNP SARDANA
Vanderbilt
WUSTL
Abdomen NOS
Adrenal gland NOS
Anus and anal canal
Base of tongue
Blood
Bones, joints and articular cartilage of limbs
Brain NOS
Bronchus and lung
Cervix uteri
Colon NOS
Connective subcutaneous and other soft tissues of head face and neck
Connective subcutaneous and other soft tissues of trunk NOS
Descending colon
Eye and adnexa
Floor of mouth NOS
Head of pancreas
Hepatic flexure of colon
Kidney
Larynx NOS
Liver and intrahepatic bile ducts
Lower limb NOS
Lower-outer quadrant of breast
Lymph nodes
Major salivary gland NOS
Middle lobe lung
Nasopharynx
Other and ill-defined digestive organs
Other and ill-defined sites within respiratory system and intrathoracic organs
Other and unspecified male genital organs
Other and unspecified parts of tongue
Ovary
Pancreas
Parietal lobe
Peripheral nerves and autonomic nervous system
Rectosigmoid junction
Renal pelvis
Retroperitoneum and peritoneum
Skin of lower limb and hip
Skin of upper limb and shoulder
Splenic flexure of colon
Testis
Tonsil
Transverse colon
Upper lobe lung
Ureter
Vagina
unknown
8000
6000
4000
2000
0

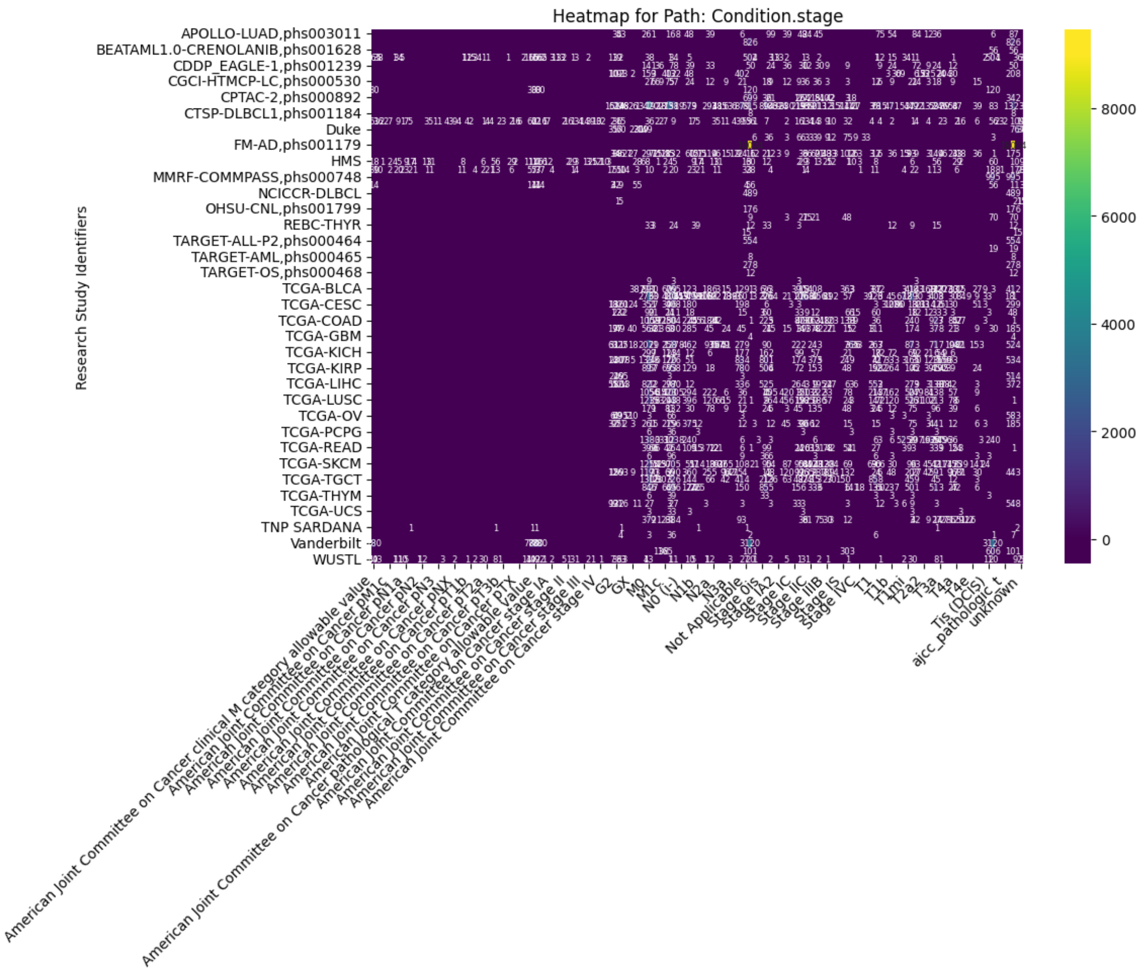

Heatmap for Path: Condition.stage
Research Study Identifiers
APOLLO-LUAD,phs003011
BEATAML1.0-CRENOLANIB,phs001628
CDDP_EAGLE-1,phs001239
CGCI-HTMCP-LC,phs000530
CPTAC-2,phs000892
CTSP-DLBCL1,phs001184
Duke
FM-AD,phs001179
HMS
MMRF-COMMPASS,phs000748
NCICCR-DLBCL
OHSU-CNL,phs001799
REBC-THYR
TARGET-ALL-P2,phs000464
TARGET-AML,phs000465
TARGET-OS,phs000468
TCGA-BLCA
TCGA-CESC
TCGA-COAD
TCGA-GBM
TCGA-KICH
TCGA-KIRP
TCGA-LIHC
TCGA-LUSC
TCGA-OV
TCGA-PCPG
TCGA-READ
TCGA-SKCM
TCGA-TGCT
TCGA-THYM
TCGA-UCS
TNP SARDANA
Vanderbilt
WUSTL
American Joint Committee on Cancer clinical M category allowable value
American Joint Committee on Cancer pathological T category allowable value
G2
GX
M0
M1c
N0 (i-)
N1b
N2a
N3a
Not Applicable
Stage 0is
Stage IA2
Stage IC
Stage IIC
Stage IIIB
Stage IS
Stage IVC
T1
T1b
T1mi
T2a2
T3a
T4a
T4e
Tis (DCIS)
ajcc_pathologic_t
unknown
Display
8000
6000
4000
2000
0

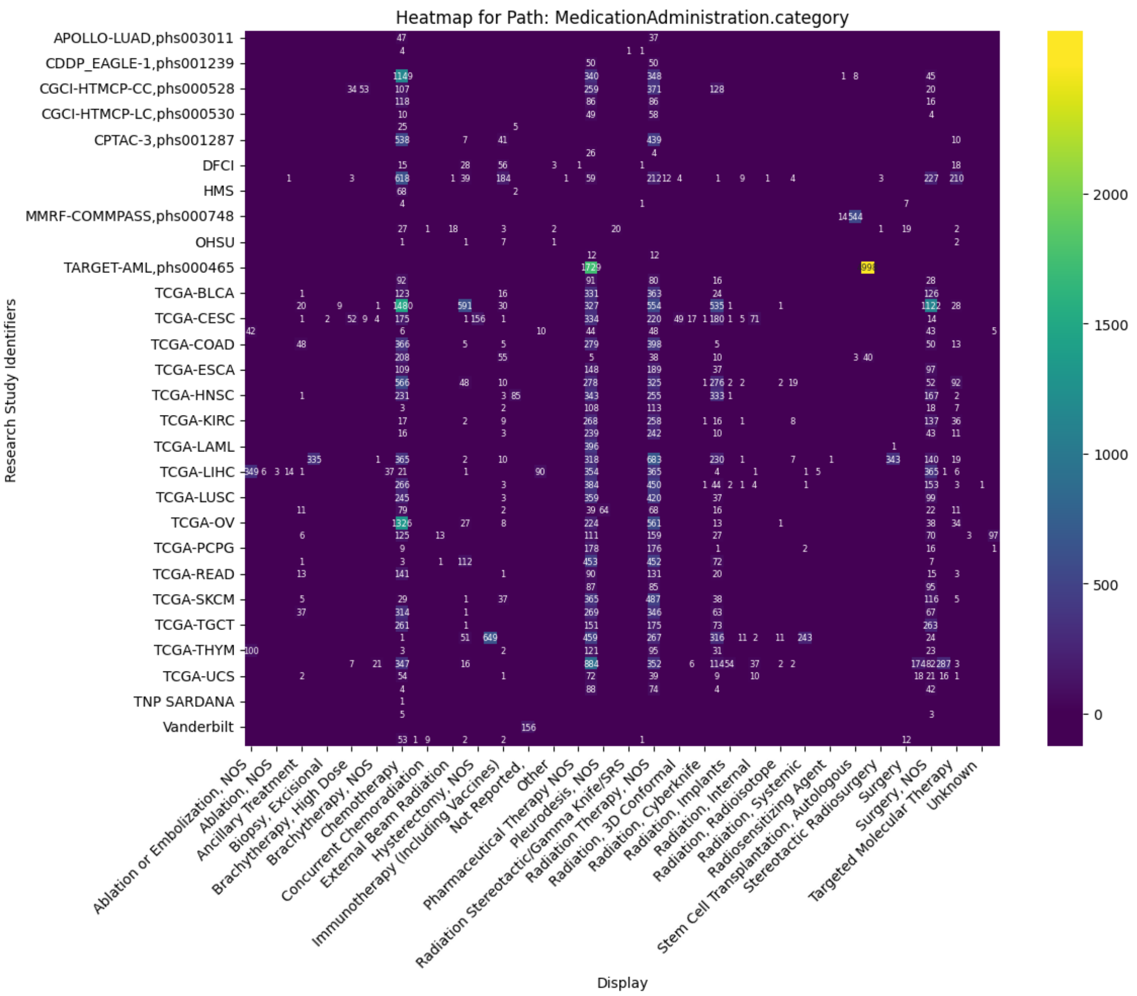

Heatmap for Path: MedicationAdministration.category
Research Study Identifiers
APOLLO-LUAD,phs003011
CDDP_EAGLE-1,phs001239
CGCI-HTMCP-CC,phs000528
CGCI-HTMCP-LC,phs000530
CPTAC-3,phs001287
DFCI
HMS
MMRF-COMMPASS,phs000748
OHSU
TARGET-AML,phs000465
TCGA-BLCA
TCGA-CESC
TCGA-COAD
TCGA-ESCA
TCGA-HNSC
TCGA-KIRC
TCGA-LAML
TCGA-LIHC
TCGA-LUSC
TCGA-OV
TCGA-PCPG
TCGA-READ
TCGA-SKCM
TCGA-TGCT
TCGA-THYM
TCGA-UCS
TNP SARDANA
Vanderbilt
Display
Ablation or Embolization, NOS
Ablation, NOS
Ancillary Treatment
Biopsy, Excisional
Brachytherapy, High Dose
Brachytherapy, NOS
Chemotherapy
Concurrent Chemoradiation
External Beam Radiation
Hysterectomy, NOS
Immunotherapy (Including Vaccines)
Not Reported,
Other
Pharmaceutical Therapy NOS
Pleurodesis, NOS
Radiation Stereotactic/Gamma Knife/SRS
Radiation Therapy, NOS
Radiation, 3D Conformal
Radiation, Cyberknife
Radiation, Implants
Radiation, Internal
Radiation, Radioisotope
Radiation, Systemic
Radiosensitizing Agent
Stem Cell Transplantation, Autologous
Stereotactic Radiosurgery
Surgery
Surgery, NOS
Targeted Molecular Therapy
Unknown
2000
1500
1000
500
0

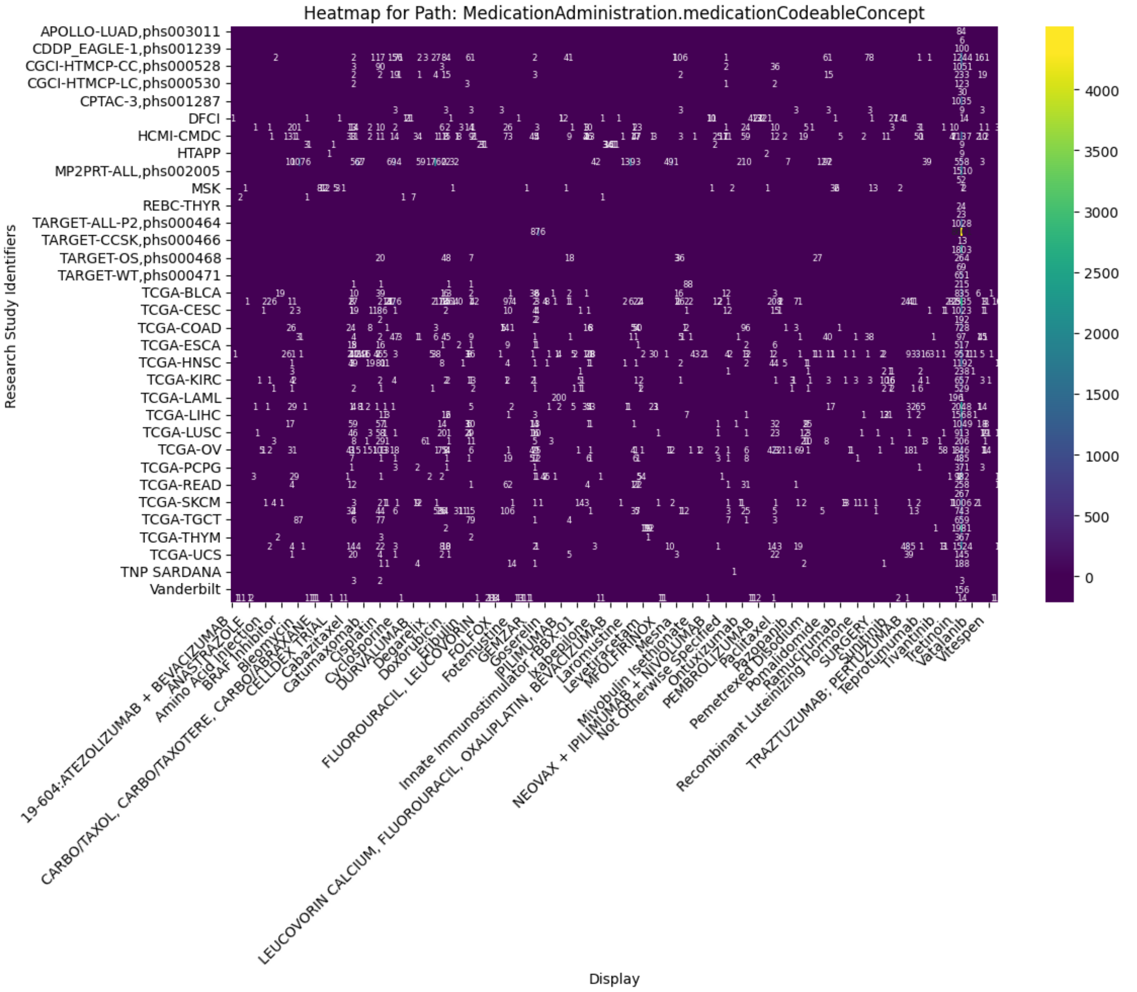
Heatmap for Path: MedicationAdministration.medicationCodeableConcept
Research Study Identifiers
APOLLO-LUAD,phs003011
CDDP_EAGLE-1,phs001239
CGCI-HTMCP-CC,phs000528
CGCI-HTMCP-LC,phs000530
CPTAC-3,phs001287
DFCI
HCMI-CMDC
HTAPP
MP2PRT-ALL,phs002005
MSK
REBC-THYR
TARGET-ALL-P2,phs000464
TARGET-CCSK,phs000466
TARGET-OS,phs000468
TARGET-WT,phs000471
TCGA-BLCA
TCGA-CESC
TCGA-COAD
TCGA-ESCA
TCGA-HNSC
TCGA-KIRC
TCGA-LAML
TCGA-LIHC
TCGA-LUSC
TCGA-OV
TCGA-PCPG
TCGA-READ
TCGA-SKCM
TCGA-TGCT
TCGA-THYM
TCGA-UCS
TNP SARDANA
Vanderbilt
19-604:ATEZOLIZUMAB + BEVACIZUMAB
ANASTRAZOLE
Amino Acid Injection
BRAF Inhibitor
Bleomycin
CARBO/TAXOL, CARBO/TAXOTERE, CARBO/ABRAXANE
CELLDEX TRIAL
Cabazitaxel
Catumaxomab
Cisplatin
Cyclosporine
DURVALUMAB
Degarelix
Doxorubicin
Eribulin
FLUOROURACIL, LEUCOVORIN
FOLFOX
Fotemustine
GEMZAR
Goserelin
IPILIMUMAB
Innate Immunostimulator rBBX-01
Ixabepilone
LEUCOVORIN CALCIUM, FLUOROURACIL, OXALIPLATIN, BEVACIZUMAB
Laromustine
Levetiracetam
MFOLFIRINOX
Mesna
Mivobulin Isethionate
NEOVAX + IPILIMUMAB + NIVOLUMAB
Not Otherwise Specified
Ontuxizumab
PEMBROLIZUMAB
Paclitaxel
Pazopanib
Pemetrexed Disodium
Pomalidomide
Ramucirumab
Recombinant Luteinizing Hormone
SURGERY
Sunitinib
TRAZTUZUMAB; PERTUZUMAB
Teprotumumab
Tivantinib
Tretinoin
Vatalanib
Vitespen
Display
4000
3500
3000
2500
2000
1500
1000
500
0

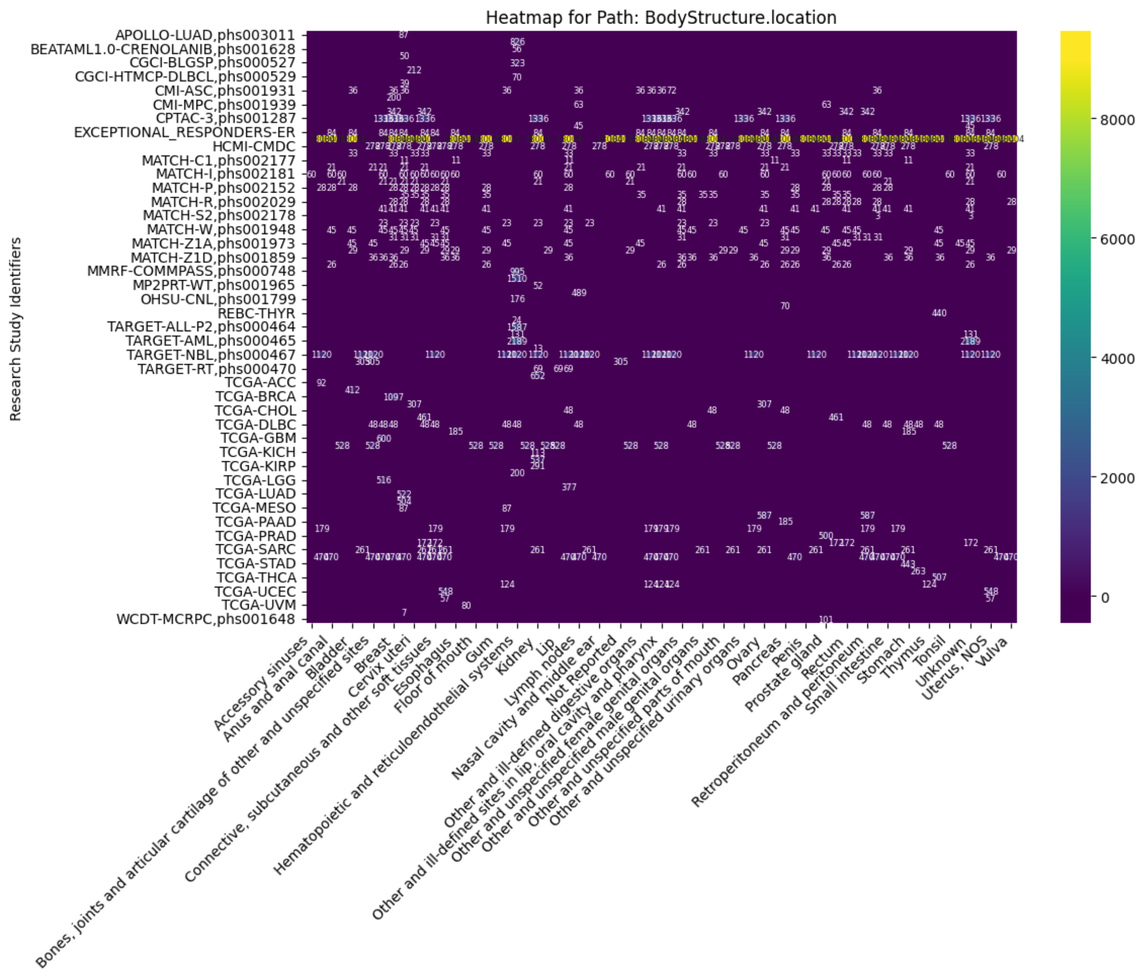
Heatmap for Path: BodyStructure.location
Research Study Identifiers
Display

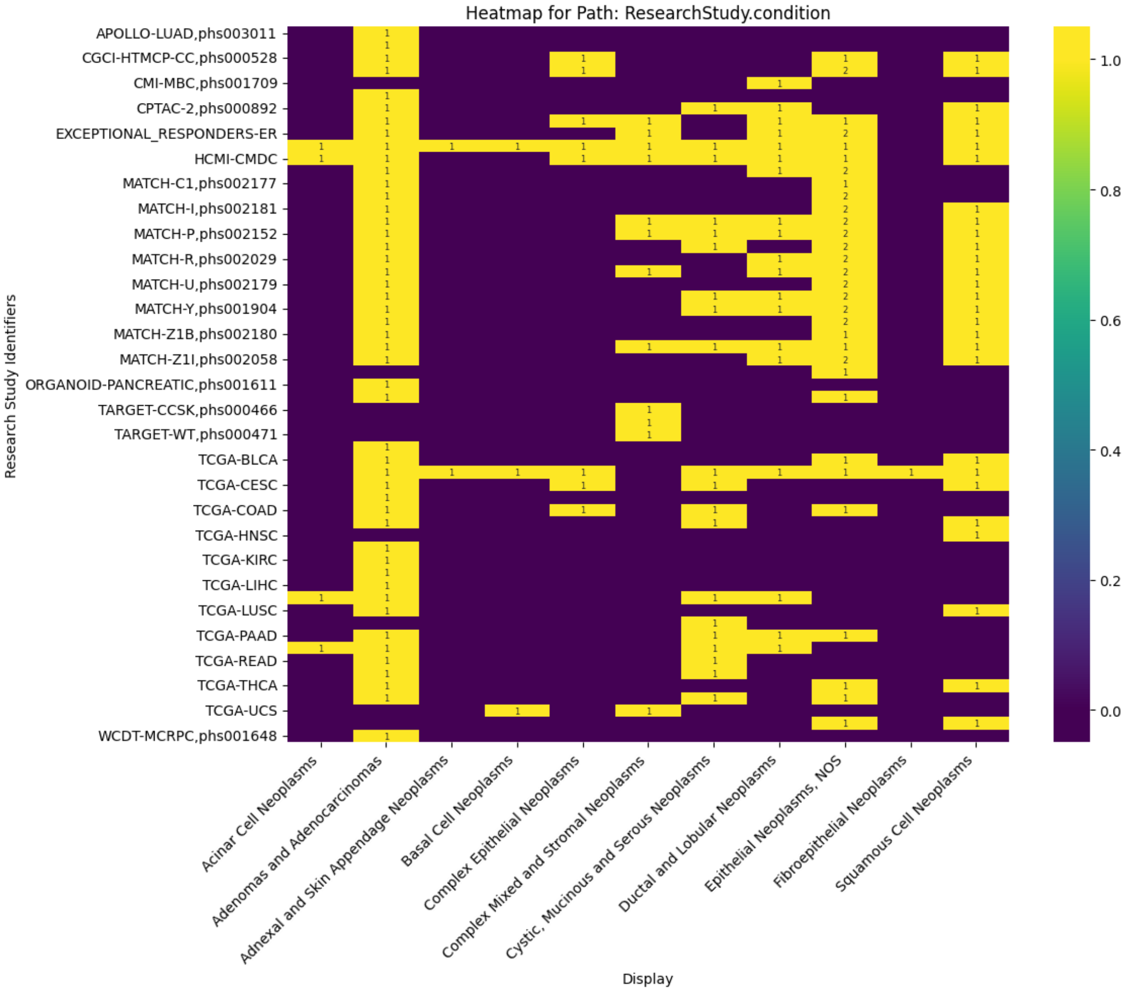

Heatmap for Path: ResearchStudy.condition
Research Study Identifiers
APOLLO-LUAD,phs003011
CGCI-HTMCP-CC,phs000528
CMI-MBC,phs001709
CPTAC-2,phs000892
EXCEPTIONAL_RESPONDERS-ER
HCMI-CMDC
MATCH-C1,phs002177
MATCH-I,phs002181
MATCH-P,phs002152
MATCH-R,phs002029
MATCH-U,phs002179
MATCH-Y,phs001904
MATCH-Z1B,phs002180
MATCH-Z1I,phs002058
ORGANOID-PANCREATIC,phs001611
TARGET-CCSK,phs000466
TARGET-WT,phs000471
TCGA-BLCA
TCGA-CESC
TCGA-COAD
TCGA-HNSC
TCGA-KIRC
TCGA-LIHC
TCGA-LUSC
TCGA-PAAD
TCGA-READ
TCGA-THCA
TCGA-UCS
WCDT-MCRPC,phs001648
Acinar Cell Neoplasms
Adenomas and Adenocarcinomas
Adnexal and Skin Appendage Neoplasms
Basal Cell Neoplasms
Complex Epithelial Neoplasms
Complex Mixed and Stromal Neoplasms
Cystic, Mucinous and Serous Neoplasms
Ductal and Lobular Neoplasms
Epithelial Neoplasms, NOS
Fibroepithelial Neoplasms
Squamous Cell Neoplasms
Display
1.0
0.8
0.6
0.4
0.2
0.0

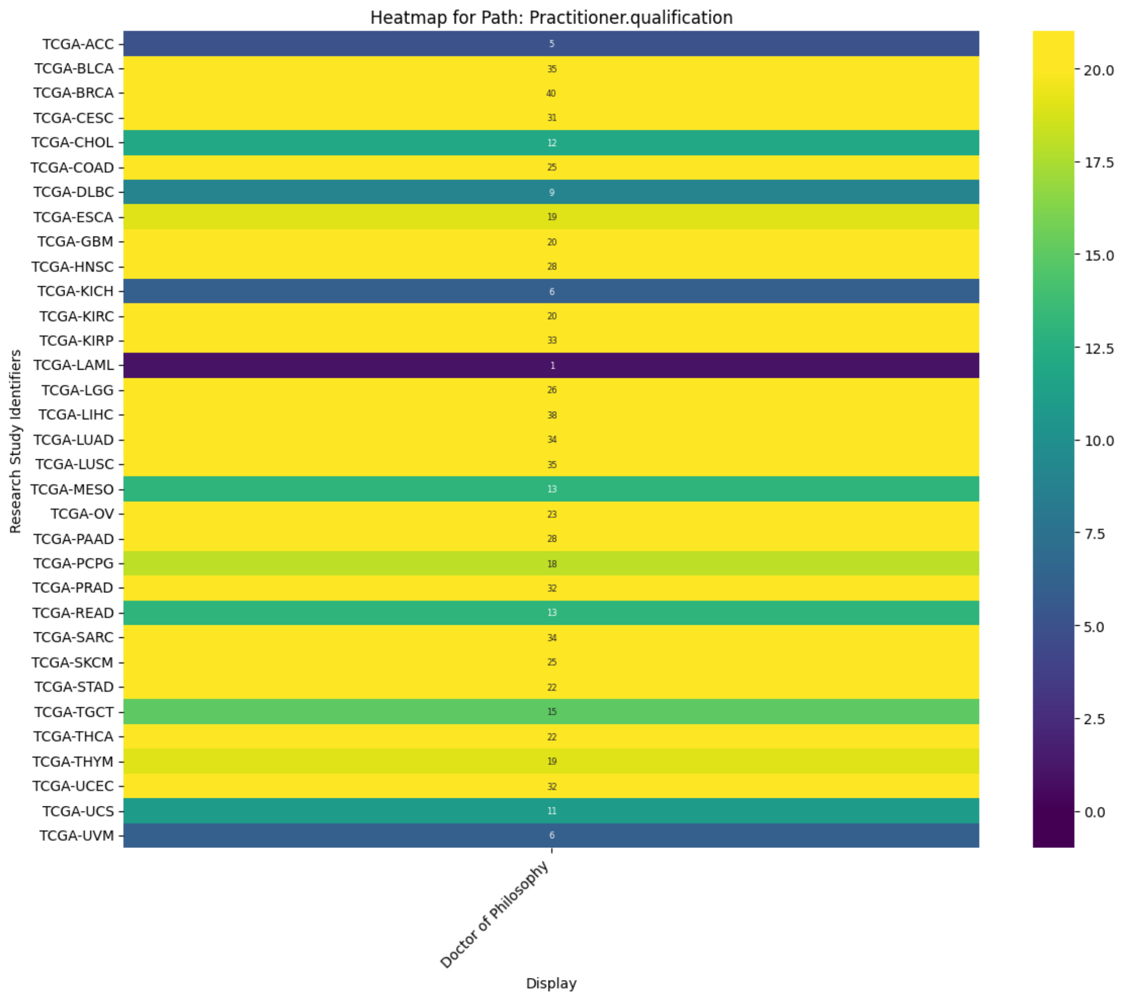
Heatmap for Path: Practitioner.qualification
Research Study Identifiers
TCGA-ACC
TCGA-BLCA
TCGA-BRCA
TCGA-CESC
TCGA-CHOL
TCGA-COAD
TCGA-DLBC
TCGA-ESCA
TCGA-GBM
TCGA-HNSC
TCGA-KICH
TCGA-KIRC
TCGA-KIRP
TCGA-LAML
TCGA-LGG
TCGA-LIHC
TCGA-LUAD
TCGA-LUSC
TCGA-MESO
TCGA-OV
TCGA-PAAD
TCGA-PCPG
TCGA-PRAD
TCGA-READ
TCGA-SARC
TCGA-SKCM
TCGA-STAD
TCGA-TGCT
TCGA-THCA
TCGA-THYM
TCGA-UCEC
TCGA-UCS
TCGA-UVM
5
35
40
31
12
25
9
19
20
28
6
20
33
1
26
38
34
35
13
23
28
18
32
13
34
25
22
15
22
19
32
11
6
Doctor of Philosophy
Display
20.0
17.5
15.0
12.5
10.0
7.5
5.0
2.5
0.0

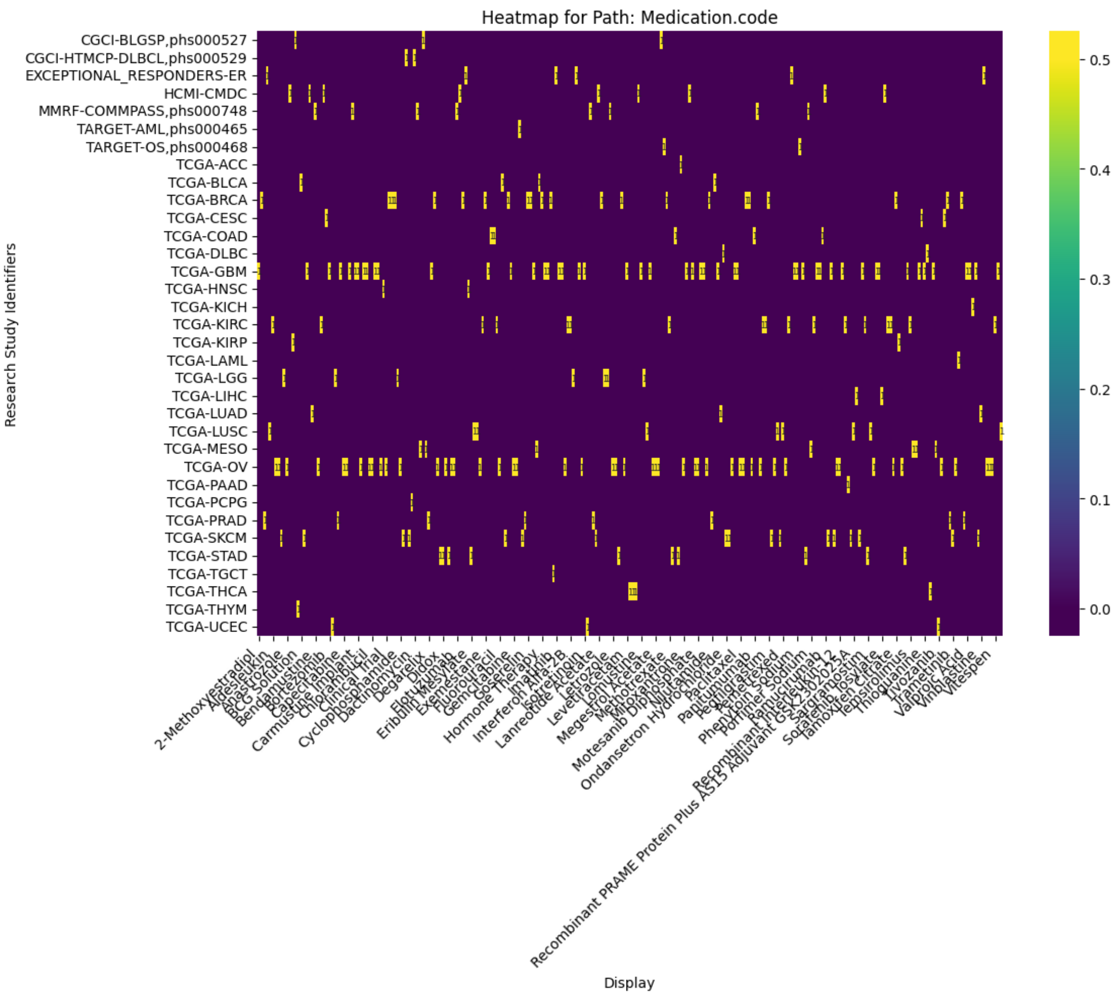


### FHIR Extensions¶

A FHIR extension provides a way to add custom data elements to existing FHIR resources, enabling the representation of information that is not defined in the core FHIR specification. They allow for flexibility and extensibility within FHIR while maintaining interoperability.

The vocuabulary Observation includes a summary of these extensions

```
# extensions with ranges
combined_extension_range_counts = df[['research_study_identifiers', 'path', 'extension_url', 'low', 'high', 'url']]
combined_extension_range_counts = combined_extension_range_counts[combined_extension_range_counts['extension_url'] != '']
combined_extension_range_counts = combined_extension_range_counts[combined_extension_range_counts['low'] != '']
combined_extension_range_counts
```

| | research_study_identifiers | path | extension_url | low | high | url |
|---|---|---|---|---|---|---|
| **12** | ICGC-LUCA_KR | DocumentReference.extension | https://nih-ncpi.github.io/ncpi-fhir-ig-2/Stru... | 365.0 | 206822704094.0 | |
| **3739** | Cellosaurus | Patient.extension | http://hl7.org/fhir/SearchParameter/patient-ex... | 1.0 | 94.0 | |
| **3771** | 1KG | DocumentReference.extension | https://nih-ncpi.github.io/ncpi-fhir-ig-2/Stru... | 36047.0 | 16099441977.0 | |
| **3881** | BEATAML1.0-CRENOLANIB,phs001628 | DocumentReference.extension | https://nih-ncpi.github.io/ncpi-fhir-ig-2/Stru... | 4220.0 | 45589228155.0 | |
| **3886** | BEATAML1.0-CRENOLANIB,phs001628 | Patient.extension | http://hl7.org/fhir/SearchParameter/patient-ex... | 24.0 | 87.0 | |
| **...** | ... | ... | ... | ... | ... | ... |
| **20886** | TCGA-HNSC | DocumentReference.extension | https://nih-ncpi.github.io/ncpi-fhir-ig-2/Stru... | 229.0 | 578035949313.0 | |
| **20899** | TCGA-HNSC | Patient.extension | http://hl7.org/fhir/SearchParameter/patient-ex... | 19.0 | 89.0 | |
| **21249** | TCGA-KIRC | DocumentReference.extension | https://nih-ncpi.github.io/ncpi-fhir-ig-2/Stru... | 229.0 | 481092668932.0 | |
| **21259** | TCGA-KIRC | Patient.extension | http://hl7.org/fhir/SearchParameter/patient-ex... | 26.0 | 89.0 | |
| **21834** | HCMI-CMDC | DocumentReference.extension | https://nih-ncpi.github.io/ncpi-fhir-ig-2/Stru... | 229.0 | 484490562985.0 | |

141 rows × 6 columns

```
# extensions with codes
combined_extension_code_counts = df[['research_study_identifiers', 'path', 'extension_url', 'display', 'url', 'count']]
combined_extension_code_counts = combined_extension_code_counts[combined_extension_code_counts['extension_url'] != '']
combined_extension_code_counts = combined_extension_code_counts[combined_extension_code_counts['display'] != '']
combined_extension_code_counts
```

| | research_study_identifiers | path | extension_url | display | url | count |
|---|---|---|---|---|---|---|
| **13** | ICGC-LUCA_KR | Patient.extension | http://hl7.org/fhir/us/core/StructureDefinitio... | M | https://google-fhir.fhir-aggregator.org/Patien... | 344.0 |
| **14** | ICGC-LUCA_KR | Patient.extension | http://hl7.org/fhir/us/core/StructureDefinitio... | F | https://google-fhir.fhir-aggregator.org/Patien... | 56.0 |
| **41** | GTEX_V10 | DocumentReference.extension | https://nih-ncpi.github.io/ncpi-fhir-ig-2/Stru... | 16K | https://google-fhir.fhir-aggregator.org/Docume... | 1.0 |
| **42** | GTEX_V10 | DocumentReference.extension | https://nih-ncpi.github.io/ncpi-fhir-ig-2/Stru... | 11K | https://google-fhir.fhir-aggregator.org/Docume... | 1.0 |
| **43** | GTEX_V10 | DocumentReference.extension | https://nih-ncpi.github.io/ncpi-fhir-ig-2/Stru... | 11M | https://google-fhir.fhir-aggregator.org/Docume... | 1.0 |
| **...** | ... | ... | ... | ... | ... | ... |
| **21841** | HCMI-CMDC | Patient.extension | http://hl7.org/fhir/us/core/StructureDefinitio... | black or african american | https://google-fhir.fhir-aggregator.org/Patien... | 16.0 |
| **21842** | HCMI-CMDC | Patient.extension | http://hl7.org/fhir/us/core/StructureDefinitio... | not hispanic or latino | https://google-fhir.fhir-aggregator.org/Patien... | 158.0 |
| **21843** | HCMI-CMDC | Patient.extension | http://hl7.org/fhir/us/core/StructureDefinitio... | not reported | https://google-fhir.fhir-aggregator.org/Patien... | 42.0 |
| **21844** | HCMI-CMDC | Patient.extension | http://hl7.org/fhir/us/core/StructureDefinitio... | Unknown | https://google-fhir.fhir-aggregator.org/Patien... | 67.0 |
| **21845** | HCMI-CMDC | Patient.extension | http://hl7.org/fhir/us/core/StructureDefinitio... | hispanic or latino | https://google-fhir.fhir-aggregator.org/Patien... | 11.0 |

934 rows × 6 columns

```
import seaborn as sns
import matplotlib.pyplot as plt
import pandas as pd

# Create heatmap for each extension

# Get unique extensions
extension_urls = combined_extension_code_counts['extension_url'].unique()
combined_extension_code_counts = combined_extension_code_counts.infer_objects(copy=False)

for extension_url in extension_urls:
    # Filter data for the current extension
    filtered_data = combined_extension_code_counts[combined_extension_code_counts['extension_url'] == extension_url]

    # Pivot the filtered data for the heatmap
    heatmap_data = filtered_data.pivot_table(index='research_study_identifiers',
                                             columns=['display'],
                                             values='count',
                                             fill_value=0.0)

    # Create the heatmap
    plt.figure(figsize=(12, 10))  # Adjust figsize as needed

    # sns.heatmap(heatmap_data, annot=True, annot_kws={"size": 6}, fmt=".0f", cmap='viridis', cbar=False)

    # Create a mask to identify zero values
    mask = heatmap_data == 0

 # Custom function to format annotations
    def heatmap_annot(val, **kwargs):
        if val == 0:  # Hide annotations for zero values
            return ""
        else:
            return f"{val:.0f}"  # Format non-zero values as integers


    # Generate heatmap with the custom annotation function
    ax = sns.heatmap(heatmap_data, annot=True, fmt=".0f", cmap='viridis',
                cbar_kws={'extend': 'both', 'extendrect': True},
                vmin=0,  # Set minimum value for color scale
                vmax=heatmap_data.max().max() / 2,  # Adjust maximum value
                annot_kws={"size": 6})  # Reduce font size


    # Apply the custom annotation function to each cell
    for text in ax.texts:
        text.set_text(heatmap_annot(float(text.get_text())))

    plt.title(f'Heatmap for Path: {extension_url}')  # Set title with path name
    plt.xlabel('Display')
    plt.ylabel('Research Study Identifiers')
    plt.xticks(rotation=45, ha='right')
    plt.tight_layout()
    plt.show()
```

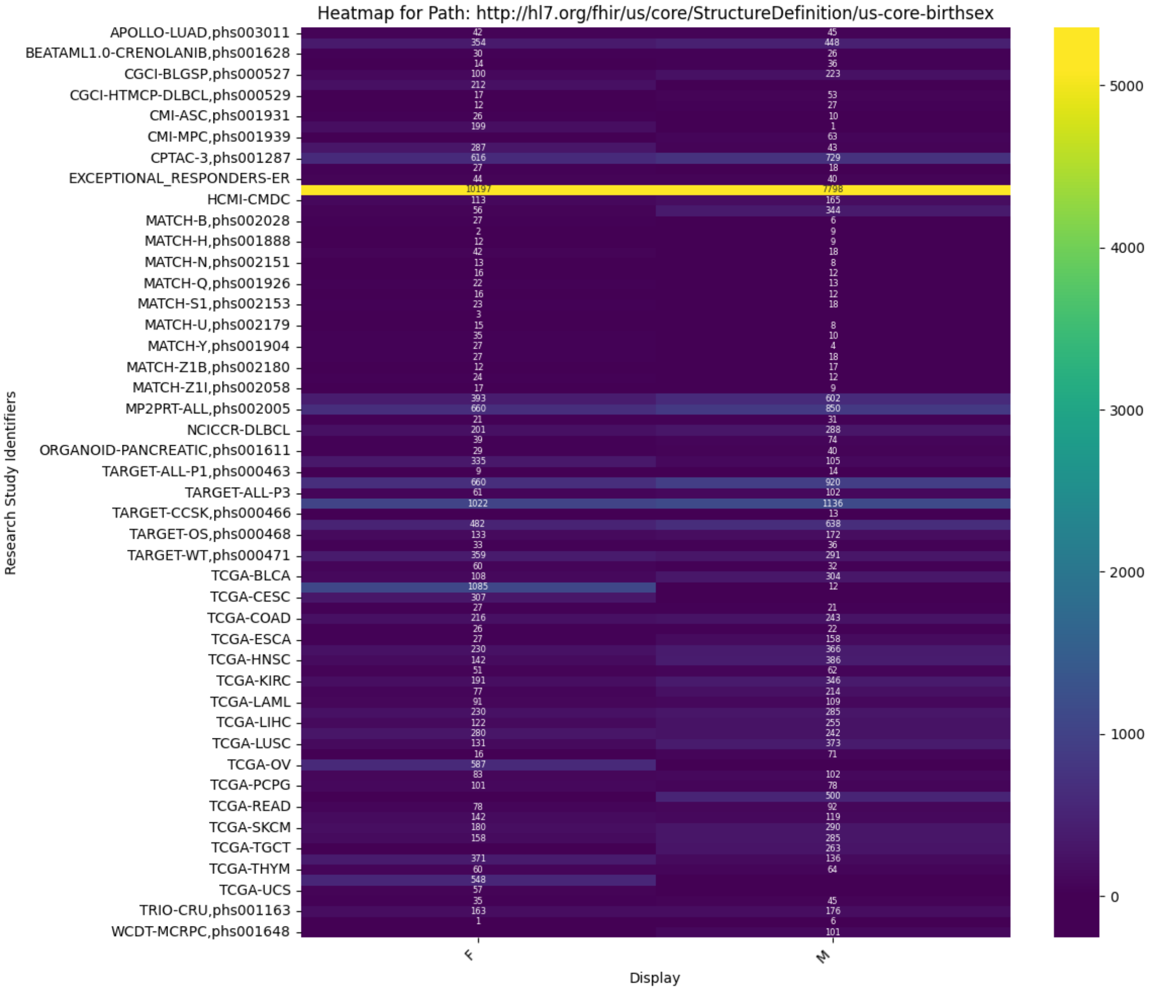

Heatmap for Path: http://hl7.org/fhir/us/core/StructureDefinition/us-core-birthsex
Research Study Identifiers
APOLLO-LUAD,phs003011
BEATAML1.0-CRENOLANIB,phs001628
CGCI-BLGSP,phs000527
CGCI-HTMCP-DLBCL,phs000529
CMI-ASC,phs001931
CMI-MPC,phs001939
CPTAC-3,phs001287
EXCEPTIONAL_RESPONDERS-ER
HCMI-CMDC
MATCH-B,phs002028
MATCH-H,phs001888
MATCH-N,phs002151
MATCH-Q,phs001926
MATCH-S1,phs002153
MATCH-U,phs002179
MATCH-Y,phs001904
MATCH-Z1B,phs002180
MATCH-Z1I,phs002058
MP2PRT-ALL,phs002005
NCICCR-DLBCL
ORGANOID-PANCREATIC,phs001611
TARGET-ALL-P1,phs000463
TARGET-ALL-P3
TARGET-CCSK,phs000466
TARGET-OS,phs000468
TARGET-WT,phs000471
TCGA-BLCA
TCGA-CESC
TCGA-COAD
TCGA-ESCA
TCGA-HNSC
TCGA-KIRC
TCGA-LAML
TCGA-LIHC
TCGA-LUSC
TCGA-OV
TCGA-PCPG
TCGA-READ
TCGA-SKCM
TCGA-TGCT
TCGA-THYM
TCGA-UCS
TRIO-CRU,phs001163
WCDT-MCRPC,phs001648
Display
F
M
0
1000
2000
3000
4000
5000

Heatmap for Path: https://nih-ncpi.github.io/ncpi-fhir-ig-2/StructureDefinition-file-size.html

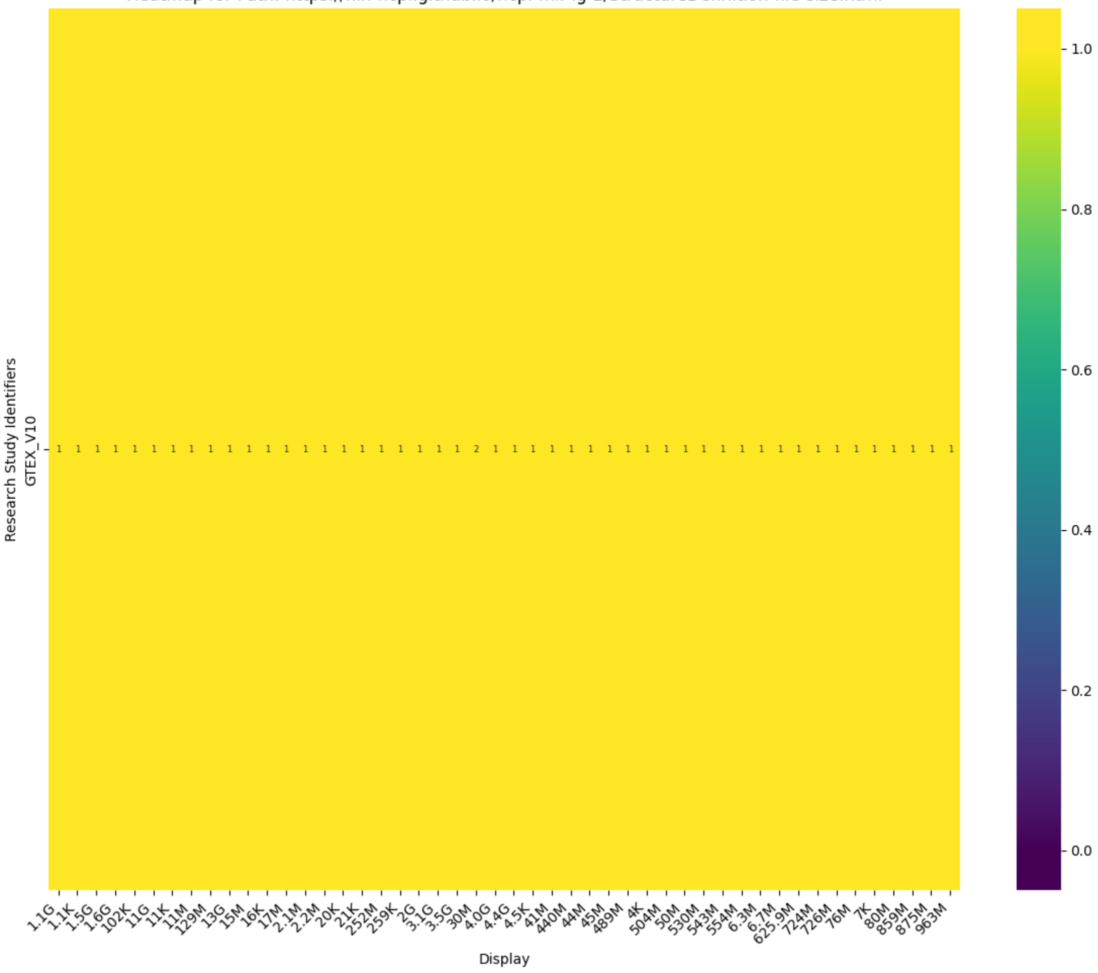

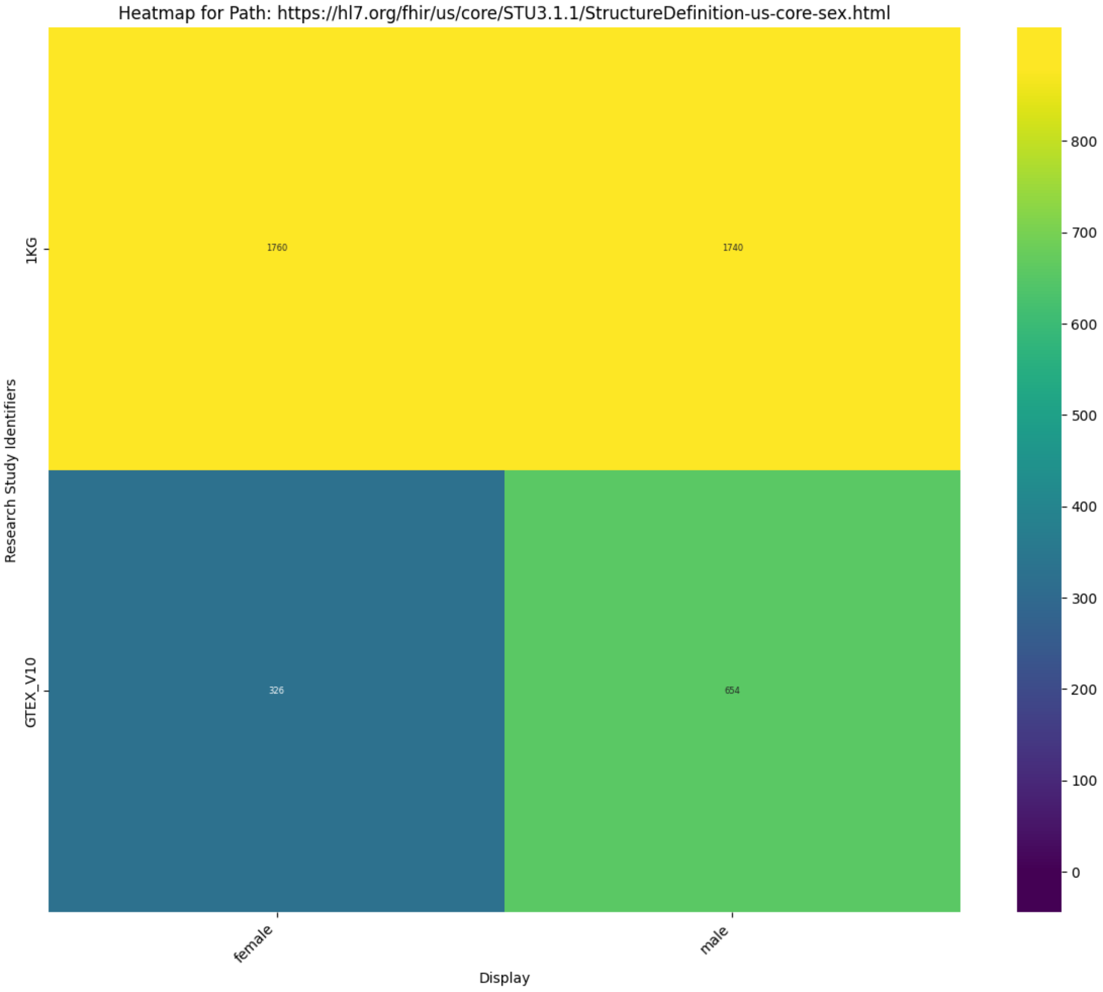

Heatmap for Path: https://hl7.org/fhir/us/core/STU3.1.1/StructureDefinition-us-core-sex.html
Research Study Identifiers
1KG
GTEX_V10
1760
1740
326
654
female
male
Display
800
700
600
500
400
300
200
100
0

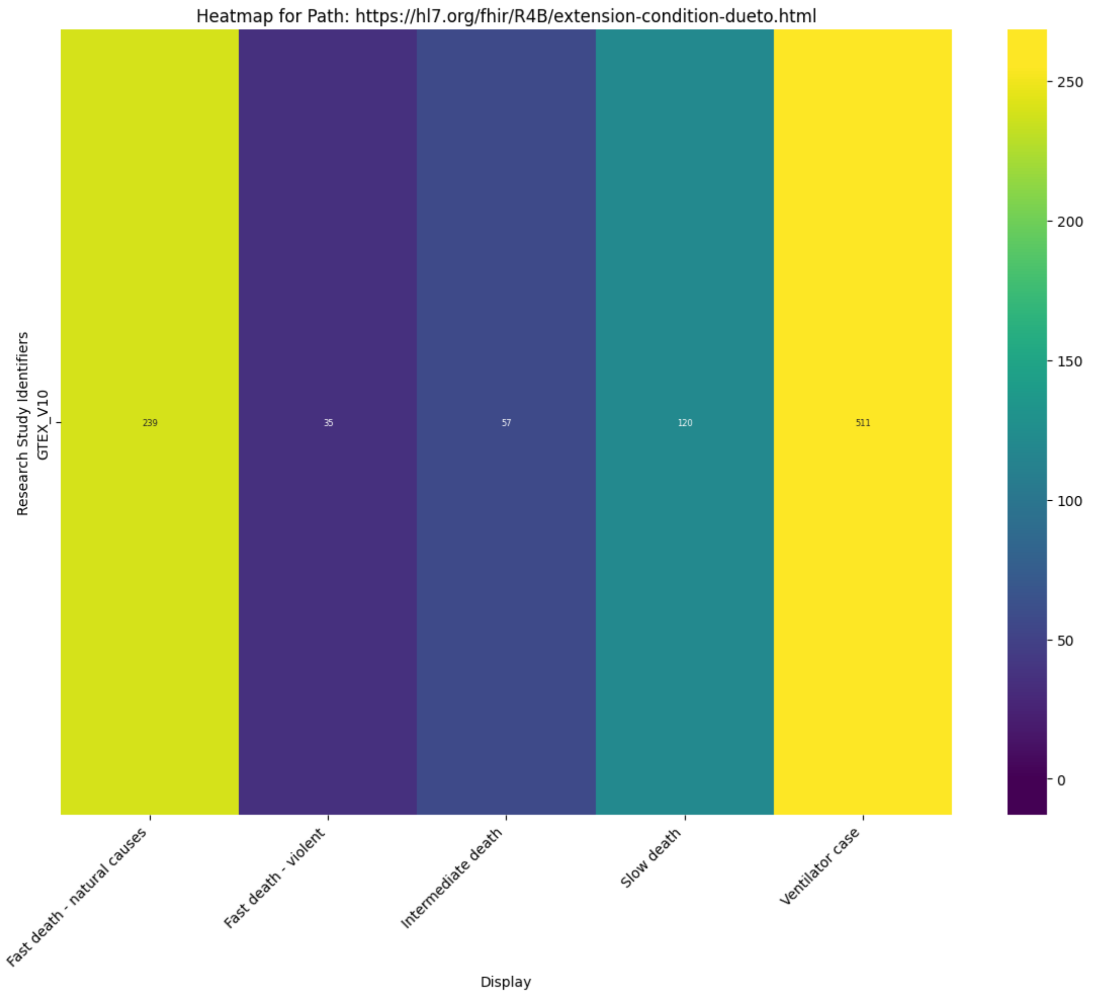
Heatmap for Path: https://hl7.org/fhir/R4B/extension-condition-dueto.html
Research Study Identifiers
GTEX_V10
239
35
57
120
511
Fast death - natural causes
Fast death - violent
Intermediate death
Slow death
Ventilator case
Display
250
200
150
100
50
0

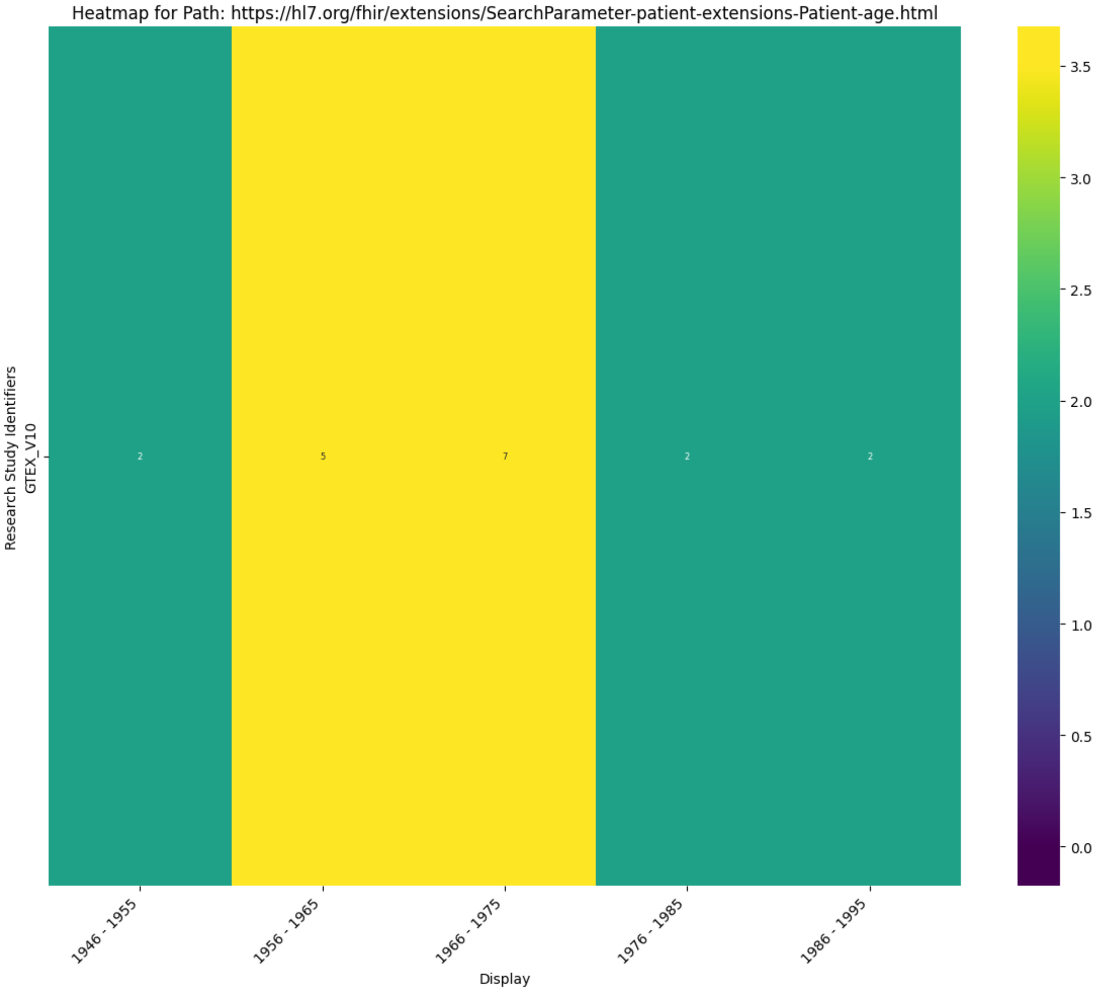
Heatmap for Path: https://hl7.org/fhir/extensions/SearchParameter-patient-extensions-Patient-age.html
Research Study Identifiers
GTEX_V10
2
5
7
2
2
1946 - 1955
1956 - 1965
1966 - 1975
1976 - 1985
1986 - 1995
Display
3.5
3.0
2.5
2.0
1.5
1.0
0.5
0.0

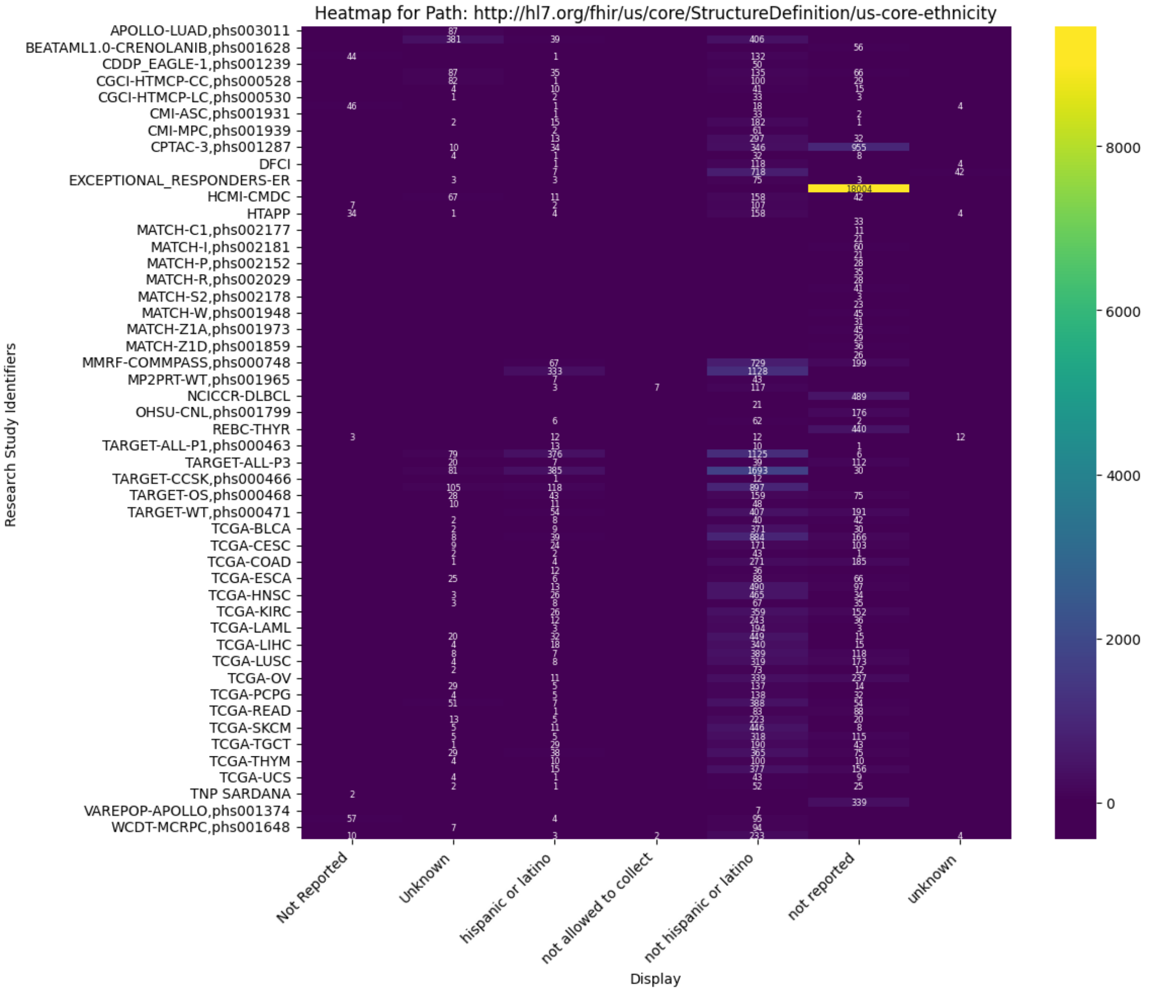

Heatmap for Path: http://hl7.org/fhir/us/core/StructureDefinition/us-core-ethnicity
Research Study Identifiers
Display
Not Reported
Unknown
hispanic or latino
not allowed to collect
not hispanic or latino
not reported
unknown
APOLLO-LUAD,phs003011
BEATAML1.0-CRENOLANIB,phs001628
CDDP_EAGLE-1,phs001239
CGCI-HTMCP-CC,phs000528
CGCI-HTMCP-LC,phs000530
CMI-ASC,phs001931
CMI-MPC,phs001939
CPTAC-3,phs001287
DFCI
EXCEPTIONAL_RESPONDERS-ER
HCMI-CMDC
HTAPP
MATCH-C1,phs002177
MATCH-I,phs002181
MATCH-P,phs002152
MATCH-R,phs002029
MATCH-S2,phs002178
MATCH-W,phs001948
MATCH-Z1A,phs001973
MATCH-Z1D,phs001859
MMRF-COMMPASS,phs000748
MP2PRT-WT,phs001965
NCICCR-DLBCL
OHSU-CNL,phs001799
REBC-THYR
TARGET-ALL-P1,phs000463
TARGET-ALL-P3
TARGET-CCSK,phs000466
TARGET-OS,phs000468
TARGET-WT,phs000471
TCGA-BLCA
TCGA-CESC
TCGA-COAD
TCGA-ESCA
TCGA-HNSC
TCGA-KIRC
TCGA-LAML
TCGA-LIHC
TCGA-LUSC
TCGA-OV
TCGA-PCPG
TCGA-READ
TCGA-SKCM
TCGA-TGCT
TCGA-THYM
TCGA-UCS
TNP SARDANA
VAREPOP-APOLLO,phs001374
WCDT-MCRPC,phs001648
8000
6000
4000
2000
0

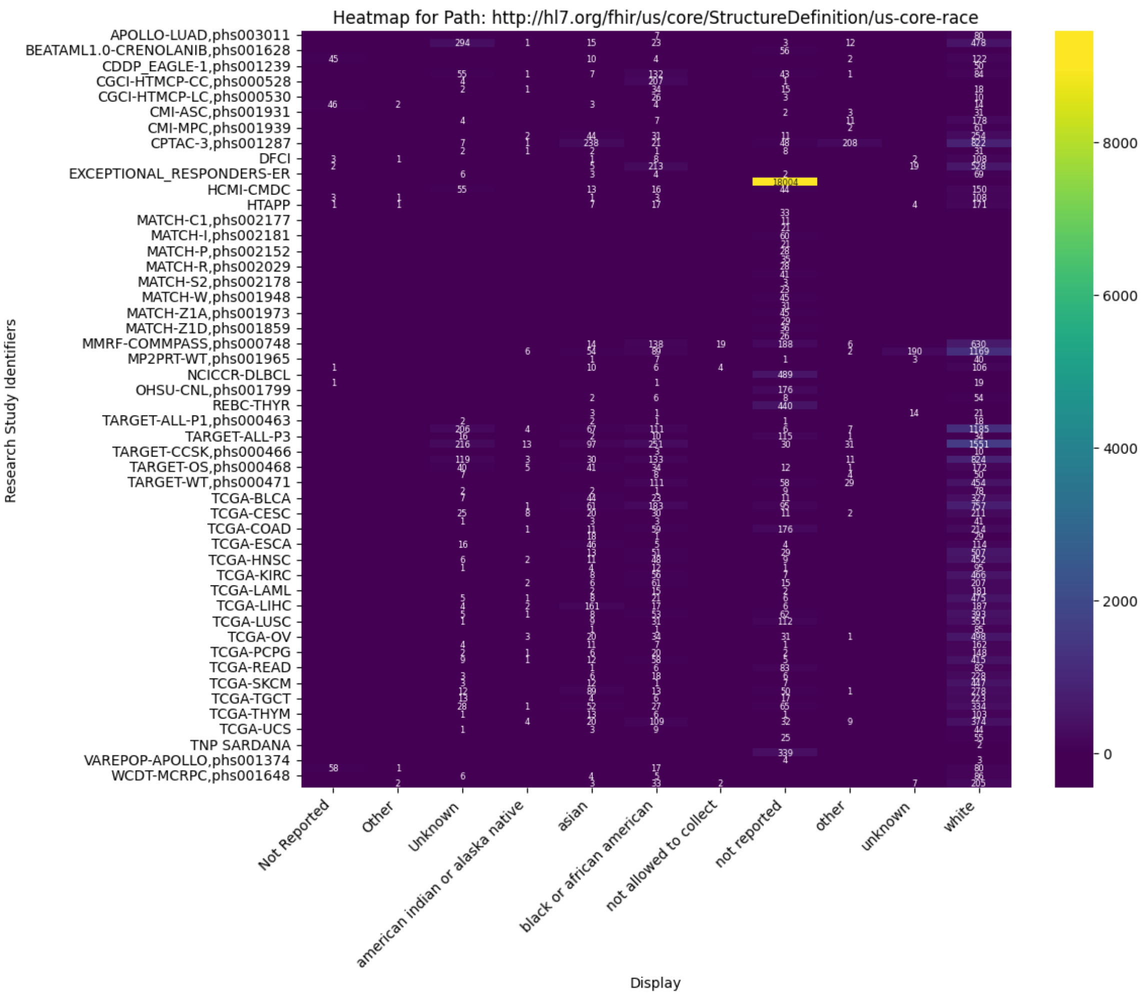
Heatmap for Path: http://hl7.org/fhir/us/core/StructureDefinition/us-core-race
Research Study Identifiers
APOLLO-LUAD,phs003011
BEATAML1.0-CRENOLANIB,phs001628
CDDP_EAGLE-1,phs001239
CGCI-HTMCP-CC,phs000528
CGCI-HTMCP-LC,phs000530
CMI-ASC,phs001931
CMI-MPC,phs001939
CPTAC-3,phs001287
DFCI
EXCEPTIONAL_RESPONDERS-ER
HCMI-CMDC
HTAPP
MATCH-C1,phs002177
MATCH-I,phs002181
MATCH-P,phs002152
MATCH-R,phs002029
MATCH-S2,phs002178
MATCH-W,phs001948
MATCH-Z1A,phs001973
MATCH-Z1D,phs001859
MMRF-COMMPASS,phs000748
MP2PRT-WT,phs001965
NCICCR-DLBCL
OHSU-CNL,phs001799
REBC-THYR
TARGET-ALL-P1,phs000463
TARGET-ALL-P3
TARGET-CCSK,phs000466
TARGET-OS,phs000468
TARGET-WT,phs000471
TCGA-BLCA
TCGA-CESC
TCGA-COAD
TCGA-ESCA
TCGA-HNSC
TCGA-KIRC
TCGA-LAML
TCGA-LIHC
TCGA-LUSC
TCGA-OV
TCGA-PCPG
TCGA-READ
TCGA-SKCM
TCGA-TGCT
TCGA-THYM
TCGA-UCS
TNP SARDANA
VAREPOP-APOLLO,phs001374
WCDT-MCRPC,phs001648
Not Reported
Other
Unknown
american indian or alaska native
asian
black or african american
not allowed to collect
not reported
other
unknown
white
Display
8000
6000
4000
2000
0

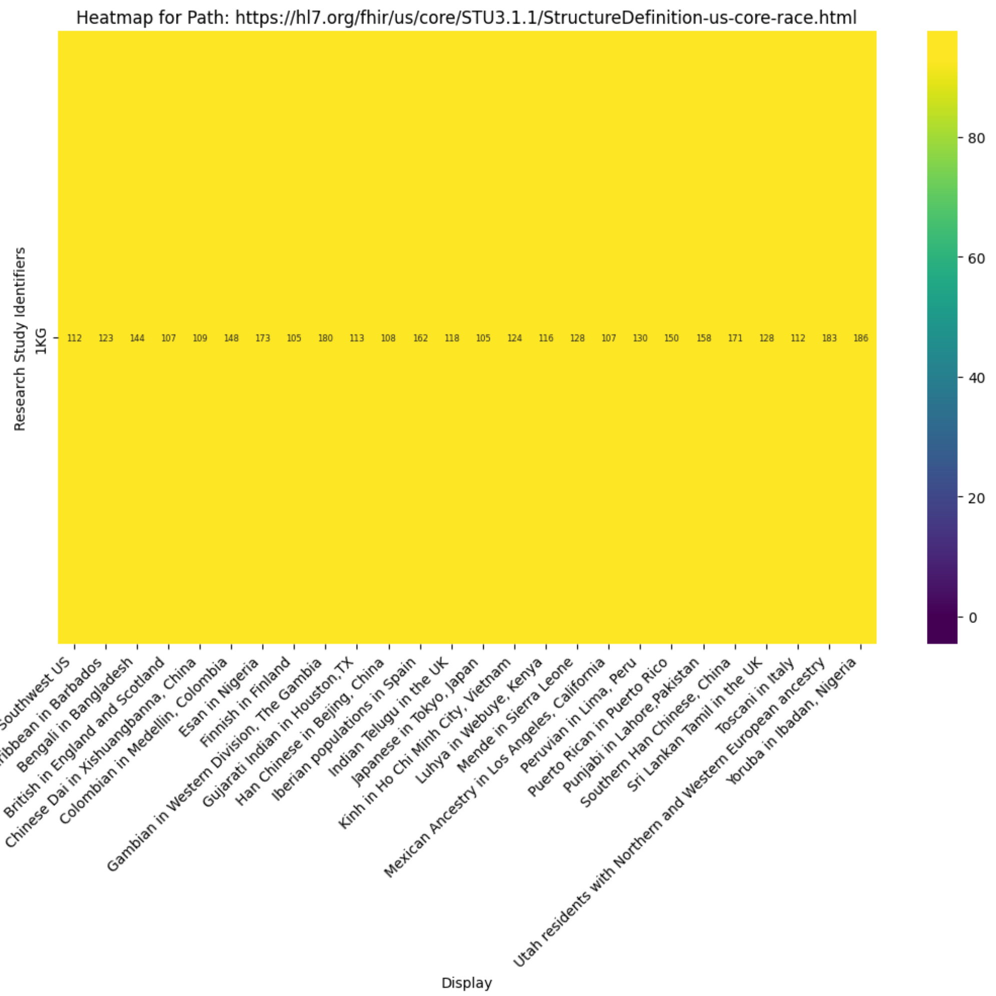
Heatmap for Path: https://hl7.org/fhir/us/core/STU3.1.1/StructureDefinition-us-core-race.html
Research Study Identifiers
1KG
112
123
144
107
109
148
173
105
180
113
108
162
118
105
124
116
128
107
130
150
158
171
128
112
183
186
80
60
40
20
0
African Ancestry in Southwest US
African Caribbean in Barbados
Bengali in Bangladesh
British in England and Scotland
Chinese Dai in Xishuangbanna, China
Colombian in Medellin, Colombia
Esan in Nigeria
Finnish in Finland
Gambian in Western Division, The Gambia
Gujarati Indian in Houston,TX
Han Chinese in Bejing, China
Iberian populations in Spain
Indian Telugu in the UK
Japanese in Tokyo, Japan
Kinh in Ho Chi Minh City, Vietnam
Luhya in Webuye, Kenya
Mende in Sierra Leone
Mexican Ancestry in Los Angeles, California
Peruvian in Lima, Peru
Puerto Rican in Puerto Rico
Punjabi in Lahore,Pakistan
Southern Han Chinese, China
Sri Lankan Tamil in the UK
Toscani in Italy
Utah residents with Northern and Western European ancestry
Yoruba in Ibadan, Nigeria
Display

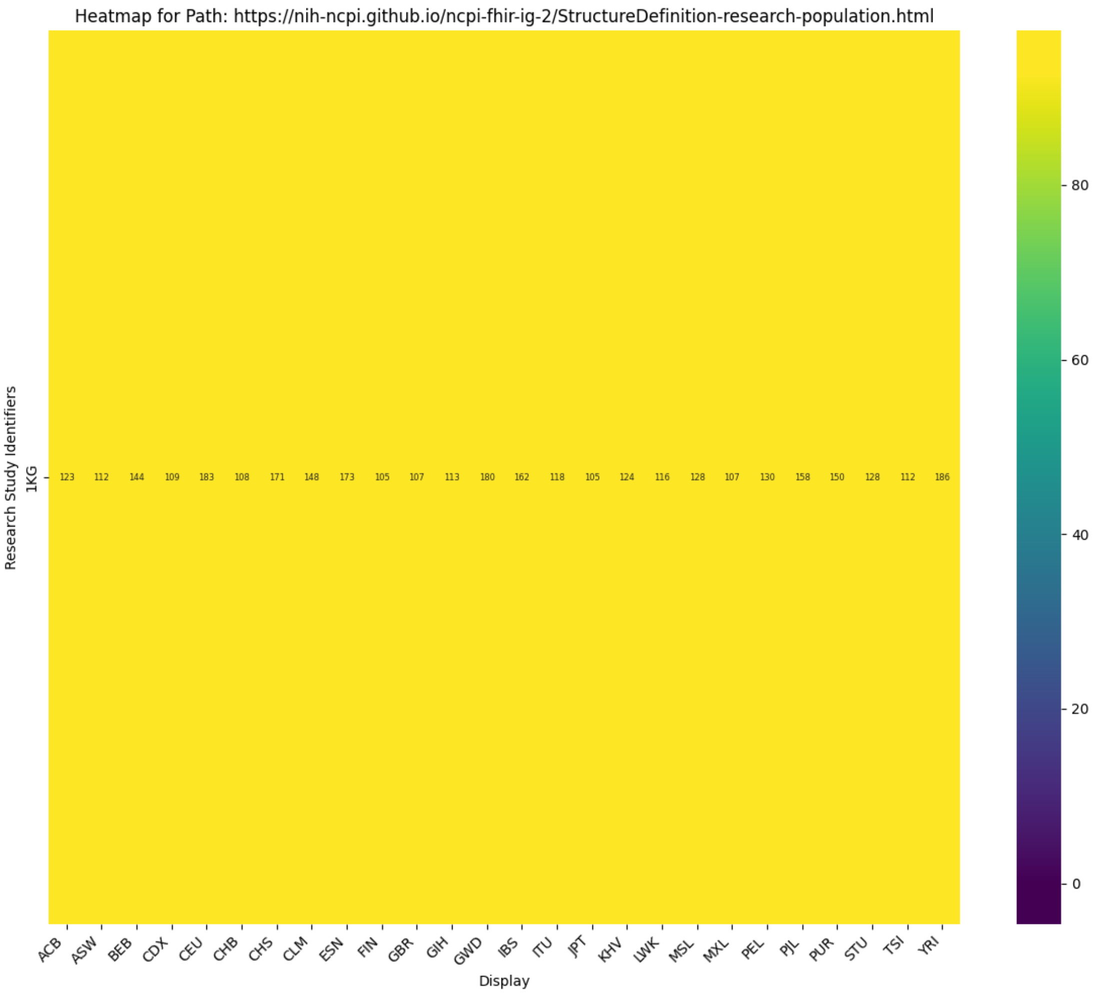


### Explore 1000 Genomes¶

Lets narrow the search to a specific Project

```
scratch_df = df[df['research_study_identifiers'].str.contains('1KG')]
scratch_df
```

| | research_study_identifiers | path | documentation | code | display | system | extension_url | count | low | high | url |
|---|---|---|---|---|---|---|---|---|---|---|---|
| **3740** | 1KG | ServiceRequest.category | https://hl7.org/fhir/R4B/servicerequest-defini... | 108252007 | Laboratory procedure | http://snomed.info/sct | | 1.0 | | | https://google-fhir.fhir-aggregator.org/Servic... |
| **3741** | 1KG | ServiceRequest.code | https://hl7.org/fhir/R4B/servicerequest-defini... | 15220000 | Laboratory test | http://snomed.info/sct | | 1.0 | | | https://google-fhir.fhir-aggregator.org/Servic... |
| **3742** | 1KG | DocumentReference.type | https://hl7.org/fhir/R4B/documentreference-def... | VCF | VCF | https://ftp.1000genomes.ebi.ac.uk/data_format | | 46.0 | | | https://google-fhir.fhir-aggregator.org/Docume... |
| **3743** | 1KG | DocumentReference.type | https://hl7.org/fhir/R4B/documentreference-def... | NEW | NEW | https://ftp.1000genomes.ebi.ac.uk/data_format | | 2.0 | | | https://google-fhir.fhir-aggregator.org/Docume... |
| **3744** | 1KG | DocumentReference.category | https://hl7.org/fhir/R4B/documentreference-def... | 1 | Chromosome 1 | https://ftp.1000genomes.ebi.ac.uk/chromosome | | 2.0 | | | https://google-fhir.fhir-aggregator.org/Docume... |
| **...** | ... | ... | ... | ... | ... | ... | ... | ... | ... | ... | ... |
| **3821** | 1KG | Patient.extension | https://hl7.org/fhir/R4B/patient-definitions.h... | LWK | LWK | | https://nih-ncpi.github.io/ncpi-fhir-ig-2/Stru... | 116.0 | | | https://google-fhir.fhir-aggregator.org/Patien... |
| **3822** | 1KG | Patient.extension | https://hl7.org/fhir/R4B/patient-definitions.h... | ASW | ASW | | https://nih-ncpi.github.io/ncpi-fhir-ig-2/Stru... | 112.0 | | | https://google-fhir.fhir-aggregator.org/Patien... |
| **3823** | 1KG | Patient.extension | https://hl7.org/fhir/R4B/patient-definitions.h... | MXL | MXL | | https://nih-ncpi.github.io/ncpi-fhir-ig-2/Stru... | 107.0 | | | https://google-fhir.fhir-aggregator.org/Patien... |
| **3824** | 1KG | Patient.extension | https://hl7.org/fhir/R4B/patient-definitions.h... | TSI | TSI | | https://nih-ncpi.github.io/ncpi-fhir-ig-2/Stru... | 112.0 | | | https://google-fhir.fhir-aggregator.org/Patien... |
| **3825** | 1KG | Patient.extension | https://hl7.org/fhir/R4B/patient-definitions.h... | GIH | GIH | | https://nih-ncpi.github.io/ncpi-fhir-ig-2/Stru... | 113.0 | | | https://google-fhir.fhir-aggregator.org/Patien... |

86 rows × 15 columns

### Explore a Condition¶

##### List the conditons found in each ResearchStudy¶

```
scratch_df = df[df['path'].str.contains('Condition.code')]
scratch_df
```

| | research_study_identifiers | path | documentation | code | display | system | extension_url | count | low | high | u |
|---|---|---|---|---|---|---|---|---|---|---|---|
| **11** | ICGC-LUCA_KR | Condition.code | https://hl7.org/fhir/R4B/condition-definitions... | C15.5 | C15.5 | https://terminology.hl7.org/5.1.0/NamingSystem... | | 400.0 | | | https://googl fhir.fhir-aggregator.o Condit... |
| **230** | TNP SARDANA | Condition.code | https://hl7.org/fhir/R4B/condition-definitions... | Mucous adenocarcinoma | Mucous adenocarcinoma | https://data.humantumoratlas.org | | 1.0 | | | https://googl fhir.fhir-aggregator.o Condit... |
| **231** | TNP SARDANA | Condition.code | https://hl7.org/fhir/R4B/condition-definitions... | Not Reported | Not Reported | https://data.humantumoratlas.org | | 1.0 | | | https://googl fhir.fhir-aggregator.o Condit... |
| **534** | HTAPP | Condition.code | https://hl7.org/fhir/R4B/condition-definitions... | Lobular and ductal carcinoma | Lobular and ductal carcinoma | https://data.humantumoratlas.org | | 9.0 | | | https://googl fhir.fhir-aggregator.o Condit... |
| **535** | HTAPP | Condition.code | https://hl7.org/fhir/R4B/condition-definitions... | Ductal carcinoma NOS | Ductal carcinoma NOS | https://data.humantumoratlas.org | | 43.0 | | | https://googl fhir.fhir-aggregator.o Condit... |
| **...** | ... | ... | ... | ... | ... | ... | ... | ... | ... | ... | ... |
| **21671** | HCMI-CMDC | Condition.code | https://hl7.org/fhir/R4B/condition-definitions... | Serous surface papillary carcinoma | Serous surface papillary carcinoma | https://gdc.cancer.gov/primary_diagnosis | | 1.0 | | | https://googl fhir.fhir-aggregator.o Condit... |
| **21672** | HCMI-CMDC | Condition.code | https://hl7.org/fhir/R4B/condition-definitions... | Carcinosarcoma, NOS | Carcinosarcoma, NOS | https://gdc.cancer.gov/primary_diagnosis | | 1.0 | | | https://googl fhir.fhir-aggregator.o Condit... |
| **21673** | HCMI-CMDC | Condition.code | https://hl7.org/fhir/R4B/condition-definitions... | Serous cystadenocarcinoma, NOS | Serous cystadenocarcinoma, NOS | https://gdc.cancer.gov/primary_diagnosis | | 1.0 | | | https://googl fhir.fhir-aggregator.o Condit... |
| **21674** | HCMI-CMDC | Condition.code | https://hl7.org/fhir/R4B/condition-definitions... | Abdominal fibromatosis | Abdominal fibromatosis | https://gdc.cancer.gov/primary_diagnosis | | 1.0 | | | https://googl fhir.fhir-aggregator.o Condit... |
| **21675** | HCMI-CMDC | Condition.code | https://hl7.org/fhir/R4B/condition-definitions... | 128660005 | Bronchio-alveolar carcinoma, mucinous | http://snomed.info/sct | | 1.0 | | | https://googl fhir.fhir-aggregator.o Condit... |

1255 rows × 15 columns

#### Find a given Condition: Neuroblastoma¶

.. FHIR Query: The curl command sends the FHIR query to the server.

2. Data Extraction and Formatting: jq extracts the system, code, and display information from the coding elements within each Condition resource and formats them as TSV.
3. Sorting: The output is piped to sort to arrange the entries alphabetically.
4. Deduplication: The -u option in sort removes any duplicate entries, leaving only unique combinations of system, code, and display.
5. Output: The final result is a sorted and deduplicated list of coding information for conditions starting with "Adenocarcinoma," presented as TSV.

```
!curl -s $FHIR_BASE'/Condition?code:text=Neuroblastoma&_count=1000&_total=accurate&_elements=subject,extension,code'  | jq -rc ' .entry[] | .resource
| .code.coding[] | [.system, .code, .display] | @tsv' | sort | uniq -c
#| jq -rc '.entry[] | .resource | [.subject.reference, (.extension[] | .valueReference.reference)] '
```

```
      2 https://data.humantumoratlas.org        Central neuroblastoma   Central neuroblastoma
      3 https://data.humantumoratlas.org        Neuroblastoma NOS       Neuroblastoma NOS
    995 https://gdc.cancer.gov/primary_diagnosis        Neuroblastoma, NOS      Neuroblastoma, NOS
```

```
! pip install fhir-aggregator-client --no-cache-dir --quiet --no-warn-script-location 2>/dev/null
! pip install lifelines --no-cache-dir --quiet --no-warn-script-location 2>/dev/null
! pip freeze | grep fhir_aggregator_client
```

```
fhir_aggregator_client==0.2.2
```

```
%env FHIR_BASE= https://google-fhir.fhir-aggregator.org/
```

```
env: FHIR_BASE=https://google-fhir.fhir-aggregator.org/
```

```
! rm -rf /root/.fhir-aggregator
!fq run cholangiocarcinoma-graph '/Condition?code:code=70179006'
```

```
Creating directory: /root/.fhir-aggregator
cholangiocarcinoma-graph is valid FHIR R5 GraphDefinition
ℹ Fetching https://google-fhir.fhir-aggregator.org/Condition?code:code=70179006
ℹ Processing Condition with 389 resources
ℹ Processing 1 links for Condition in parallel.
ℹ Processing link: Patient/_id={ref} with 354 Condition(s)
✔ Processed link: Patient/_id={ref}
ℹ Processing Patient with 389 resources
ℹ Processing 10 links for Patient in parallel.
ℹ Processing link: ResearchSubject/individual={ref}&_include=ResearchSubject:study with 354 Patient(s)
ℹ Processing link: Group/member={ref} with 354 Patient(s)
ℹ Processing link: Specimen/subject={ref} with 354 Patient(s)
ℹ Processing link: Observation/subject={ref}&code=NCIT_C156418,NCIT_C156419,NCIT_C164934&_count=1000&_total=accurate with 354 Patient(s)
ℹ Processing link: Procedure/subject={ref} with 354 Patient(s)
ℹ Processing link: DocumentReference/subject={ref}&_count=1000&_total=accurate with 354 Patient(s)
✖ Could not find any resources for Patient->Group link: {'params': 'member={ref}&_count=1000&_total=accurate', 'sourceId': 'Patient', 'targetId': 'Group',
'path': 'Patient.id'}
ℹ Processing link: ImagingStudy/subject={ref}&_count=1000&_total=accurate with 354 Patient(s)
ℹ Processing link: MedicationAdministration/subject={ref}&_count=1000&_total=accurate with 354 Patient(s)
ℹ Processing link: Encounter/subject={ref}&_count=1000&_total=accurate with 354 Patient(s)
✔ Processed link: ResearchSubject/individual={ref}&_include=ResearchSubject:study
✔ Processed link: Encounter/subject={ref}&_count=1000&_total=accurate
✔ Processed link: Group/member={ref}
✔ Processed link: MedicationAdministration/subject={ref}&_count=1000&_total=accurate
✔ Processed link: Procedure/subject={ref}
✔ Processed link: DocumentReference/subject={ref}&_count=1000&_total=accurate
✔ Processed link: ImagingStudy/subject={ref}&_count=1000&_total=accurate
✔ Processed link: Specimen/subject={ref}
✔ Processed link: Observation/subject={ref}&code=NCIT_C156418,NCIT_C156419,NCIT_C164934&_count=1000&_total=accurate
ℹ Processing ResearchSubject with 354 resources
ℹ Processing Specimen with 411 resources
ℹ Processing ResearchStudy with 1418 resources
ℹ Processing 1 links for ResearchSubject in parallel.
ℹ Processing 1 links for Specimen in parallel.
ℹ Processing 1 links for ResearchStudy in parallel.
✔ Processed link: ResearchStudy/
ℹ Processing link: ServiceRequest/specimen={ref} with 1418 Specimen(s)
ℹ Processing link: DocumentReference/subject={ref}&_count=1000&_total=accurate with 18 ResearchStudy(s)
✔ Processed link: DocumentReference/subject={ref}&_count=1000&_total=accurate
✔ Processed link: ServiceRequest/specimen={ref}
Aggregated Results: {'Condition': 389, 'Observation': 7257, 'Patient': 354, 'ResearchStudy': 18, 'ResearchSubject': 411, 'Specimen': 1418}
database available at: /root/.fhir-aggregator/fhir-graph.sqlite
```

```
!fq results visualize
```

```
Wrote: fhir-graph.html
```

```
from IPython.display import HTML
with open('fhir-graph.html', 'r') as file:
    html_content = file.read()
display(HTML("<div style='height: 800px;'>{}</div>".format(html_content)))
```

```
!fq results dataframe
```

```
Saved fhir-graph.tsv
```

```
import sqlite3
import pandas as pd

path = "/root/.fhir-aggregator/fhir-graph.sqlite"
conn = sqlite3.connect(path)

df_raw = pd.read_sql_query("SELECT * FROM resources", conn)

conn.close()
df_raw
```

| | id | resource_type | key | resource |
|---|---|---|---|---|
| **0** | f1f5d536-ec6a-565b-9bb4-e7a459e47de3 | Condition | Condition/f1f5d536-ec6a-565b-9bb4-e7a459e47de3 | {"category": [{"coding": [{"code": "encounter-... |
| **1** | bcc646fb-6b16-50e6-bff0-9850431eb764 | Condition | Condition/bcc646fb-6b16-50e6-bff0-9850431eb764 | {"category": [{"coding": [{"code": "encounter-... |
| **2** | 8fcb93f3-9687-5603-94fa-3dce12f747b8 | Condition | Condition/8fcb93f3-9687-5603-94fa-3dce12f747b8 | {"category": [{"coding": [{"code": "encounter-... |
| **3** | c1a0b7bb-13e1-5243-94e3-55f013df77de | Condition | Condition/c1a0b7bb-13e1-5243-94e3-55f013df77de | {"category": [{"coding": [{"code": "encounter-... |
| **4** | 709b9fe4-373d-5620-b778-e4d5a633bc55 | Condition | Condition/709b9fe4-373d-5620-b778-e4d5a633bc55 | {"category": [{"coding": [{"code": "encounter-... |
| **...** | ... | ... | ... | ... |
| **9842** | a1791515-0e91-5107-8b5b-712bb8d9dbf0 | Observation | Observation/a1791515-0e91-5107-8b5b-712bb8d9dbf0 | {"category": [{"coding": [{"code": "laboratory... |
| **9843** | 04c5e0e2-f8b8-584a-99e7-cfd56b144a87 | Observation | Observation/04c5e0e2-f8b8-584a-99e7-cfd56b144a87 | {"category": [{"coding": [{"code": "laboratory... |
| **9844** | 60a9d4e9-9653-56e2-baf6-3645af7f49c8 | Observation | Observation/60a9d4e9-9653-56e2-baf6-3645af7f49c8 | {"category": [{"coding": [{"code": "laboratory... |
| **9845** | 0d3524a9-db47-5c86-bbfd-09e80a1db41a | Observation | Observation/0d3524a9-db47-5c86-bbfd-09e80a1db41a | {"category": [{"coding": [{"code": "laboratory... |
| **9846** | 0eeb3627-38c1-529e-aeb3-dcf2e632ae45 | Observation | Observation/0eeb3627-38c1-529e-aeb3-dcf2e632ae45 | {"category": [{"coding": [{"code": "laboratory... |

9847 rows × 4 columns

```
import orjson

research_studies = df_raw[df_raw.resource_type.isin(["ResearchStudy"])]

titles = [orjson.loads(resource)['title'] for resource in research_studies.resource]
print(titles)
```

```
['FM-AD', 'MATCH-U', 'MATCH-Z1A', 'MATCH-H', 'tcga_chol', 'tcga_lihc', 'phs002431', 'MATCH-B', 'EXCEPTIONAL_RESPONDERS-ER', 'MATCH-W', 'MATCH-Z1D',
'tcga_uvm', 'MATCH-S1', 'MATCH-I', 'HCMI-CMDC', 'MATCH-C1', 'MATCH-Q', 'MATCH-P']
```

```
import json
import pandas as pd
import numpy as np
import matplotlib.pyplot as plt
import seaborn as sns
import warnings
warnings.filterwarnings('ignore')

plt.style.use('default')
sns.set_palette("husl")

# create resource_dfs from df_raw
resource_dfs = {}
```

```
for resource_type in df_raw['resource_type'].unique():
    resource_dfs[resource_type] = df_raw[df_raw['resource_type'] == resource_type].copy()

# extract and create clean datasets
print("extracting datasets for summary plots...")

# 1. patient demographics
patients = []
df_patient = resource_dfs['Patient']
for i, row in df_patient.iterrows():
    try:
        patient_json = json.loads(row['resource'])
        patient_data = {'patient_id': patient_json.get('id')}

        if 'extension' in patient_json:
            for ext in patient_json['extension']:
                if 'url' in ext:
                    if 'us-core-birthsex' in ext['url']:
                        patient_data['gender'] = ext.get('valueCode')
                    elif 'us-core-race' in ext['url']:
                        patient_data['race'] = ext.get('valueCode', ext.get('valueString'))
                    elif 'us-core-ethnicity' in ext['url']:
                        patient_data['ethnicity'] = ext.get('valueCode', ext.get('valueString'))
        patients.append(patient_data)
    except:
        continue
df_patients = pd.DataFrame(patients)

# 2. research studies and patient assignments
studies = []
df_research_study = resource_dfs['ResearchStudy']
for i, row in df_research_study.iterrows():
    try:
        study_json = json.loads(row['resource'])
        studies.append({
            'study_id': study_json.get('id'),
            'study_title': study_json.get('title', 'unknown'),
            'study_status': study_json.get('status', 'unknown')
        })
    except:
        continue
df_studies = pd.DataFrame(studies)

patient_studies = []
df_research_subject = resource_dfs['ResearchSubject']
for i, row in df_research_subject.iterrows():
    try:
        subject_json = json.loads(row['resource'])
        if ('individual' in subject_json and 'reference' in subject_json['individual'] and
            'study' in subject_json and 'reference' in subject_json['study']):
            patient_studies.append({
                'patient_id': subject_json['individual']['reference'].replace('Patient/', ''),
                'study_id': subject_json['study']['reference'].replace('ResearchStudy/', '')
            })
    except:
        continue
df_patient_studies = pd.DataFrame(patient_studies)

# 3. specimens
specimens = []
df_specimen = resource_dfs['Specimen']
```

```python
for i, row in df_specimen.iterrows():
    try:
        specimen_json = json.loads(row['resource'])
        specimen_data = {
            'specimen_id': specimen_json.get('id'),
            'specimen_type': 'unknown'
        }

        if 'type' in specimen_json and 'coding' in specimen_json['type']:
            if len(specimen_json['type']['coding']) > 0:
                specimen_data['specimen_type'] = specimen_json['type']['coding'][0].get('display', 'unknown')

        if 'subject' in specimen_json and 'reference' in specimen_json['subject']:
            specimen_data['patient_id'] = specimen_json['subject']['reference'].replace('Patient/', '')

        specimens.append(specimen_data)
    except:
        continue
df_specimens = pd.DataFrame(specimens)

# 4. genetic variants
variants = []
df_obs = resource_dfs['Observation']
for i, row in df_obs.iterrows():
    try:
        obs_json = json.loads(row['resource'])

        patient_id = None
        if 'subject' in obs_json and 'reference' in obs_json['subject']:
            patient_id = obs_json['subject']['reference'].replace('Patient/', '')

        if patient_id:
            obs_type = None
            if 'code' in obs_json and 'coding' in obs_json['code']:
                if len(obs_json['code']['coding']) > 0:
                    obs_type = obs_json['code']['coding'][0].get('display')

            if obs_type == 'Performed Genetic Observation Result Mutation Type Code':
                variant_data = {
                    'patient_id': patient_id,
                    'variant_class': None,
                    'hugo_symbol': None,
                    'project': None
                }

                if 'component' in obs_json:
                    for comp in obs_json['component']:
                        if 'code' in comp and 'valueString' in comp:
                            comp_code = None
                            if 'coding' in comp['code'] and len(comp['code']['coding']) > 0:
                                comp_code = comp['code']['coding'][0].get('display')

                            comp_value = comp['valueString']

                            if comp_code == 'variant_class':
                                variant_data['variant_class'] = comp_value
                            elif comp_code == 'hugo_symbol':
                                variant_data['hugo_symbol'] = comp_value
                            elif comp_code == 'project_short_name':
                                variant_data['project'] = comp_value
```

```
                variants.append(variant_data)
    except:
        continue
df_variants = pd.DataFrame(variants)

# merge key datasets
df_main = df_patient_studies.merge(df_patients, on='patient_id', how='left')
df_main = df_main.merge(df_studies, on='study_id', how='left')

# add specimen counts
specimen_summary = df_specimens.groupby(['patient_id', 'specimen_type']).size().unstack(fill_value=0)
if 'Normal' in specimen_summary.columns:
    specimen_summary['normal_specimens'] = specimen_summary['Normal']
if 'Tumor' in specimen_summary.columns:
    specimen_summary['tumor_specimens'] = specimen_summary['Tumor']
specimen_summary['total_specimens'] = specimen_summary.sum(axis=1)
df_main = df_main.merge(specimen_summary[['normal_specimens', 'tumor_specimens', 'total_specimens']],
                       on='patient_id', how='left')
df_main[['normal_specimens', 'tumor_specimens', 'total_specimens']] = df_main[['normal_specimens', 'tumor_specimens', 'total_specimens']].fillna(0)

# add variant counts
variant_summary = df_variants.groupby('patient_id').size().reset_index(name='total_variants')
df_main = df_main.merge(variant_summary, on='patient_id', how='left')
df_main['total_variants'] = df_main['total_variants'].fillna(0)

print(f"final dataset: {len(df_main)} patients")

# create summary plots
fig, axes = plt.subplots(2, 3, figsize=(18, 12))
axes = axes.flatten()

# plot 1: gender distribution
if 'gender' in df_main.columns and df_main['gender'].notna().any():
    gender_counts = df_main['gender'].value_counts()
    colors = ['lightblue', 'pink'][:len(gender_counts)]
    gender_counts.plot(kind='pie', ax=axes[0], autopct='%1.1f%%', colors=colors, startangle=90)
    axes[0].set_title(f'gender distribution\n(n={gender_counts.sum()})', fontweight='bold')
    axes[0].set_ylabel('')
else:
    axes[0].text(0.5, 0.5, 'no gender data', ha='center', va='center', transform=axes[0].transAxes)
    axes[0].set_title('gender distribution')

# plot 2: research studies
if 'study_title' in df_main.columns and df_main['study_title'].notna().any():
    study_counts = df_main['study_title'].value_counts().head(10)
    study_counts.plot(kind='barh', ax=axes[1], color='lightgreen')
    axes[1].set_title('research studies (top 10)', fontweight='bold')
    axes[1].set_xlabel('patient count')
    for i, v in enumerate(study_counts.values):
        axes[1].text(v + max(study_counts.values) * 0.01, i, str(v), va='center', fontweight='bold')
else:
    axes[1].text(0.5, 0.5, 'no study data', ha='center', va='center', transform=axes[1].transAxes)
    axes[1].set_title('research studies')

# plot 3: specimen types
specimen_data = df_specimens['specimen_type'].value_counts()
if not specimen_data.empty:
    colors = plt.cm.Set3(np.linspace(0, 1, len(specimen_data)))
    specimen_data.plot(kind='bar', ax=axes[2], color=colors, alpha=0.8)
    axes[2].set_title('specimen type distribution', fontweight='bold')
    axes[2].set_ylabel('count')
```

```
    axes[2].tick_params(axis='x', rotation=45)
    for i, v in enumerate(specimen_data.values):
        axes[2].text(i, v + max(specimen_data.values) * 0.01, str(v), ha='center', va='bottom', fontweight='bold')
else:
    axes[2].text(0.5, 0.5, 'no specimen data', ha='center', va='center', transform=axes[2].transAxes)
    axes[2].set_title('specimen types')

# plot 4: patients with genetic data
if 'total_variants' in df_main.columns:
    genetic_status = (df_main['total_variants'] > 0).value_counts()
    genetic_status.index = ['no variants', 'has variants']
    genetic_status.plot(kind='pie', ax=axes[3], autopct='%1.1f%%',
                       colors=['lightcoral', 'lightsteelblue'], startangle=90)
    axes[3].set_title('patients with genetic variants', fontweight='bold')
    axes[3].set_ylabel('')
else:
    axes[3].text(0.5, 0.5, 'no variant data', ha='center', va='center', transform=axes[3].transAxes)
    axes[3].set_title('genetic variants')

# plot 5: variant classes
if not df_variants.empty and 'variant_class' in df_variants.columns:
    variant_classes = df_variants['variant_class'].value_counts().head(8)
    colors = plt.cm.viridis(np.linspace(0, 1, len(variant_classes)))
    variant_classes.plot(kind='bar', ax=axes[4], color=colors, alpha=0.8)
    axes[4].set_title('genetic variant classes', fontweight='bold')
    axes[4].set_ylabel('count')
    axes[4].tick_params(axis='x', rotation=45)
    for i, v in enumerate(variant_classes.values):
        axes[4].text(i, v + max(variant_classes.values) * 0.01, str(v), ha='center', va='bottom', fontweight='bold')
else:
    axes[4].text(0.5, 0.5, 'no variant classes', ha='center', va='center', transform=axes[4].transAxes)
    axes[4].set_title('variant classes')

# plot 6: top mutated genes
if not df_variants.empty and 'hugo_symbol' in df_variants.columns and df_variants['hugo_symbol'].notna().any():
    top_genes = df_variants['hugo_symbol'].value_counts().head(10)
    colors = plt.cm.plasma(np.linspace(0, 1, len(top_genes)))
    top_genes.plot(kind='bar', ax=axes[5], color=colors, alpha=0.8)
    axes[5].set_title('most frequently mutated genes', fontweight='bold')
    axes[5].set_ylabel('mutation count')
    axes[5].tick_params(axis='x', rotation=45)
    for i, v in enumerate(top_genes.values):
        axes[5].text(i, v + max(top_genes.values) * 0.01, str(v), ha='center', va='bottom', fontweight='bold')
else:
    axes[5].text(0.5, 0.5, 'no gene symbols\navailable', ha='center', va='center', transform=axes[5].transAxes)
    axes[5].set_title('mutated genes')

plt.tight_layout()
plt.show()

# print summary statistics
print(f"\ndataset summary:")
print(f"patients: {len(df_patients)}")
print(f"specimens: {len(df_specimens)} ({df_specimens['patient_id'].nunique()} patients)")
print(f"genetic variants: {len(df_variants)} ({df_variants['patient_id'].nunique()} patients)")
print(f"research studies: {len(df_studies)}")
print(f"patient-study assignments: {len(df_patient_studies)}")

if 'gender' in df_patients.columns:
    print(f"gender: {df_patients['gender'].value_counts().to_dict()}")
```

```
if not df_specimens.empty:
    print(f"specimen types: {df_specimens['specimen_type'].value_counts().to_dict()}")

if not df_variants.empty and 'variant_class' in df_variants.columns:
    print(f"variant classes: {df_variants['variant_class'].value_counts().to_dict()}")

extracting datasets for summary plots...
final dataset: 411 patients
```

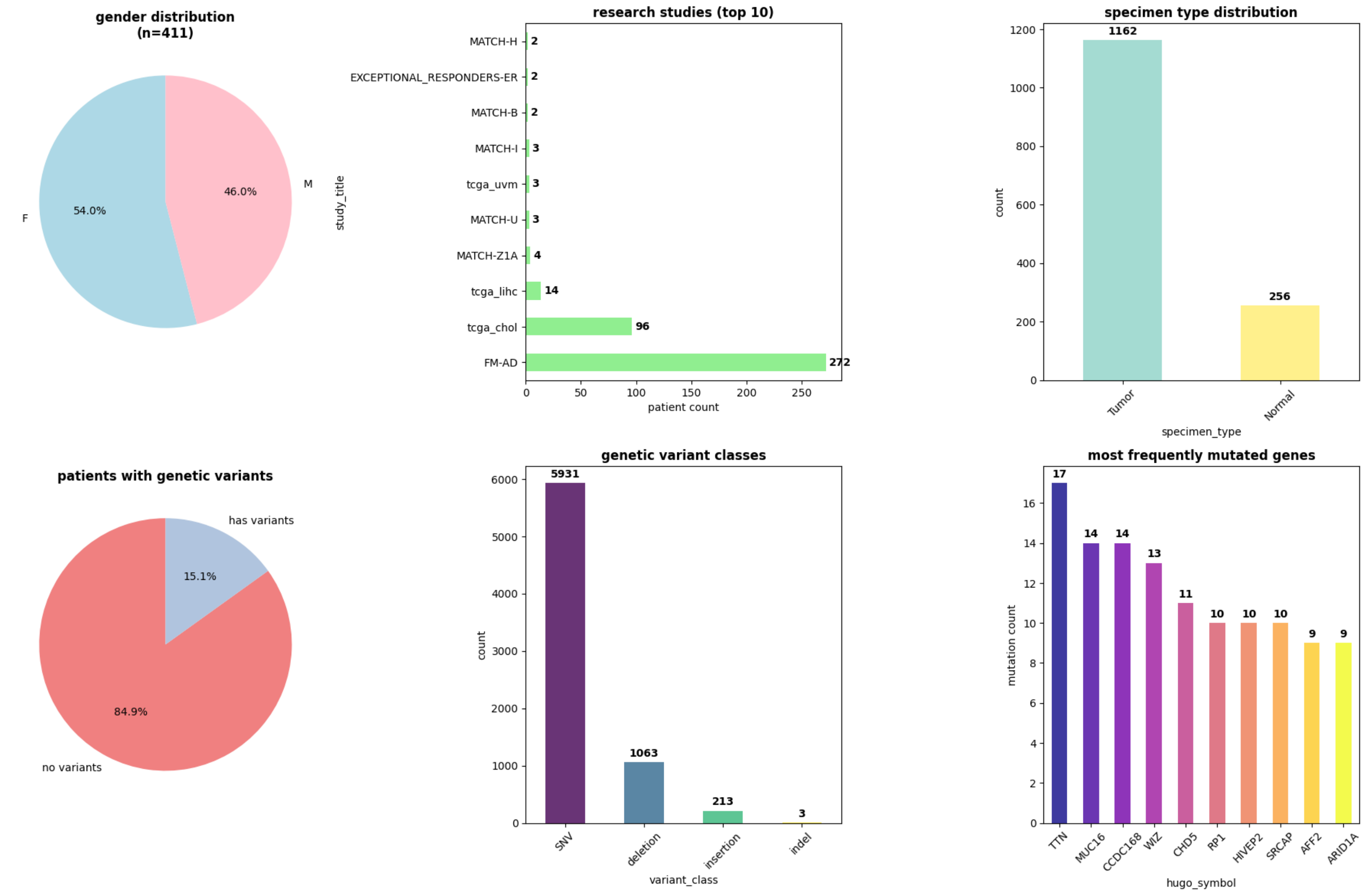


```
dataset summary:
patients: 354
specimens: 1418 (154 patients)
genetic variants: 7210 (31 patients)
research studies: 18
patient-study assignments: 411
gender: {'F': 191, 'M': 163}
specimen types: {'Tumor': 1162, 'Normal': 256}
variant classes: {'SNV': 5931, 'deletion': 1063, 'insertion': 213, 'indel': 3}
```

```python
import pandas as pd
import numpy as np
import matplotlib.pyplot as plt
from lifelines import KaplanMeierFitter
import seaborn as sns
import warnings
warnings.filterwarnings('ignore')

plt.style.use('default')
sns.set_palette("husl")

print("extracting temporal data for kaplan-meier analysis...")

# extract temporal observations
temporal_data = []
df_obs = resource_dfs['Observation']

for i, row in df_obs.iterrows():
    try:
        obs_json = json.loads(row['resource'])

        patient_id = None
        if 'subject' in obs_json and 'reference' in obs_json['subject']:
            patient_id = obs_json['subject']['reference'].replace('Patient/', '')

        if patient_id:
            obs_type = None
            if 'code' in obs_json and 'coding' in obs_json['code']:
                if len(obs_json['code']['coding']) > 0:
                    obs_type = obs_json['code']['coding'][0].get('display')

            # extract temporal values
            if obs_type in ['Days between Diagnosis and Death', 'Days Between Birth and Diagnosis']:
                if 'valueQuantity' in obs_json and 'value' in obs_json['valueQuantity']:
                    temporal_data.append({
                        'patient_id': patient_id,
                        'obs_type': obs_type,
                        'days': obs_json['valueQuantity']['value']
                    })
    except:
        continue

df_temporal = pd.DataFrame(temporal_data)
print(f"extracted {len(df_temporal)} temporal observations")

# prepare survival data
survival_data = []
diagnosis_death = df_temporal[df_temporal['obs_type'] == 'Days between Diagnosis and Death']

if not diagnosis_death.empty:
    print(f"found {len(diagnosis_death)} diagnosis-to-death observations")

    # merge with patient data to get demographics and study info
    df_survival_base = df_main.merge(diagnosis_death[['patient_id', 'days']], on='patient_id', how='inner')
    df_survival_base = df_survival_base.rename(columns={'days': 'survival_days'})
    df_survival_base['survival_years'] = df_survival_base['survival_days'] / 365.25

    # create event indicator (1 = death observed, 0 = censored)
    # since we have "days to death", these are all death events
    df_survival_base['event'] = 1
```

```python
print(f"survival analysis dataset: {len(df_survival_base)} patients")
print(f"mean survival: {df_survival_base['survival_years'].mean():.1f} years")
print(f"median survival: {df_survival_base['survival_years'].median():.1f} years")

# create kaplan-meier plots - show all groups
fig, axes = plt.subplots(1, 3, figsize=(18, 6))

# plot 1: survival by gender (all genders)
ax = axes[0]
if 'gender' in df_survival_base.columns and df_survival_base['gender'].notna().sum() > 0:
    gender_groups = df_survival_base['gender'].value_counts()
    colors = plt.cm.Set1(np.linspace(0, 1, len(gender_groups)))

    for i, (gender, color) in enumerate(zip(gender_groups.index, colors)):
        if gender_groups[gender] >= 1:  # show all groups with at least 1 patient
            gender_data = df_survival_base[df_survival_base['gender'] == gender]
            kmf.fit(gender_data['survival_years'], gender_data['event'],
                   label=f'{gender} (n={len(gender_data)})')
            kmf.plot_survival_function(ax=ax, color=color, linewidth=3)

    ax.set_title('survival by gender', fontweight='bold', fontsize=14)
    ax.set_xlabel('years from diagnosis')
    ax.set_ylabel('survival probability')
    ax.grid(True, alpha=0.3)
    ax.legend()
else:
    ax.text(0.5, 0.5, 'no gender data', ha='center', va='center', transform=ax.transAxes)
    ax.set_title('survival by gender')

# plot 2: survival by race (all races)
ax = axes[1]
if 'race' in df_survival_base.columns and df_survival_base['race'].notna().sum() > 0:
    race_groups = df_survival_base['race'].value_counts()
    colors = plt.cm.Set2(np.linspace(0, 1, len(race_groups)))

    for i, (race, color) in enumerate(zip(race_groups.index, colors)):
        if race_groups[race] >= 1:  # show all races with at least 1 patient
            race_data = df_survival_base[df_survival_base['race'] == race]
            kmf.fit(race_data['survival_years'], race_data['event'],
                   label=f'{race} (n={len(race_data)})')
            kmf.plot_survival_function(ax=ax, color=color, linewidth=3)

    ax.set_title('survival by race', fontweight='bold', fontsize=14)
    ax.set_xlabel('years from diagnosis')
    ax.set_ylabel('survival probability')
    ax.grid(True, alpha=0.3)
    ax.legend()

elif 'ethnicity' in df_survival_base.columns and df_survival_base['ethnicity'].notna().sum() > 0:
    # fallback to ethnicity if no race data
    eth_groups = df_survival_base['ethnicity'].value_counts()
    colors = plt.cm.Set2(np.linspace(0, 1, len(eth_groups)))

    for i, (ethnicity, color) in enumerate(zip(eth_groups.index, colors)):
        if eth_groups[ethnicity] >= 1:
            eth_data = df_survival_base[df_survival_base['ethnicity'] == ethnicity]
            kmf.fit(eth_data['survival_years'], eth_data['event'],
                   label=f'{ethnicity} (n={len(eth_data)})')
            kmf.plot_survival_function(ax=ax, color=color, linewidth=3)

    ax.set_title('survival by ethnicity', fontweight='bold', fontsize=14)
```

```python
    ax.set_xlabel('years from diagnosis')
    ax.set_ylabel('survival probability')
    ax.grid(True, alpha=0.3)
    ax.legend()
else:
    ax.text(0.5, 0.5, 'no race/ethnicity data', ha='center', va='center', transform=ax.transAxes)
    ax.set_title('survival by race')

# plot 3: survival by variant class (SNV vs deletion vs others)
ax = axes[2]

# merge with variant data to get variant classes
df_survival_variants = df_survival_base.merge(
    df_variants[df_variants['variant_class'].notna()][['patient_id', 'variant_class']],
    on='patient_id', how='left'
)

# debug variant class distribution
print(f"\ndebugging variant class data:")
print(f"survival patients: {len(df_survival_base)}")

if not df_variants.empty and 'variant_class' in df_variants.columns:
    variant_class_counts = df_variants['variant_class'].value_counts()
    print(f"variant classes in all data: {variant_class_counts.to_dict()}")

    # check how many survival patients have each variant class
    survival_variant_counts = df_survival_variants['variant_class'].value_counts()
    print(f"variant classes in survival cohort: {survival_variant_counts.to_dict()}")

    # create broader groups for better sample sizes
    def categorize_variant(variant_class):
        if pd.isna(variant_class):
            return 'no mutations'
        elif variant_class == 'SNV':
            return 'SNV'
        elif variant_class in ['deletion', 'insertion', 'indel']:
            return 'indel/deletion'
        else:
            return 'other mutations'

    df_survival_variants['mutation_category'] = df_survival_variants['variant_class'].apply(categorize_variant)

    mutation_groups = df_survival_variants['mutation_category'].value_counts()
    print(f"mutation categories in survival cohort: {mutation_groups.to_dict()}")

    # plot survival curves for each category with sufficient patients
    colors = plt.cm.Set1(np.linspace(0, 1, len(mutation_groups)))
    plotted_groups = 0

    for i, (category, color) in enumerate(zip(mutation_groups.index, colors)):
        if mutation_groups[category] >= 1:  # show all groups
            cat_data = df_survival_variants[df_survival_variants['mutation_category'] == category]
            kmf.fit(cat_data['survival_years'], cat_data['event'],
                   label=f'{category} (n={len(cat_data)})')
            kmf.plot_survival_function(ax=ax, color=color, linewidth=3)
            plotted_groups += 1

    if plotted_groups > 0:
        ax.set_title('survival by mutation type', fontweight='bold', fontsize=14)
        ax.set_xlabel('years from diagnosis')
        ax.set_ylabel('survival probability')
```

```
            ax.grid(True, alpha=0.3)
            ax.legend()
        else:
            ax.text(0.5, 0.5, 'insufficient mutation data', ha='center', va='center', transform=ax.transAxes)
            ax.set_title('survival by mutation type')
    else:
        ax.text(0.5, 0.5, 'no variant class data', ha='center', va='center', transform=ax.transAxes)
        ax.set_title('survival by mutation type')

    plt.tight_layout()
    plt.show()

    # comprehensive summary statistics
    print(f"\ncomprehensive survival analysis:")
    print(f"patients analyzed: {len(df_survival_base)}")
    print(f"overall median survival: {df_survival_base['survival_years'].median():.2f} years")

    if 'gender' in df_survival_base.columns and df_survival_base['gender'].notna().sum() > 0:
        print(f"\nsurvival by all genders:")
        for gender in df_survival_base['gender'].dropna().unique():
            gender_data = df_survival_base[df_survival_base['gender'] == gender]
            print(f"  {gender}: n={len(gender_data)}, median={gender_data['survival_years'].median():.2f} years")

    # show all races
    if 'race' in df_survival_base.columns and df_survival_base['race'].notna().sum() > 0:
        print(f"\nsurvival by all races:")
        for race in df_survival_base['race'].dropna().unique():
            race_data = df_survival_base[df_survival_base['race'] == race]
            print(f"  {race}: n={len(race_data)}, median={race_data['survival_years'].median():.2f} years")
    elif 'ethnicity' in df_survival_base.columns and df_survival_base['ethnicity'].notna().sum() > 0:
        print(f"\nsurvival by all ethnicities:")
        for ethnicity in df_survival_base['ethnicity'].dropna().unique():
            eth_data = df_survival_base[df_survival_base['ethnicity'] == ethnicity]
            print(f"  {ethnicity}: n={len(eth_data)}, median={eth_data['survival_years'].median():.2f} years")

    # show mutation data
    if not df_variants.empty and 'variant_class' in df_variants.columns:
        print(f"\nsurvival by mutation categories:")
        for category in df_survival_variants['mutation_category'].dropna().unique():
            cat_data = df_survival_variants[df_survival_variants['mutation_category'] == category]
            if len(cat_data) > 0:
                print(f"  {category}: n={len(cat_data)}, median={cat_data['survival_years'].median():.2f} years")

else:
    print("no diagnosis-to-death temporal data found for survival analysis")
    print("available temporal observations:")
    if not df_temporal.empty:
        print(df_temporal['obs_type'].value_counts())

# alternative: specimen collection timeline analysis if no survival data
if diagnosis_death.empty and not df_specimens.empty:
    print("\nusing specimen collection as timeline proxy...")

    # this would be specimen collection timing, not survival
    birth_diagnosis = df_temporal[df_temporal['obs_type'] == 'Days Between Birth and Diagnosis']

    if not birth_diagnosis.empty:
        df_timeline = df_main.merge(birth_diagnosis[['patient_id', 'days']], on='patient_id', how='inner')
        df_timeline['age_at_diagnosis'] = df_timeline['days'] / 365.25

        plt.figure(figsize=(12, 6))
```

```
        plt.subplot(1, 2, 1)
        plt.hist(df_timeline['age_at_diagnosis'], bins=20, alpha=0.7, color='skyblue', edgecolor='black')
        plt.title('age at diagnosis distribution')
        plt.xlabel('age (years)')
        plt.ylabel('frequency')
        plt.grid(True, alpha=0.3)

        plt.subplot(1, 2, 2)
        if 'gender' in df_timeline.columns and df_timeline['gender'].notna().sum() > 5:
            for gender in df_timeline['gender'].dropna().unique():
                gender_data = df_timeline[df_timeline['gender'] == gender]
                plt.hist(gender_data['age_at_diagnosis'], alpha=0.6,
                        label=f'{gender} (n={len(gender_data)})', bins=15)
            plt.title('age at diagnosis by gender')
            plt.xlabel('age (years)')
            plt.ylabel('frequency')
            plt.legend()
            plt.grid(True, alpha=0.3)

        plt.tight_layout()
        plt.show()

        print(f"\nage at diagnosis summary:")
        print(f"patients: {len(df_timeline)}")
        print(f"mean age: {df_timeline['age_at_diagnosis'].mean():.1f} years")
        print(f"median age: {df_timeline['age_at_diagnosis'].median():.1f} years")
        print(f"age range: {df_timeline['age_at_diagnosis'].min():.1f} - {df_timeline['age_at_diagnosis'].max():.1f} years")
```

```
extracting temporal data for kaplan-meier analysis...
extracted 47 temporal observations
found 16 diagnosis-to-death observations
survival analysis dataset: 33 patients
mean survival: 1.6 years
median survival: 1.5 years

debugging variant class data:
survival patients: 33
variant classes in all data: {'SNV': 5931, 'deletion': 1063, 'insertion': 213, 'indel': 3}
variant classes in survival cohort: {'SNV': 1144, 'deletion': 72, 'insertion': 22}
mutation categories in survival cohort: {'SNV': 1144, 'indel/deletion': 94}
```

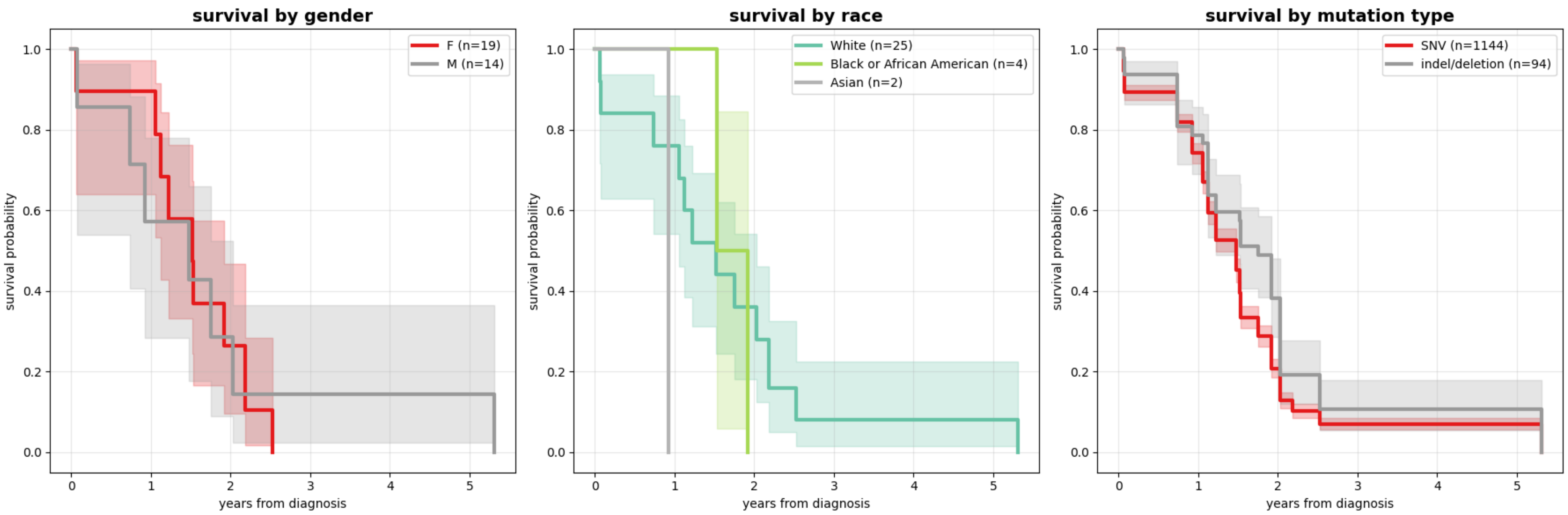


```
comprehensive survival analysis:
patients analyzed: 33
overall median survival: 1.52 years

survival by all genders:
  F: n=19, median=1.52 years
  M: n=14, median=1.48 years

survival by all races:
  White: n=25, median=1.52 years
  Black or African American: n=4, median=1.73 years
  Asian: n=2, median=0.93 years

survival by mutation categories:
  SNV: n=1144, median=1.48 years
  indel/deletion: n=94, median=1.75 years
```

```
import pandas as pd
import numpy as np
import matplotlib.pyplot as plt
import matplotlib.patches as patches
from matplotlib import gridspec
import seaborn as sns
from scipy.stats import fisher_exact
from collections import defaultdict
import warnings
warnings.filterwarnings('ignore')

# hardcoded pathway definitions for cholangiocarcinoma
CHOLANGIOCARCINOMA_PATHWAYS = {
    "MAPK signaling": ["KRAS", "BRAF", "NRAS", "MAP2K1", "MAP2K4", "MAPK1", "MAPK3"],
    "PI3K/AKT pathway": ["PIK3CA", "PTEN", "AKT1", "MTOR", "PIK3R1"],
    "p53 pathway": ["TP53", "ATM", "CHEK2", "MDM2", "CDKN1A", "CDKN2A"],
```

```python
    "Chromatin remodeling": ["ARID1A", "ARID1B", "BAP1", "SMARCA4", "SMARCB1", "ARID2"],
    "DNA repair": ["BRCA1", "BRCA2", "ATR", "PALB2", "BRIP1", "RAD51C"],
    "Cell cycle": ["CDKN2A", "CCND1", "RB1", "E2F1", "CDK4", "CDK6"],
    "Wnt signaling": ["APC", "CTNNB1", "AXIN1", "GSK3B", "AXIN2", "TCF7L2"],
    "Metabolism": ["IDH1", "IDH2", "MSMO1", "TYMP"],
    "Other targets": ["PCLO", "PLEC", "KDR", "DMD", "BCAN", "BCOR", "PSCA", "TTN"]
}

MUTATION_COLORS = {
    "SNV": "#4CAF50",
    "deletion": "#F44336",
    "insertion": "#2196F3",
    "indel": "#FF9800",
    "multiple": "#424242"
}

def pathway_enrichment_analysis(mutated_genes):
    """perform fisher exact test for pathway enrichment"""
    enriched_pathways = []

    for pathway_name, pathway_genes in CHOLANGIOCARCINOMA_PATHWAYS.items():
        mutated_in_pathway = len(set(mutated_genes).intersection(set(pathway_genes)))
        if mutated_in_pathway > 0:
            enriched_pathways.append({
                'pathway': pathway_name,
                'mutated_genes': mutated_in_pathway,
                'total_genes': len(pathway_genes),
                'genes': [g for g in pathway_genes if g in mutated_genes]
            })

    return sorted(enriched_pathways, key=lambda x: x['mutated_genes'], reverse=True)

def get_pathway_ordered_genes(df_mutations):
    """order genes by pathway groups and mutation frequency"""
    # get all mutated genes
    gene_counts = df_mutations['gene'].value_counts()
    mutated_genes = gene_counts.index.tolist()

    # perform enrichment
    enriched_pathways = pathway_enrichment_analysis(mutated_genes)

    # organize genes by pathway
    ordered_genes = []
    used_genes = set()

    # add genes from enriched pathways in order
    for pathway_info in enriched_pathways:
        pathway_genes = pathway_info['genes']
        # sort by frequency within pathway
        pathway_genes_sorted = sorted(pathway_genes,
                                      key=lambda g: gene_counts.get(g, 0), reverse=True)
        for gene in pathway_genes_sorted:
            if gene not in used_genes:
                ordered_genes.append(gene)
                used_genes.add(gene)

    # add remaining genes not in pathways
    for gene in mutated_genes:
        if gene not in used_genes:
            ordered_genes.append(gene)
```

```python
    return ordered_genes[:25], enriched_pathways  # limit to top 25 genes

def create_waterfall_oncoprint(df_variants):
    """create waterfall-style oncoprint"""

    # extract mutations
    mutations = []
    for _, row in df_variants.iterrows():
        if pd.notna(row.get('hugo_symbol')) and pd.notna(row.get('patient_id')):
            mutations.append({
                'patient_id': row['patient_id'],
                'gene': row['hugo_symbol'],
                'variant_class': row.get('variant_class', 'SNV')
            })

    df_mutations = pd.DataFrame(mutations)
    if df_mutations.empty:
        return None

    # get ordered genes and pathways
    ordered_genes, pathway_info = get_pathway_ordered_genes(df_mutations)

    # order patients by total mutation count (waterfall effect)
    patient_mutation_counts = df_mutations.groupby('patient_id').size().sort_values(ascending=False)
    ordered_patients = patient_mutation_counts.head(50).index.tolist()  # top 50 patients

    # create figure with proper layout
    fig = plt.figure(figsize=(18, 12))
    gs = gridspec.GridSpec(3, 1, height_ratios=[1.5, 8, 0.8], hspace=0.1)

    # mutation burden barplot (top)
    ax_burden = fig.add_subplot(gs[0, 0])
    burden_values = [patient_mutation_counts.get(p, 0) for p in ordered_patients]

    bars = ax_burden.bar(range(len(ordered_patients)), burden_values,
                        color='#2E86AB', alpha=0.8, width=0.9)

    # add median line
    median_val = np.median(burden_values)
    ax_burden.axhline(y=median_val, color='#F24236', linestyle='--', linewidth=2)

    ax_burden.set_xlim(-0.5, len(ordered_patients) - 0.5)
    ax_burden.set_ylabel('mutations per patient', fontweight='bold')
    ax_burden.set_xticks([])
    ax_burden.spines['top'].set_visible(False)
    ax_burden.spines['right'].set_visible(False)
    ax_burden.grid(axis='y', alpha=0.3)

    # main oncoprint grid (center)
    ax_onco = fig.add_subplot(gs[1, 0])

    # create mutation lookup
    mutation_dict = {}
    for _, row in df_mutations.iterrows():
        key = (row['gene'], row['patient_id'])
        if key not in mutation_dict:
            mutation_dict[key] = []
        mutation_dict[key].append(row['variant_class'])

    # plot mutation grid
    for i, gene in enumerate(ordered_genes):
```

```python
    y_pos = len(ordered_genes) - i - 1

    for j, patient in enumerate(ordered_patients):
        rect = patches.Rectangle(
            (j, y_pos), 1, 1,
            linewidth=0.3, edgecolor='#E0E0E0',
            facecolor='white'
        )
        ax_onco.add_patch(rect)

        key = (gene, patient)
        if key in mutation_dict:
            mutations_list = mutation_dict[key]
            if len(mutations_list) == 1:
                color = MUTATION_COLORS.get(mutations_list[0], '#666666')
            else:
                color = MUTATION_COLORS['multiple']
            rect.set_facecolor(color)

# add gene labels with frequencies
for i, gene in enumerate(ordered_genes):
    y_pos = len(ordered_genes) - i - 1 + 0.5

    # gene name
    ax_onco.text(-0.5, y_pos, gene, ha='right', va='center',
                fontweight='bold', fontsize=10)

    # frequency
    gene_patients = set(df_mutations[df_mutations['gene'] == gene]['patient_id'])
    freq = (len(gene_patients.intersection(set(ordered_patients))) / len(ordered_patients)) * 100
    ax_onco.text(-0.1, y_pos, f'{freq:.0f}%', ha='right', va='center',
                fontsize=9, color='#666666')

# add pathway groupings on the right
y_position = len(ordered_genes)
pathway_colors = ['#1f77b4', '#ff7f0e', '#2ca02c', '#d62728', '#9467bd',
                 '#8c564b', '#e377c2', '#7f7f7f', '#bcbd22']
color_idx = 0

for pathway_data in pathway_info:
    pathway_name = pathway_data['pathway']
    pathway_genes = [g for g in pathway_data['genes'] if g in ordered_genes]

    if pathway_genes:
        n_genes = len(pathway_genes)
        y_position -= n_genes

        # draw bracket
        bracket_x = len(ordered_patients) + 1
        bracket_color = pathway_colors[color_idx % len(pathway_colors)]

        # vertical line
        ax_onco.plot([bracket_x, bracket_x], [y_position, y_position + n_genes],
                    color=bracket_color, linewidth=3)

        # top and bottom horizontals
        ax_onco.plot([bracket_x, bracket_x + 0.3], [y_position, y_position],
                    color=bracket_color, linewidth=2)
        ax_onco.plot([bracket_x, bracket_x + 0.3], [y_position + n_genes, y_position + n_genes],
                    color=bracket_color, linewidth=2)
```

```
            # pathway label
            ax_onco.text(bracket_x + 0.5, y_position + n_genes/2, pathway_name,
                        va='center', ha='left', fontsize=10, fontweight='bold',
                        color=bracket_color, rotation=0)

            color_idx += 1

    ax_onco.set_xlim(-1.2, len(ordered_patients) + 8)
    ax_onco.set_ylim(0, len(ordered_genes))
    ax_onco.set_xticks([])
    ax_onco.set_yticks([])

    # remove all spines
    for spine in ax_onco.spines.values():
        spine.set_visible(False)

    # legend (bottom)
    ax_legend = fig.add_subplot(gs[2, 0])
    legend_elements = []

    # only show mutation types that exist in the data
    existing_types = set()
    for mutations_list in mutation_dict.values():
        existing_types.update(mutations_list)

    for variant_type in ['SNV', 'deletion', 'insertion', 'indel', 'multiple']:
        if variant_type in existing_types or variant_type == 'multiple':
            color = MUTATION_COLORS[variant_type]
            legend_elements.append(patches.Patch(color=color, label=variant_type))

    ax_legend.legend(handles=legend_elements, loc='center', ncol=len(legend_elements),
                    frameon=False, fontsize=12)
    ax_legend.axis('off')

    plt.suptitle('cholangiocarcinoma mutation landscape',
                fontsize=16, fontweight='bold', y=0.98)

    return fig

# create the waterfall oncoprint
if 'df_variants' in globals() and not df_variants.empty:
    fig = create_waterfall_oncoprint(df_variants)
    if fig:
        plt.tight_layout()
        plt.show()
```

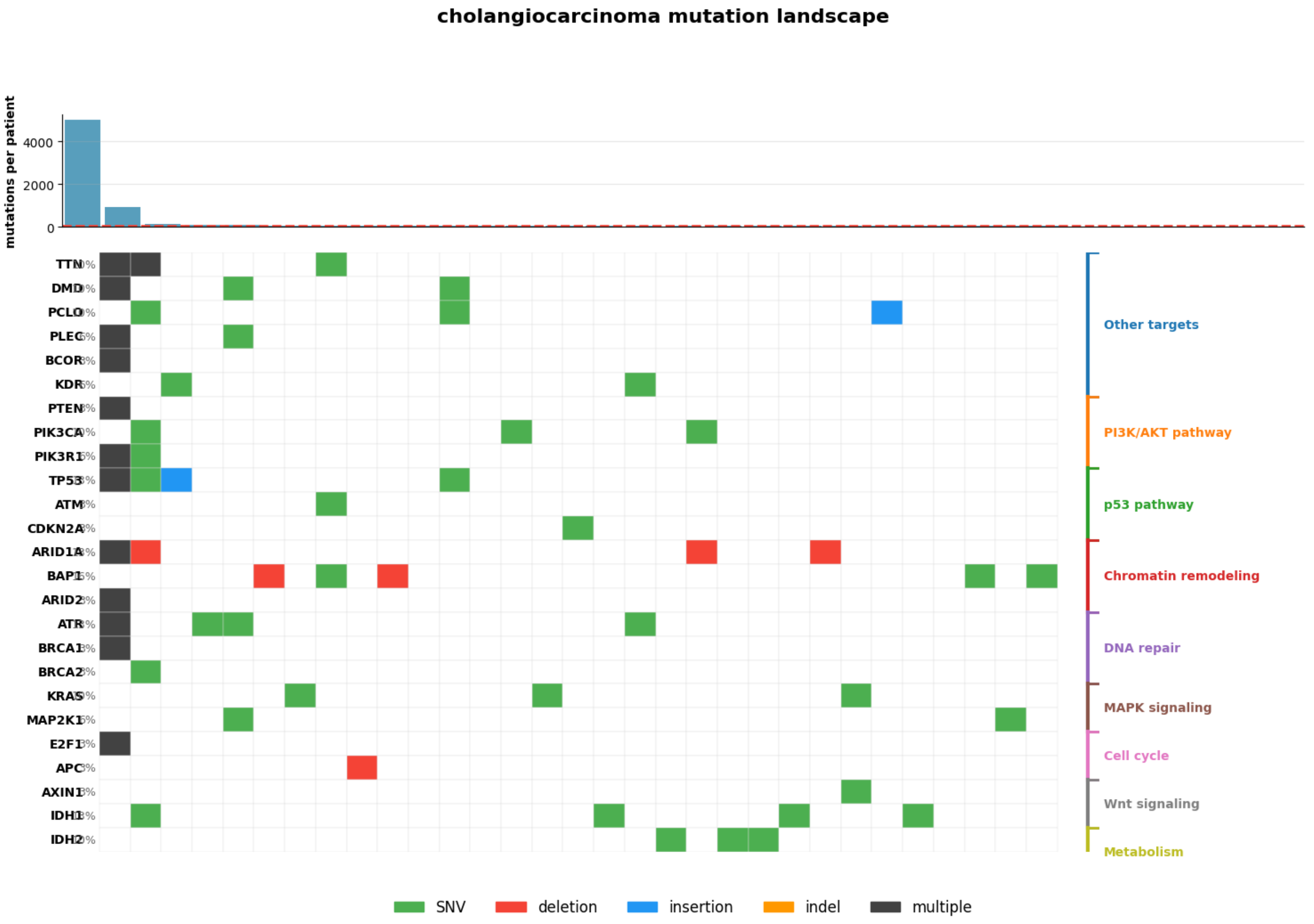
cholangiocarcinoma mutation landscape
mutations per patient
4000
2000
0
TTN
DMD
PCLO
PLEC
BCOR
KDR
PTEN
PIK3CA
PIK3R1
TP53
ATM
CDKN2A
ARID1A
BAP1
ARID2
ATR
BRCA1
BRCA2
KRAS
MAP2K1
E2F1
APC
AXIN1
IDH1
IDH2
Other targets
PI3K/AKT pathway
p53 pathway
Chromatin remodeling
DNA repair
MAPK signaling
Cell cycle
Wnt signaling
Metabolism
SNV
deletion
insertion
indel
multiple
